# A vast space of compact strategies for highly efficient decisions

**DOI:** 10.1101/2022.08.10.503471

**Authors:** Tzuhsuan Ma, Ann M Hermundstad

## Abstract

When foraging in dynamic and uncertain environments, animals can benefit from basing their decisions on smart inferences about hidden properties of the world. Typical theoretical approaches to understand the strategies that animals use in such settings combine Bayesian inference and value iteration to derive optimal behavioral policies that maximize total reward given changing beliefs about the environment. However, specifying these beliefs requires infinite numerical precision; with limited resources, this problem can no longer be separated into optimizing inference and optimizing action selections. To understand the space of behavioral policies in this constrained setting, we enumerate and evaluate all possible behavioral programs that can be constructed from just a handful of states. We show that only a small fraction of the top-performing programs can be constructed by approximating Bayesian inference; the remaining programs are structurally or even functionally distinct from Bayesian. To assess structural and functional relationships among all programs, we developed novel tree embedding algorithms; these embeddings, which are capable of extracting different relational structures within the program space, reveal that nearly all good programs are closely connected through single algorithmic “mutations”. We demonstrate how one can use such relational structures to efficiently search for good solutions via an evolutionary algorithm. Moreover, these embeddings reveal that the diversity of non-Bayesian behaviors originates from a handful of key mutations that broaden the functional repertoire within the space of good programs. The fact that this diversity of behaviors does not significantly compromise performance suggests a novel approach for studying how these strategies generalize across tasks.

## INTRODUCTION

To thrive in a changing world, animals need to gather information about hidden states of the world to guide decisions and plan future actions. This is true of many tasks, such as localizing a food source from noisy measurements [1, 2, 3], predicting the location of a moving target during pursuit [4, 5], or planning efficient routes to traverse through a set of subgoals [6, 7]. In these and other domains, there are in principle many possible ways for an animal to leverage knowledge about its surroundings in order to make inferences and guide future actions. These different strategies can vary in their complexity (e.g. updating an internal estimate of a food source using finely-encoded sequences of odor samples versus coarsely-averaged odor concentrations [3]), and in their performance (e.g. slowly versus quickly localizing a food source based on this internal estimate [2]). A common theoretical approach is to identify the optimal strategy for solving a particular task or set of tasks, given some assumptions about the prior knowledge that an animal maintains about its surroundings, or given assumptions about the algorithmic or circuit architecture that implements this strategy [8, 9, 10]. This optimal strategy can then be dissected to understand the architectural features that enable good performance, or it can be used as a benchmark to compare to neural and behavioral data. However, by focusing on a single best strategy, this type of approach does not provide a natural way of understanding the many possible strategies that could be used to achieve good enough, if not optimal, performance. Here, we develop an approach for defining and exploring such a strategy *space* in terms of different resource limitations that constrain the complexity and performance of individual strategies.

To illustrate this space, we focus on a well-studied task in which an animal, or agent, gathers rewards from two ports whose reward probabilities change dynamically over time according to a hidden world state [11, 12, 13, 14, 15, 16]. To improve performance on this task, the agent can leverage knowledge about the volatility of the environment and the probability of rewards at each port in order to update an internal estimate of the world state based on the outcomes of its past actions. This inference can then be used to guide future actions that will maximize the number of rewards that the animal can collect from the ports [17]. Recent studies have suggested that animals leverage such inferences to guide decisions in the face of uncertainty [18, 19, 20]. In the absence of such inferences, other studies have shown how animals could directly estimate the value of individual ports, and use these estimates to guide behavior [21, 14, 15]. These studies focus on a small number of candidate strategies that can optimally guide behavior based on precise internal estimates of world variables.

Here, we consider how behavior in this task could be guided by a diversity of different strategies that are limited in their resources, and that differ in their structure, behavior, and performance. We impose this resource limitation by formulating an agent that uses a limited number of internal states to guide behavior; this limitation, in turn, imposes a constraint on how precisely the agent can maintain and update internal estimates about the world. In contrast to the single optimal agent (e.g. Bayesian agent) that can only be derived in the absence of resource limitations, there are many possible ways to build a constrained agent, each of which can be expressed as a small program that guides future behavior based on the consequences of past actions [22]. We enumerate the set of all unique small programs that can be realized up to a certain number of internal states, and we compare these diverse programs to those ones that can be obtained directly from naively discretizing an optimal Bayesian agent. This enables us to study relationships between the internal wiring, behavior, and performance of different programs, and identify structural and functional features that enable better-than-chance performance.

We show that the set of good programs that achieve high performance occupy a small fraction of the full program space, but are nevertheless diverse and exhibit structural and functional properties that differ from discretized Bayesian agents. We find that good programs are connected to one another through single mutations that minimally alter the internal wiring of a program. This finding enables us to construct an evolutionary algorithm that can efficiently search for good programs within the full program space. To understand how these good programs are related to each other, we use both structure (defined by the wiring of individual programs) and function (defined by sequences of actions produced in response to different outcomes) to link individual programs within a hierarchical tree; by designing this tree to prioritize structural and functional similarities between programs, we can identify those key mutations in the wiring of individual programs that lead to significant changes in behavior, and distinguish these from sloppy mutations that do not significantly impact behavior. Together, this work provides a framework for enumerating and exploring spaces of possible strategies that could be used to solve the same behavioral task, and it reveals that the space of “good enough” solutions that emerge in the presence of resource limitations can be diverse and qualitatively different from the optimal strategies derived in the absence of such limitations.

## RESULTS

We consider a general scenario in which an agent uses incoming observations to make inferences about unobservable world states, and uses those inferences to guide actions that lead to rewards (Figure 1). In the absence of resource constraints, the optimal strategy for maximizing rewards can be derived via two separable approaches: (i) Bayesian inference can be used to derive the ideal observer that constructs and updates an internal belief about the set of world states (see SI Section 5), and (ii) value iteration can be used to derive the optimal policy that maximizes rewards (see SI Section 5.2), given the observer’s belief (Figure 1a). However, this optimal solution requires infinite resources in order to specify an internal belief with arbitrarily fine precision; given limited resources to specify this belief, the problem is no longer separable, and it is not clear how to efficiently search for or construct the optimal behavioral strategy (Figure 1b).

**Figure 1:**
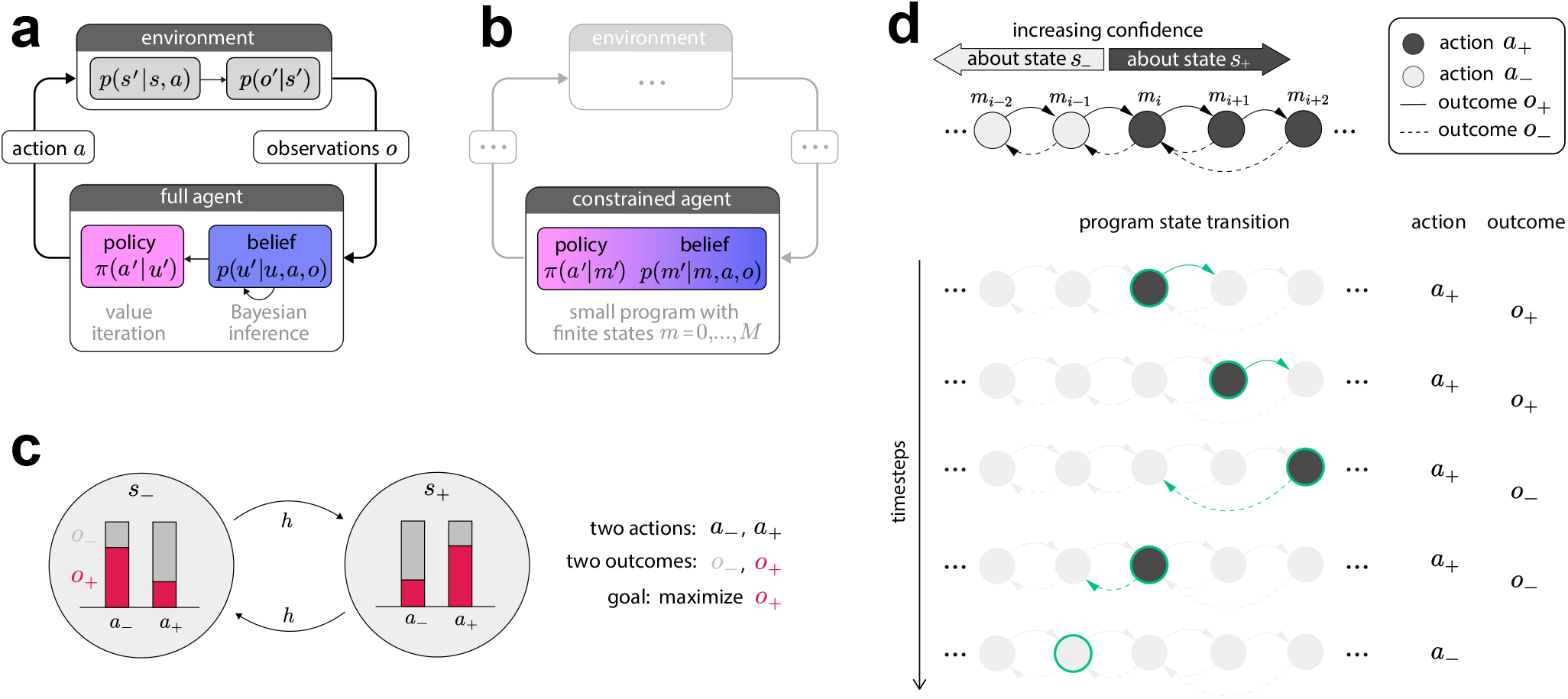
Problem setup: specifying a resource-constrained agent as a finite-state behavioral program. **(a)** An optimal (unconstrained) agent can be specified via two separate modules: an internal belief derived via Bayesian inference (purple), and a policy optimized via value iteration in belief space (pink). This separation enables efficient optimization of the behavioral policy, conditioned on the internal belief. **(b)** To optimize a constrained agent with limited resources, we must simultaneously specify the belief update *p*(*m*′|*m, a, o*) and the policy *π*(*a*′|*m*′). This lack of separability prohibits an efficient search for the optimal behavioral policy. **(c)** We consider a non-stationary two-armed bandit task in which an agent can gather rewards from two ports whose reward probabilities can change over time according to an unobservable world state. In order to maximize reward, the agent must continually update its belief about this world state in order to select the appropriate action to gather reward. The task is specified by the hazard rate *h*, the reward contrast Δ*p*, and baseline reward rate 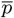. **(d)** We formulate a constrained agent as a small program that consists of a finite set of states that can be represented as nodes in a Markov chain. A deterministic policy for such a program can then be specified by labelling these nodes with actions *a*_+_ or *a*_−_; a set of deterministic state transitions is specified with outcome-dependent edges between nodes (dashed and solid arrows). The example program illustrates how the machinery of a Bayesian belief update can be qualitatively captured by such state transitions, by increasing confidence upon a consistent action-outcome pair, or by decreasing confidence upon a conflicting action-outcome pair.

To circumvent this, we enumerate all possible ways to construct this internal belief under the assumption that it can be represented by a deterministic Markov chain with a finite number of states. We refer to this Markov chain as a small program; this small program uses its internal states to select one of a discrete set of actions conditioned on past observations. To illustrate this enumeration and study the programs derived from it, we study a non-stationary two-armed bandit task in which an agent must continually infer a changing world state to gather rewards (Figure 1c). On each timestep, the agent can select one of two actions, *a*_+_ and *a*_−_, and can receive one of two outcomes, *o*_+_(rewarding) and *o*_−_ (unrewarding). A binary world state *s* determines the probability that a given action will produce a given observation; we assume that *s* switches between two states, *s*_+_ and *s*_−_, at a fixed probability *h* per timestep with either a high reward probability *p*_high_ ≡ *p*(*o*_+_|*s*_±_, *a*_±_) when *a* and *s* are aligned or a low reward probability *p*_low_ ≡ *p*(*o*_+_|*s*_±_, *a*_∓_) when they are misaligned. Together, this task can be fully specified by the baseline reward rate of the two arms 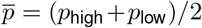, the reward contrast between the two arms Δ*p* = *p*_high_ – *p*_low_, and the hazard rate *h* (see SI Section 5).

If a world state is observable, the optimal strategy would be to choose the action whose sign is aligned with the current world state. However, without knowing the true value of the world state, the optimal achievable strategy (in the absence of resource constraints) is to combine incoming observations with existing task knowledge in order to update an internal belief about the world state, and select the action whose sign is aligned with that belief. In the presence of resource constraints, a small program can instead use a fixed number of states to select an action based on incoming observations (Figure 1d). One of the simplest and best-known small program implements a win-stay, lose-go (WSLG) strategy (Figure 2c); this program will continue taking the same action so long as the action yields a reward; if the action does not yield a reward, the program will take the alternative action. Such small programs can take on many different forms, depending on the number of states, and precise transition structure between states.

**Figure 2:**
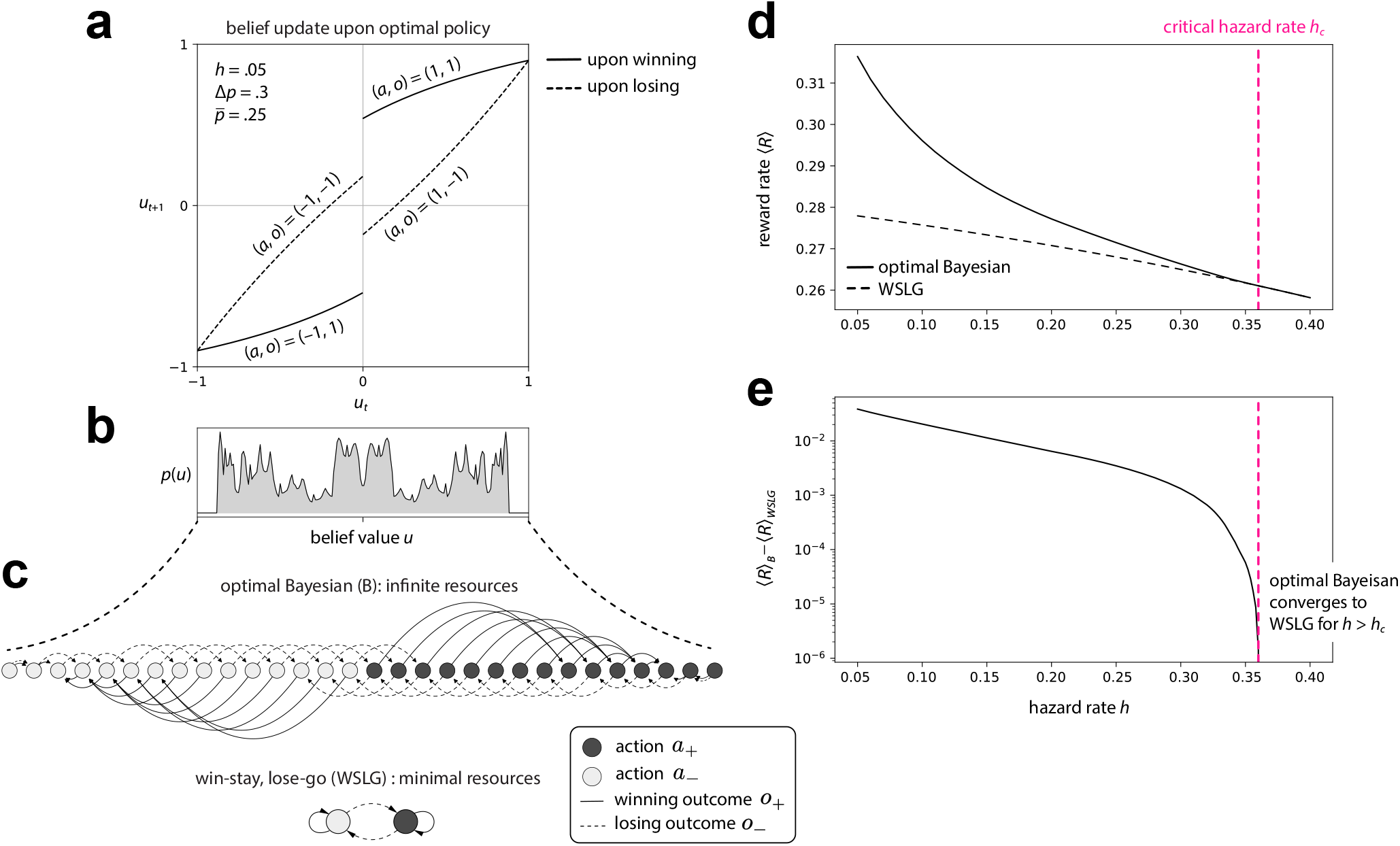
The space of good behavioral programs is bounded by both the most and the least resource-constrained agent. **(a)** Illustration of a single step of the Bayesian belief update under the optimal policy; the belief value is defined as the difference in the posterior belief about each of the two world states—i.e., *u* ≡ *p*(*s* = *s*_+_ | ⋯) – *p*(*s* = *s*_−_ | ⋯). This update rule can be qualitatively understood as monotonically increasing/decreasing the belief value *u* upon winning/losing. **(b)** The infinite-horizon distribution of belief values is obtained via Belief Distribution Propagation (See SI Section 8.1). This distribution can then be used to compute the reward rate in panels (d,e). **(c)** An optimal Bayesian agent can be approximated by a small program with a large but finite number of states (shown here with 30 states). In contrast, the best-performing agent with the smallest number of states is the win-stay, lose-go (WSLG) program. **(d-e)** The performance gap between WSLG and the optimal Bayesian agent decreases with an increasing hazard rate. This gap decreases to zero at a critical hazard rate *h_c_* = .36 discussed in (see SI Section 6). The large performance gap at lower hazard rates suggests the existence of many good programs that could achieve intermediate performance; we use this observation to select a hazard rate of *h* = 0.05 for evaluating and studying the performance of these programs.

### Different task parameters allow for many good behavioral programs

By adjusting the number of internal states that we use to construct small programs, we can generate a variety of programs that vary in performance. At one extreme, as the number of states becomes infinite, we can exactly reproduce the optimal Bayesian agent (B) that uses a precise internal belief to maximize performance (Figure 2a-c). At the other extreme, a small program such as WSLG achieves much lower, albeit better-than-chance, performance (Figure 2c, lower); note that an even smaller, memoryless program can be constructed to implement a biased random walk with no internal state transitions, and achieves a mere chance reward rate 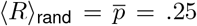. The WSLG program therefore uses the minimum resources needed to improve upon this baseline performance. We use these two extreme programs (B vs WSLG) to bound the space of “good programs”; the performance gap between these two extremes varies as a function of task parameters, and determines the number of good programs that can achieve intermediate performance (See Figure S7 in SI). This gap is largest in the limit that the environment is stable (Figure 2d). As the environment becomes more volatile, the performance of both B and WSLG decrease, and the gap between them shrinks to zero (Figure 2e). Note that while these performance curves are smooth, the Bayesian agent exhibits abrupt transitions in its behavior, which give rise to discontinuities such as the sudden collapse of the good program space around *h* = .36; see Section 6 in SI for details and discussion.

### A tree embedding reveals the relational structure of program space

We use the performance gap between B and WSLG discussed above to select the task parameters that yield a large number of good programs 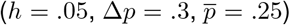. For this set of task parameters, we enumerate and evaluate all unique programs that have up to five internal states (Figure 3a-c shows examples of these programs; see SI Section 7 for details). We use two different types of enumeration: (i) we translate an optimal Bayesian agent into a small program by uniformly discretizing its belief space, and enumerate over possible discretizations (Figure 3b), and (ii) we enumerate all possible programs with up to five states (Figure 3a-e; note that this set is complete and includes the set of discretized Bayesian programs). The first enumeration produces a set of programs that exhibit the structural features of Bayesian inference; these programs have two different integration directions that can be interpreted as increasing the certainty that the environment is in one state or the other (Figure 3b). We enumerate 88 such “structurally-Bayesian” programs; of these, 58 (65.9%) exhibit good performance. The second enumeration yields a wide diversity of different program structures that, in general, do not exhibit the structural features of the Bayesian programs (Figure 3c). There are 268,536 such programs; of these, 4230 (1.6%) exhibit good performance (Figure 3d). Together, this highlights that the space of small programs is vast (Figure 3e); a mere 1.6% of the space amounts to a staggering 4230 good programs. Among these good programs, only 58 (1.4%) can be efficiently found by naively discretizing an optimal Bayesian agent.

**Figure 3:**
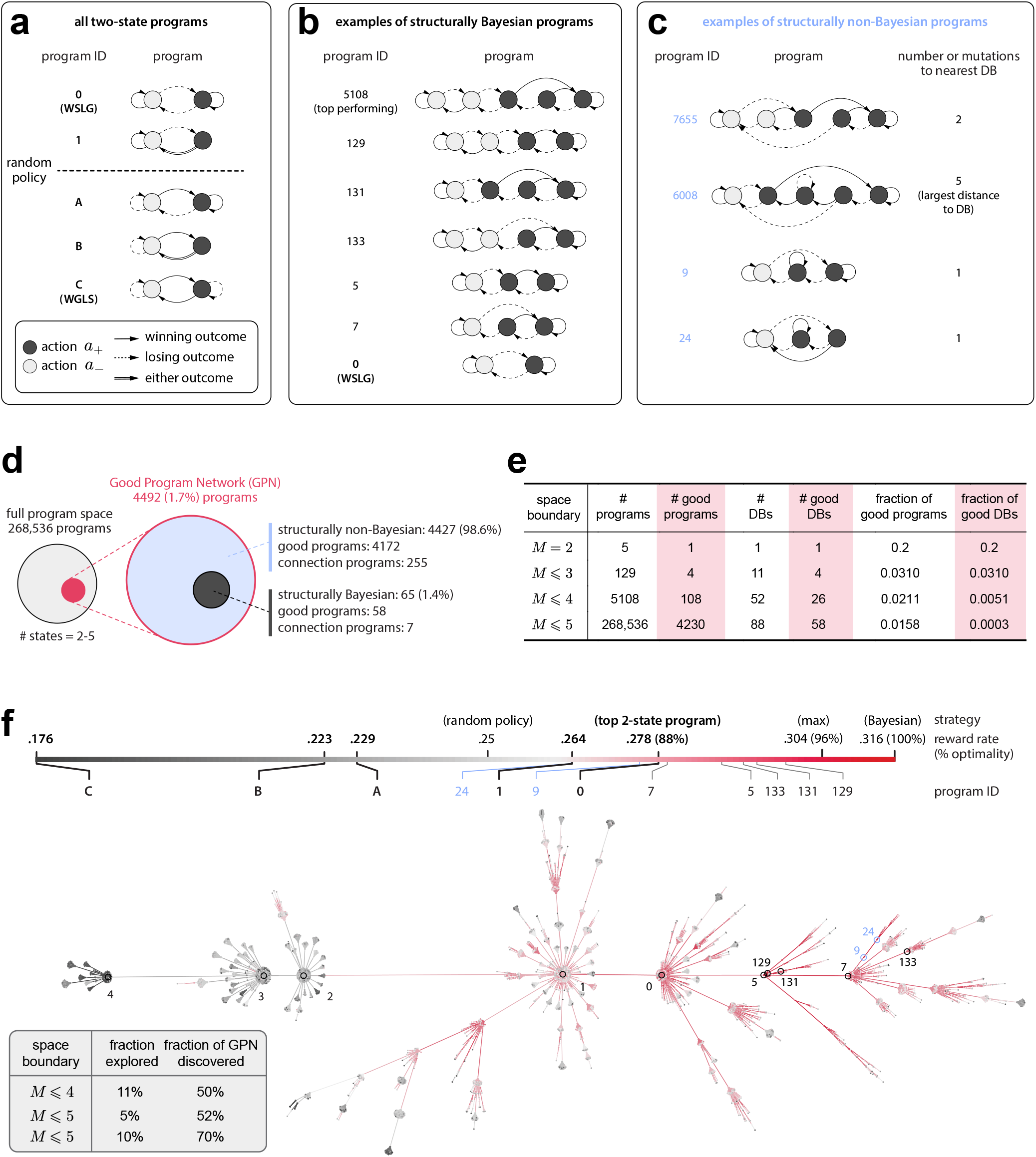
The space of behavioral programs is highly structured. **(a)** Enumeration reveals 5 unique two-state programs; the top-performing program is the WSLG program. Sorting these programs by performance reveals that each neighboring pair of programs differ by a single mutation (a relabeling of a single node or a relocation of a single arrow). **(b)** Example programs that are structurally similar to the optimal Bayesian agent, sorted by performance (highest at top); the smallest of these is the WSLG program. These “structurally Bayesian” programs can improve their performance by increasing their resource-use (i.e., increasing the number of nodes) while maintaining two distinct directions of integration within the nodes corresponding to each of the two actions. **(c)** Example programs that are structurally non-Bayesian, sorted by performance (highest at top). **(d)** Although good programs comprise a small fraction of the full program space, they nevertheless amount to a numerous 4230 programs in total. Among these good programs, only a small fraction can be found by discretizing an optimal Bayesian agent. Note that, in GPN, we keep a small number of lower-performing “connection” programs that glue all disjoint branches into a single subtree (see Section 9.6 for details). **(e)** As the maximal number of resources *M* increases, the fraction of structurally Bayesian programs becomes exponentially small. **(f)** We perform a tree embedding to extract the minimal relational structure among all programs, shown here for all programs up to *M* = 4 states (see SI Section 9.5 for a full visualization up to *M* = 5). This embedding, visualized with Gephi [23] using the Yifan Hu layout [24], reveals that high-performing programs are much more likely to connect with one another than with low-performing programs. The fact that the space is highly structured can be exploited to design a more efficient search via an evolutionary algorithm that can discover good programs with high probability from searching a small fraction of the full space (inset).

To understand whether these good programs exhibit similarities in their structure, we examined how structural differences impact performance. When we rank-ordered the set of 5 two-state programs by performance (Figure 3a), we found that neighboring programs are separated by a single mutation (defined by relocating one arrow or relabeling one node). We used this observation to design a tree embedding algorithm that preferentially assigns a small and high-performance parent program to each children program, provided that they are within a single mutation of one another (Figure 3f; Figure S10 shows a tree-embedding performed with randomized performance). With this embedding, we find that nearly all good programs are closely connected to form a singular subtree, which we term the “good program network” (GPN). We then used the structure in the GPN to design an efficient evolutionary search algorithm that identified 70% of all good programs by exploring only 10% of the full program space (inset of Figure 3f).

### A functional tree embedding reveals behaviorally-distinct regions of program space

The previous tree embedding reveals relationships between structurally distinct programs that exhibit a similar performance. To better understand how program structure impacts performance, we evaluated the sets of behavioral sequences that enable different programs to achieve high performance (Figure 4a). These behavioral sequences can be defined in terms of outcome-action pairs; the operations ”win-stay” (i.e., given that I observed a positive outcome, repeat the same action) and “lose-go” (i.e., given that I observed a negative outcome, do not repeat the same action) are examples of such outcome-action pairs. For each program, we enumerated all sequences produced by the program up to maximum sequence length of 10, which produced large ensembles of sequences that define the behaviors of each program.

**Figure 4:**
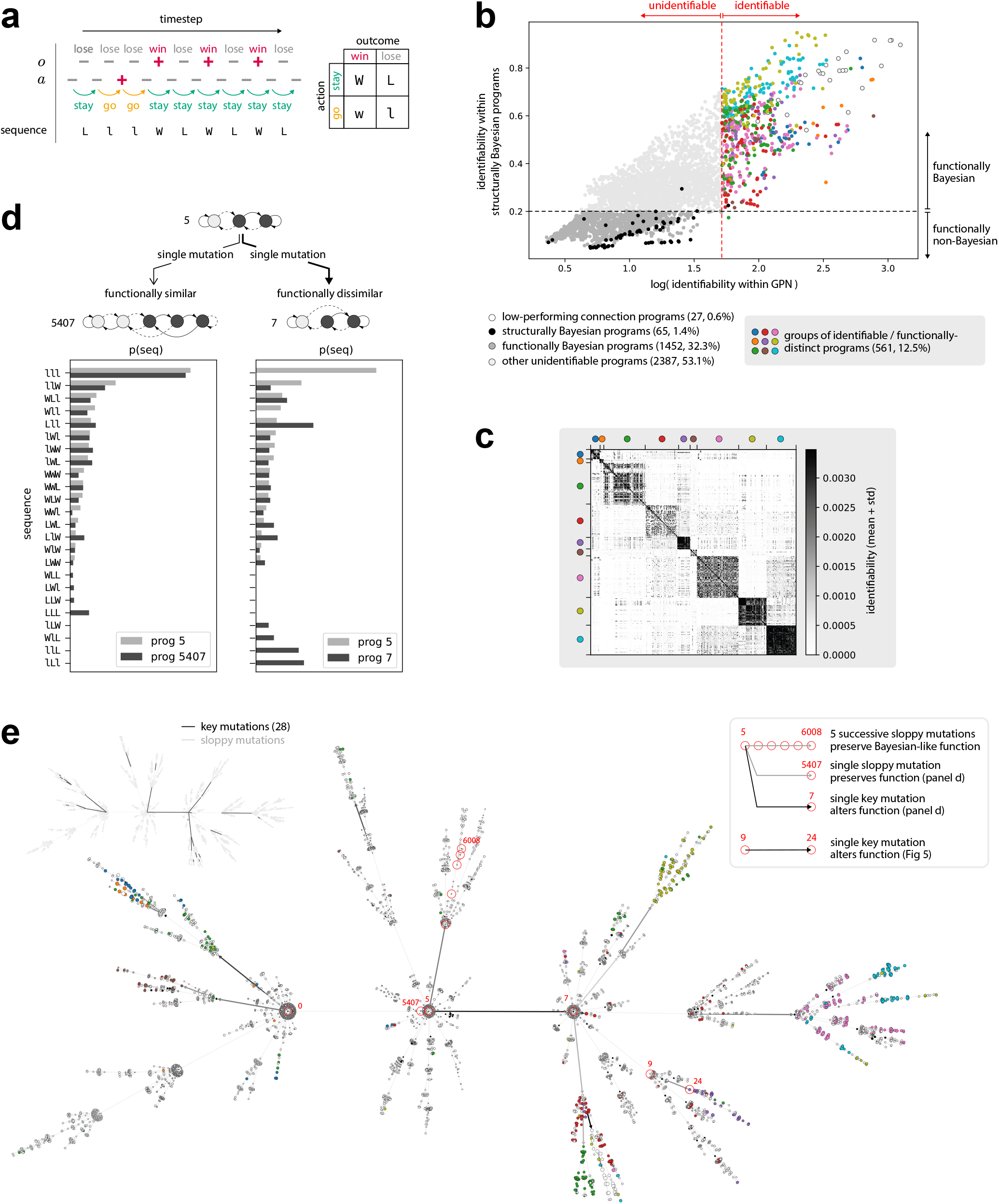
Behaviorally distinct programs emerge from a handful of key mutations. **(a)** From the symmetry of our task, an outcome-action sequence can be expressed as a combination of outcomes (“win” vs “lose”) and actions (“stay”, i.e., repeat the previous action, vs “go”, i.e., select a different action) **(b)** All programs within the good program network (GPN) are labelled based on their functional properties. Specifically, we measure how much each program can be identified among all other programs in the GPN, and how much it can be identified among all structurally Bayesian programs; the latter measures the similarity of a program’s behavior to that of a Bayesian agent. The threshold for a program to be labeled “identifiable” (color dots) is defined such that the fewest number of identifiable programs can cover half of the total identifiability within the entire GPN. The threshold for a program to be labeled “functionally Bayesian” (darker gray dots) is defined such that all functionally Bayesian programs do not exceed the maximum identifiability within the set of structurally Bayesian programs (note that we excluding two outliers in setting this threshold). Surprisingly, we observe a a clear separation between the set of identifiable programs and the set of functionally Bayesian programs; all (but one outlier) identifiable programs are thus qualitatively distinct from the optimal Bayesian agent. **(c)** A clustering using `community_louvain` [26] on all identifiable programs reveals several functional groups that are approximately orthogonal to one another, highlighting that the GPN as a whole is functionally diverse. **(d-e)** A functional tree embedding (fTE) of the GPN reveals that the sets of functionally distinct programs emerge from a small number of key mutations. **(d)** A single mutation can cause either a small or a large change in the distribution of behavioral sequences. We define a key mutation as one that generates a large functional change, whereas a sloppy mutation is one that generates a minimal functional change. **(e)** fTE reveals a handful (28 out of 4492) of key mutations in an otherwise functionally-smooth GPN. By traversing from the root node (WSLG program) towards the leaf nodes, an accumulation of a few key mutations along each lineage is sufficient to generate functionally distinct ensembles of programs (colored nodes near leaf nodes). Note that a parent program can also undergo many mutations without significantly changing in its functions (e.g., program 5 accumulates several mutations to become the functionally-similar program 6008, as highlighted in the insert).

We first assessed how easily we could identify individual programs based on this ensemble of sequences (see SI Section 10.1 for details; see also [25]); we measured this identifiability with respect to all programs within the GPN, and separately with respect to the subset of discretized Bayesian programs. We found that a large subset (32.3%) of programs were not easily distinguished from the set of Bayesian programs (Figure 4b, dark gray points); we refer to this subset as “functionally Bayesian”. Moreover, these functionally-Bayesian programs were also easily confused with the ensemble of other programs within the GPN. In contrast, a distinct subset (comprising 12.5%) of programs were easily distinguished from all programs within the GPN and from all functionally-Bayesian programs (colored points in Figure 4b), highlighting the fact that many programs exhibit non-Bayesian behavior that makes them identifiable from the bulk of other programs. To understand the nature of this non-Bayesian behavior, we performed a clustering on these functionally distinct programs based on the similarity of the sequences that they produced, and found that this subset of programs generates several functionally-distinct classes of behavior (Figure 4c).

To further understand how these different functional groups relate to differences in program structure, we designed a second tree embedding that preferentially assigns a small and functionally-similar parent program to each children programs, provided again that they are within a single mutation of one another. This functional tree embedding (fTE) attempts to create a functionally-smooth space; enforcing such smoothness ensures that any functional discontinuities in this space arise from single structural mutations that are necessary to generate functionally-distinct behaviors (see Figure 4d for an example of how such a mutation alters the behavior of a program). Figure 4e shows this fTE performed on the programs within the GPN, with nodes colored according to the functional classifications shown in Figure 4b-c. We find that the set of functionally-Bayesian programs are clustered around the root node of the tree (corresponding to the WSLG program), whereas those programs with distinct behaviors largely occupy regions near the leaf nodes. Individual mutations (i.e., edges between nodes) alter both the structure and the functional repertoire of individual programs; we define a “key” mutation as a mutation for which a descendant program can be easily distinguished from its parent using short behavioral sequences. We find that there exists only a small ensemble of key mutations that lead to large functional differences, and we distinguish those from the remaining much larger ensemble of “sloppy mutations” that do not lead to large functional differences (inset of Figure 4e; see SI Section 10.5 for details regarding the identification of key mutations).

### Sloppy mutations create structural diversity; key mutations create functional diversity

The previous results highlight two different ways to create diversity within the space of small programs: the first is through key mutations, which generate functional diversity; the second is through sloppy mutations, which generate structural diversity. To better illustrate the nature of this diversity, we highlight one particular local view of the GPN that is anchored to a single key mutation (Figure 5a). Each key mutation significantly alters the functional repertoire of a program, and persists through the repertoires of its descendants; this is because each key mutation is embedded within a neighborhood of sloppy mutations that do not largely change the functionality of individual programs, and thus enable a function to persist through many generations of programs.

**Figure 5:**
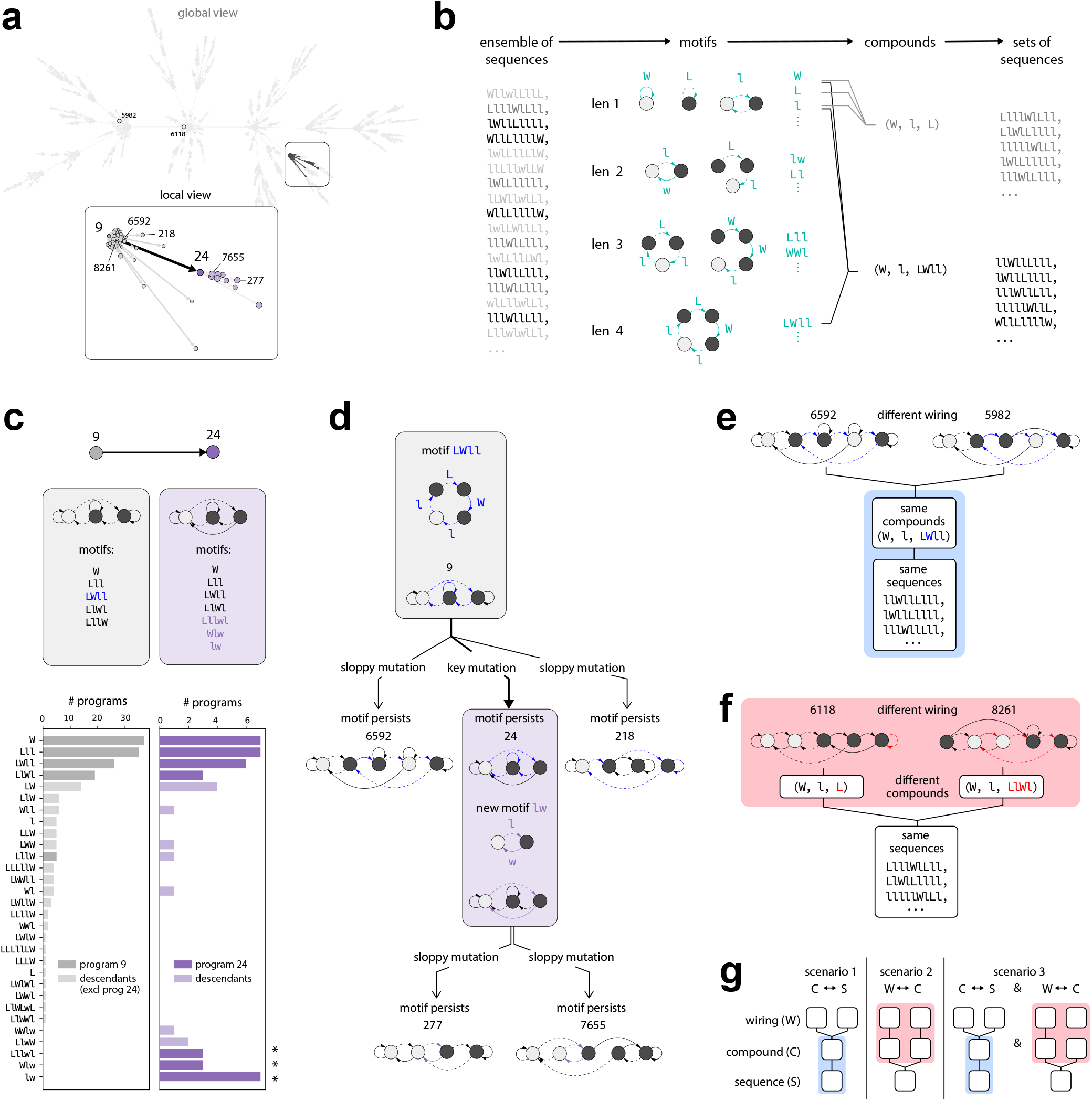
Compositional properties of functional motifs give rise to a large number of good solutions. **(a)** A single key mutation from program 9 to 24 is embedded in a local branch of programs. Two example programs (5982 and 6118) on distant lineages of the tree share some of the functions that exist within this local branch. **(b)** We perform a motif decomposition (See SI Section 10.7) to directly extract functional motifs from all behavioral sequences of a single program; these motifs are expressed in terms of outcome-action loops. Each sequence is then decomposed into a compound set of these extracted motifs. Together, these motifs and compounds represent the functional identity of a specific program. **(c)** An example key mutation creates multiple novel motifs within program 24 (starred bars in purple histogram) that are inherited by its descendants. These novel motifs distinguish the function of this lineage from the descendants of program 9 (gray histogram). **(d)** An example lineage that highlights the creation of novel motifs and the inheritance of existing ones. Motif `LWll` from program 9 (highlighted in blue) is largely inherited by its descendants through sloppy mutations. Program 24, as a descendant of program 9, not only inherits these common motifs, but also creates novel ones via a key mutation; `lw`, highlighted in purple, is one such novel motif. These novel and functionally distinct motifs are then inherited by the descendants of program 24 through sloppy mutations. **(e-f)** Two examples of functional convergence between a descendant of program 9 and a program on a distant lineage. **(e)** A functional convergence can happen when two different wirings generate the same motif compounds, which then generate the same sequences. **(f)** Alternatively, a functional convergence can happen when two different wirings generate different motif compounds, nevertheless produce the same sequences. **(g)** The fact that both scenario 1 (exemplified in (e)) and scenario 2 (exemplified in (f)) co-exist within the GPN (as shown in scenarion 3) highlights the fact that the descriptive level of “motif” cannot be merged with either the level of “wiring” nor the level of “sequence”, and thus emerges as a non-redundant description of behavior.

We define this functional repertoire in terms of the set of sequences that each program produces; for sequences defined up to length 10, this repertoire consists of hundreds of thousands of sequences. In order to capture the essence of such a large ensemble, we represent sequences in terms of functional motifs that are closely linked to the structure of individual programs (Figure 5b; see SI Section 10.7 for more details). By decomposing an ensemble of sequences into its constituent motifs, we can represent each program by a functional fingerprint that consists of tens of such motifs. We can then measure how this fingerprint changes under a key or sloppy mutation.

Figure 5c highlights one program (program 9) before a key mutation; all but one of its descendants share a similar functional fingerprint after a sloppy mutation (Figure 5c, gray histogram). One of its descendants, program 24, differs from program 9 by a single key mutation (bold arrows in the inset of Figure 5a) that endows the program with an additional set of new motifs (starred bars in the purple histogram of Figure 5c) that persist within its descendants. Figure 5d highlights one motif within this new set, and illustrates how this motif is implemented within two structurally-distinct descendants of program 24 (purple box in Figure 5d; gray box highlights the same type of structural diversity for program 9). As a result, a diversity of structurally-distinct programs can either 1) use the same set of functional motifs to generate a similar ensemble of behavioral sequences, or 2) combine motifs from different sets to generate those sequences. We can capture this additional layer of variability by decomposing individual sequences into a compound set of motifs (Figure 5b); in this way, the motif compound captures how a single program strings together multiple motifs, which can in turn generate a diversity of different sequences. We highlight this point with two examples: Figure 5e shows how the same set of sequences can be generated by a compound whose constituent motifs are implemented differently in two structurally-distinct programs, whereas Figure 5f shows how the same set of sequences can be generated by two different compounds whose distinct motifs are implemented differently in two structurally-distinct programs. If all structural and functional diversity is of the type illustrated in the first example, there is no need to distinguish sequences from the compounds that generate them (left column of Figure 5g); conversely, if all diversity is of the type illustrated in the second example, there is no need to distinguish the structure of programs from the compounds that they generate (middle column of Figure 5g). However, the GPN exhibits both types of diversity (right column of Figure 5g), and thus motifs emerge as an intermediate level of description that can capture different relationships between the structure of individual programs and the behavioral sequences that they generate. Altogether, our results highlight how diversity in the space of small programs can arise at multiple different levels, through structural mutations that give rise to different functional motifs, and through the combinations of those motifs that give rise to behavioral sequences.

## DISCUSSION

To guide behavior in the face of uncertainty and change, animals can leverage knowledge about their surroundings in order to make inferences and guide actions [27, 28]. There are in principle many possible ways that the brain can leverage such knowledge, depending on resource constraints and performance demands. Here, we focused on the important role that resource constraints can play in guiding “good enough” performance. We focused on a well-studied behavioral task in which animals can use inferences about a changing state of the environment to guide future actions [18, 19, 20]. Rather than constructing individual strategies that could be used to best solve this task, we enumerated and studied relationships between all possible strategies that achieve better than chance performance. The focus on efficiency and robustness [29, 30, 31], rather than strict optimality, is conceptually similar to earlier AI approaches that explored heuristics for efficiently solving particular tasks [32, 33, 34]; the focus on the discovery, rather than the construction, of candidate behavioral strategies bears some similarity to model deduction from animal or human behavior [35, 36]; the focus on studying a spectrum of good solutions, rather than a single optimum, is conceptually related to work that formalizes the degree of optimality of a system [37]. By taking this approach, we enumerated a complete space of good solutions, and we used this space to study relationships in the structure and function of different solutions that underlie good performance.

Our approach differs from those that aim to optimize and reverse-engineer a single black-box model for solving a particular task. Instead, we focused on extracting insights from relational structures observed across the entire space of good solutions. A tree embedding of this space revealed that nearly all good solutions are closely connected through single mutations; not only did this relationship between wiring and performance enable us to devise an efficient evolutionary search algorithm, it revealed the “smooth” nature of this space—suggesting that there could exist other highly organized relationships in structure and function that underlie good performance. Consistent with this idea, we showed that a second tree embedding could be used to extract functionally smooth relationships among all good solutions, and in doing so we uncovered several functionally-distinct regions of solution space whose behavior was qualitatively distinct from the set of optimal Bayesian solutions. It is worth emphasizing that while this approach enables us to summarize relational structures, we did not attempt to reduce the high-dimensional nature of this space into a low-dimensional description. Instead, we used these minimal relational structures to traverse different evolutionary lineages in a hierarchical manner while preserving detailed knowledge about individual solutions along each lineage. As such, this approach differs from more commonly-used clustering approaches that provide a low-dimensional summary of a space by averaging away individual variability.

Together, these results highlight that the behavioral repertoire for this task is large, and the behavior of individual solutions can deviate from the norm without significantly compromising performance. This behavioral freedom resembles the notion of sloppiness observed in multiparameter models [38, 39, 40]. Furthermore, provided that a local structural relationship between two solutions contains information about their functional difference, we were able to uncover the *origin* of functionally distinct solutions as an accumulation of a handful of key mutations that can alter patterns of behavior, and we were similarly able to understand the *persistence* of these distinct solutions in terms of the much larger ensemble of sloppy mutations that do not alter functionality. Together, this characterizations highlights how structural and functional diversity among good solutions emerges from sloppy and key mutations, respectively.

While our focus here was on enumerating and understanding relationships between the many different solutions that could be used to solve a single task, it is easy to imagine the reverse, in which one studies how a single versatile solution might perform on multiple different tasks [41]. While such an approach is commonly used to study the notion of generalizability, studying a single solution in isolation precludes the ability to identify which features of that solution are specifically required for generalization. Instead, the approaches adopted here could be used to construct and study a joint task-solution space in which many tasks collectively map onto many solutions (see SI Part I for more discussion on this point). In this way, the relational structures in such a joint space could be used to identify the minimal components needed to capture the diversity of generalizable behavioral strategies that are ubiquitous in biology.

## ACKNOWLEDGEMENTS

We thank Rishika Mohanta, Maanasa Natrajan, Max Manakov, Hanqing Wang, Jinyao Yan, Diptodip Deb, Angel Stanoev, Boaz Mohar, Sue Ann Koay, Marcella Noorman, and Yipei Guo for many valuable discussions. This work was funded by the Howard Hughes Medical Institute.

## AUTHOR CONTRIBUTIONS

Tzuhsuan Ma and Ann Hermundstad conceived the study, formalized the conceptual and theoretical framework, developed the enumeration, evaluation, embedding, and evolutionary algorithms, conducted the mathematical analysis, performed the simulations, interpreted the results, and wrote the paper.

## CODE AVAILABILITY

Code was written in Python and is available at https://github.com/HermundstadLab/ProgEnum

## SUPPLEMENTARY INFORMATION

### Part I IDEAS & LESSONS

#### 1 Why adopt a Tree Embedding (TE)?

##### 1.1 Breaking down a large dataset into manageable chunks

One major goal of doing an embedding for a large dataset is to uncover hidden structures among individual entities by decomposing the entire dataset into some smaller and more manageable chunks. For example:

1. An embedding algorithm can group entities based on how similar they are along a certain common attribute, say, “performance.”
2. A different grouping could emerge as a result of using another attribute, say, “structural complexity.”
3. Alternatively, a similar grouping could emerge from a third attribute, e.g. “functional difference from a Bayesian agent.”

From the above three embeddings, one extracts certain relations among these attributes, i.e., “structural complexity” does not predict “performance,” but “functional difference from Bayesian” does. And since these common attributes are assigned to individual entities, one can successfully capture the relationships among individual entities as well. Moreover, after establishing these relational structures, one is left with multiple ways of grouping entities, and can thereby zoom in and analyze each group separately.

##### 1.2 Deciding which relations to capture

For our purpose, the relevant attributes for describing a program can be:

1. attributes related to the *structure* of a program:
  a. program size
  b. number of mutations from a base program (e.g., discrete Bayesian program (DBs), the WSLG program, etc.)
  c. wiring features (e.g., bidirectional integration, action-outcome loops, etc.)
2. attributes related to *function* of a program:
  a. performance
  b. joint state-belief occupancy
  c. action-outcome statistics

As an example, our tree embedding (TE) algorithm uncovers the following relational structures:

1. Nearly all *high-performing* programs are closely connected through single *mutations;*
2. A good program is likely to *mutate* into another *functionally* similar program;
3. A large *functional* change only happen occasionally, and following a rare *key mutation;*
4. Nearly all *functionally-distinct* programs are clustered close to leaf nodes, because it takes a several *key mutations from the root* to accumulate sufficiently many new behavioral sequences to become functionally distinct.

**Figure S1:**
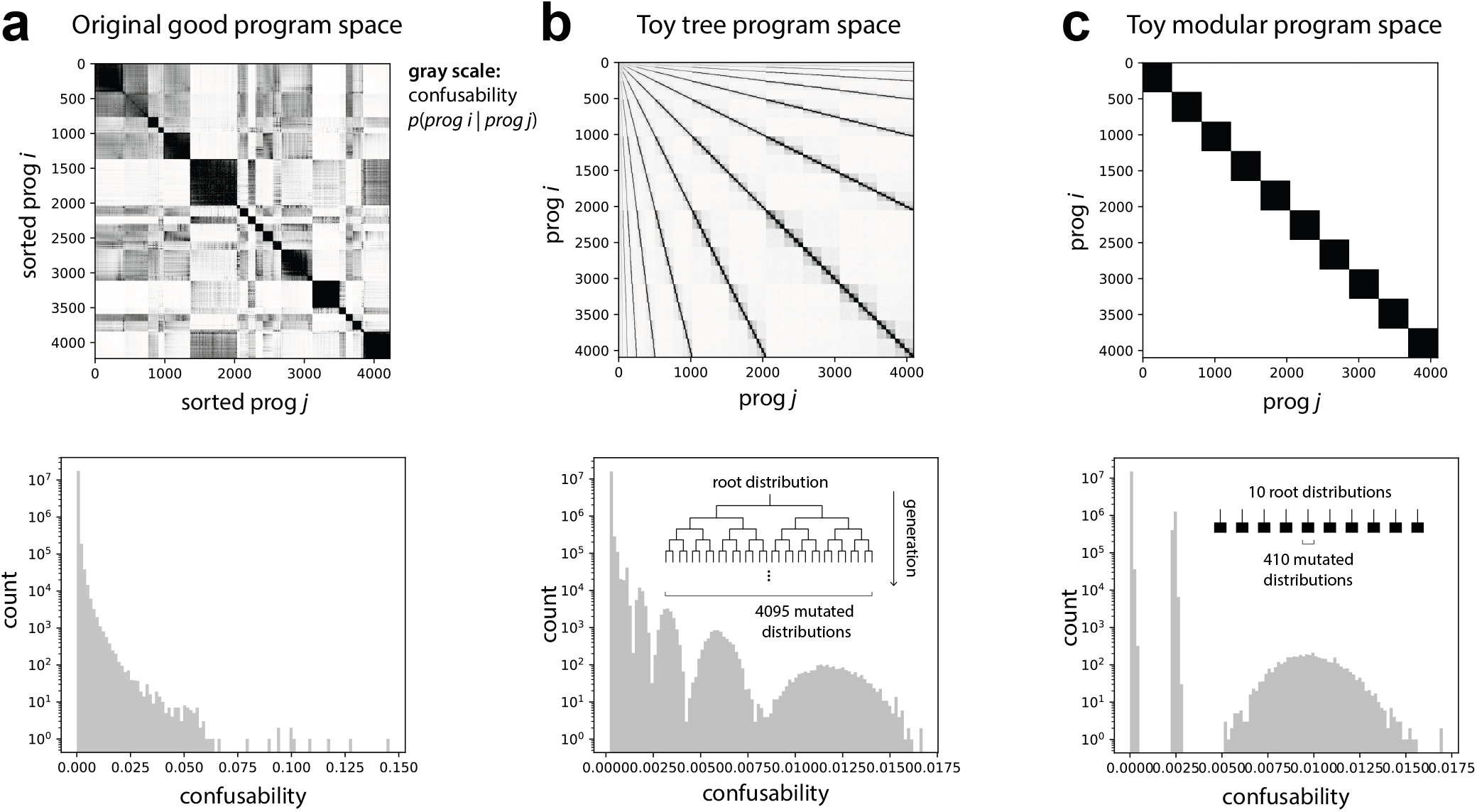
The structure of the good program space is less modular, and more hierarchical. **(a)** The confusion matrix of the good program space, sorted with the `community_louvain` [26] clustering algorithm. The matrix is not fully block-diagonal, which indicates that the space is not fully modular. The lower panel shows the histogram of all values in the matrix. **(b)** A toy tree program space contains 4095 hierarchically-connected distributions. This space produces a non-modular confusion matrix (upper panel), and the corresponding distribution of entries has a long tail (lower panel). **(c)** A toy modular program space contains ten orthogonal groups, each with 410 programs (upper panel). The corresponding distribution of entries is multimodal, rather than long-tailed (lower panel).

##### 1.3 The space of good programs is endowed with hierarchical structure

In this work, we decided to adopt a TE algorithm after realizing that the program space itself is inherently non-modular. This realization came about after we performed TSNE on the program space (discussed in more detail below). Instead of being modular, we observed that the majority of relevant structures within the program space were hierarchical. To see this, we demonstrate that 1) a toy program space with an *iterative branching structure* (i.e, a tree) can better capture the original action-outcome statistics than 2) a toy program space with a *modular structure* Figure S1.

###### The original program space is not modular

In Figure S1a, one can see that the functional confusion matrix from all 4230 good programs does not have a clear block-diagonal structure (note that these programs are sorted using `community_louvain` algorithm), which suggests that the good program space cannot be broken down into a few mutually orthogonal clusters within which programs are functionally similar. To support this observation, one can see that the corresponding histogram (Figure S1a, lower panel) shows a continuous, long-tailed distribution. This long tail implies that:

1. the size of program clusters, if they exist, could vary extensively (e.g., a larger cluster—within which most programs are confused with each other—would contribute to lower values of the distribution, whereas a smaller cluster would contribute to higher values);
2. most clusters, if they exist, could be partially mixed with other neighboring clusters, yielding an overall distribution that is smooth.

If the latter factor dominates the behavior of the program space, it is not particularly helpful to view the program space as modular.

###### A toy tree program space displays a long-tailed distribution of confusability values

Figure S1b illustrates the confusion matrix for a toy tree program. To generate this space, we grew a toy tree from a root program with a sparse random distribution (a normalized binary random pattern) of behavioral sequences. Next, we added a small binary noise to the root distribution `(0+1=1, 1 + 1=0)` to generate two descendants. These two newly-generated distributions differ slightly from their parent. We then apply this rule to iteratively mutate distributions up to 11 generations, thereby creating a space of 1 + 2+2^2^ +… + 2^11^ = 4095 programs. In Figure S1b, one can see a nested non-modular structure in the confusion matrix, and the corresponding long-tailed distribution within each successive generation creates a new lump of slightly larger values of confusability. This distribution closely resembles what we observe in the original space of good programs (Figure S1a).

###### A toy modular program space displays a multi-modal distribution of confusability values

A counterexample is provided in Figure S1c, where 10 orthogonal distributions are seeded to generate an entire set of toy programs. By adding moderate noise to one of the distributions, we fill in one-tenth of the program space. By doing the same for remaining 9 distributions, we have a complete modular program space. To see how the histogram in Figure S1c comes to be, we can imagine a toy space without adding any additive noise. In such a space, all programs should have the same level of confusability within their own groups; this would manifest as a single, delta-function spike at the location of the middle peak in the lower panel of Figure S1c (this single spike emerges because all clusters have the same size). If one then adds noise to each cluster, some fraction of this peak will shift to the right, thereby forming an overall trimodal distribution. This distribution much less resembles what we observe in Figure S1a than does the toy tree example above.

In retrospect, the lack of modular structure in the good program space is not altogether surprising, given that several relevant program attributes operate within near-continuous domains:

1. The full program space is inherently unstructured because of the inclusion of all possible deterministic Markov chains.
2. For a large enough program space (268,536 programs with *M* ≤ 5), the task can be solved in virtually infinitely-many ways, and thus the performance spectrum is effectively continuous.

Building upon the ideas above, one can infer that there is no obvious way to define a performance threshold that breaks apart the full program space into isolated clusters with similar performance.

##### 1.4 Tree embeddings with different objectives can capture different hierarchical structures

Knowing that the structure of the good program space is better described as hierarchical than modular, it is reasonable to perform a tree embedding on the program space to extract that structure (in the form of relationships between programs). It’s important to keep in mind that there could be more than one hierarchical structure embedded within the program space, and implementing different objectives for a tree embeddingcould reveal different hierarchical structures. For example, in the main text, we perform two different tree embeddings: one based on performance (TE), and another based on function (fTE).

TE has two objectives: 1) preferentially find a parent program of a smaller size, and 2) preferentially find a parent program with a higher performance. Applying these two objectives reveals that programs with similar *performance* and *wiring* are hierarchically connected together (Figure 3f in the main text).

fTE, on the other hand, has a different set of objective: 1) preferentially find a parent program of a smaller size, and 2) preferentially find a parent program with a similar set of behavioral sequences. Applying these objectives reveals that most mutations are sloppy; However, a small set of key mutations scattered throughout the tree is sufficient to generate a large number of functionally-distinct programs near the leaf nodes of the tree.

##### 1.5 Tree embeddings provide a useful way to make sense of a high-dimensional space

It is worth pointing out that by performing an embedding, one often forces a certain artificial structure onto their dataset. Although we argue that the inherent structure of the program space is hierarchical, the minimal relational structure that is captured by TE or fTE is not guaranteed to fully capture the structure of the program space. To see this, consider the fact that a tree has on average only has two neighbors per node; one parent and one descendant for each node. Yet, individual programs in the good program space can have between 2 and 340 single-mutation neighbors (for an average of 14 neighbors). Despite the fact that TE does not represent all aspects of the program space, it has enabled us to extract several different insights from the program space, as discussed above.

More broadly, this type of TE could be a useful tool in making sense of other high-dimensional spaces, and could generate different insights than commonly-used clustering, dimensionality reduction, or embedding algorithms. This is because:

1. A true high-dimensional space has certain irreducible attributed assigned to individual entities within the space.
2. Conventional clustering algorithms typically reduce the dimensionality of a space by erasing these individual attributes.
3. A TE instead focuses on simplifying the relationships among entities, while preserving their individual attributes.

In a way, the goal of TE is to extract some relational structures that could, in turn, comprise a set compact rules for iteratively generating the high dimensionality of the original space. As a result, it could be a more compatible approach for understanding a high-dimensional space in its entirety. Beyond extracting a better understanding from such a dataset, one can imagine translating such a relational structure into a generative or search algorithm. In the main text, we discuss how an evolutionary algorithm can efficiently discover good programs by searching only a fraction of the full space. In Section 4, we discuss another potential application for a more efficient program enumeration based on outcome-action motifs.

##### 1.6 Limitations of a commonly-used clustering algorithm, TSNE

Above, we discussed why conventional clustering algorithms might be less useful in extracting relational structures in high-dimensional spaces. Here, we show results from one of our initial attempts to making sense of the program space using TSNE. Having gone through this exercise without much progress, we decided to design our own tree embedding algorithm.

The first problem we need to address before using TSNE is to construct feature vector of the same size for programs of different sizes (note that this issue of program-size compatibility is something that we need to solve in designing TE as well; see Section 9.1 for details). We construct a compatible feature vector for all programs using the following steps:

1. We note that every transition in a program can be labeled by a 2-tuple (*O, A*), where *O* ∈ {win, lose}, and *A* ∈ {stay, go} (the WSLG program exemplifies this in its naming).
2. This labeling, however, is ambiguous, and it largely erases the wiring structure of a program (e.g., a “stay” transition can happen either through a self-loop transition, or if the destination node is labeled with the same action).
3. To preserve more structural information, we thus used a 3-tuple (*O, A, D*) to specify a node instead. In this tuple, *D* ∈ {0, 1, 2, 3, 4} measures distance to a destination node, with *D* = 0 denoting a self-loop-transition. As an example, in Figure S2, we show 3-tuples (formalized as discussed below) and the corresponding feature vectors for the WSLG program, and for the top-performaning discrete Bayesian (DB) program.
4. For each program, we find a specialized node ordering that minimizes the transition distance. The idea behind this representational formalization is to make all programs as similar as possible, such that the distance is most likely to capture real difference between the underlying graphs (invariant upon different representational choices). Having done this, we proceed with TSNE using this formalized feature vector.

We need one last step to formalize the node ordering, because two programs can be either structurally similar or distant depending on which representation one chooses among all possible node permutations (note that we don’t have this problem in designing TE, since the number of mutations from one program to another is invariant across all possible representation).

**Figure S2:**
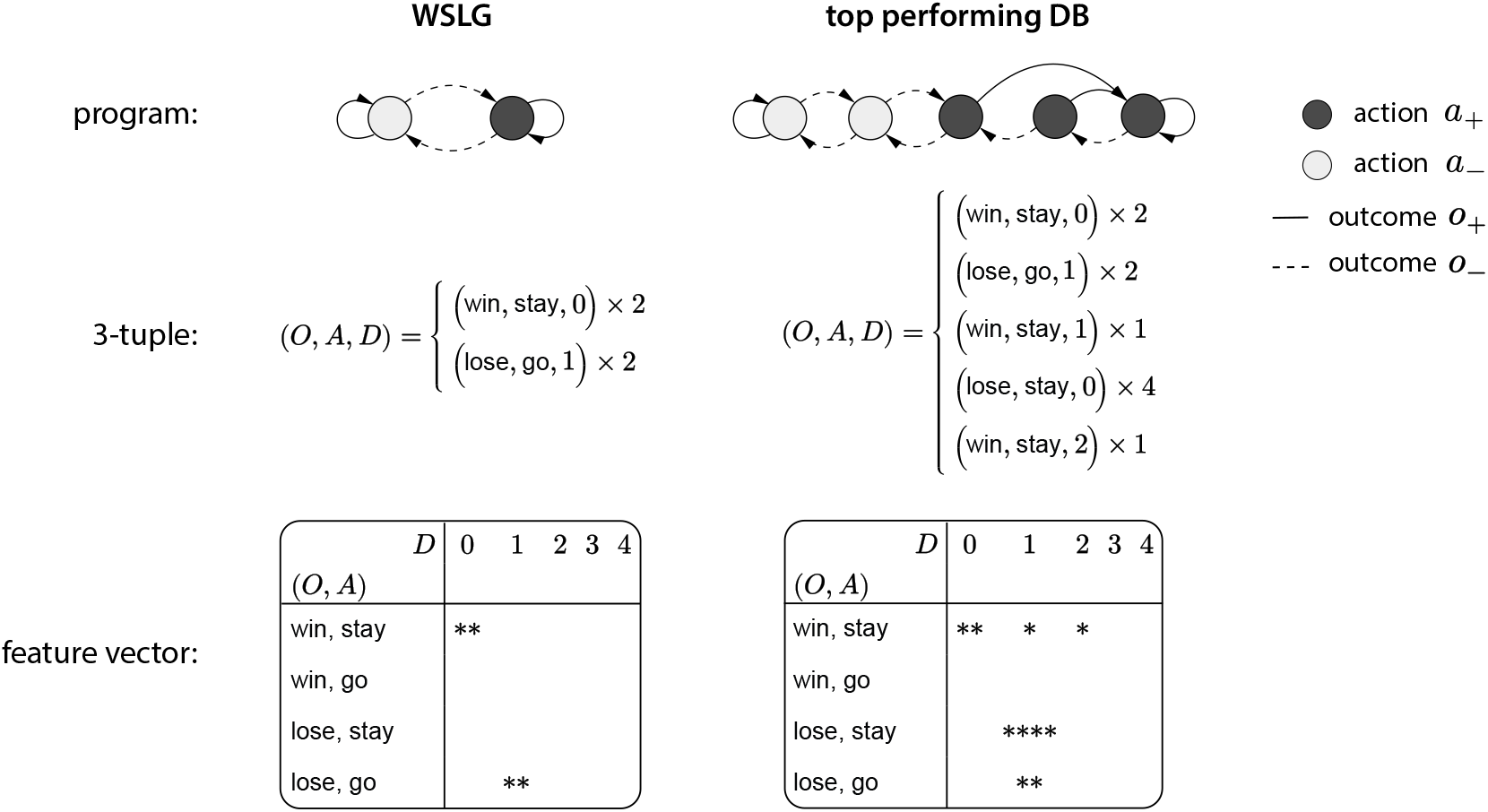
TSNE requires a formalized feature vector (for wiring) that is compatible with all program sizes. The feature vector is a flattened version of a one-hot structure matrix, as shown in the bottom row.

**Figure S3:**
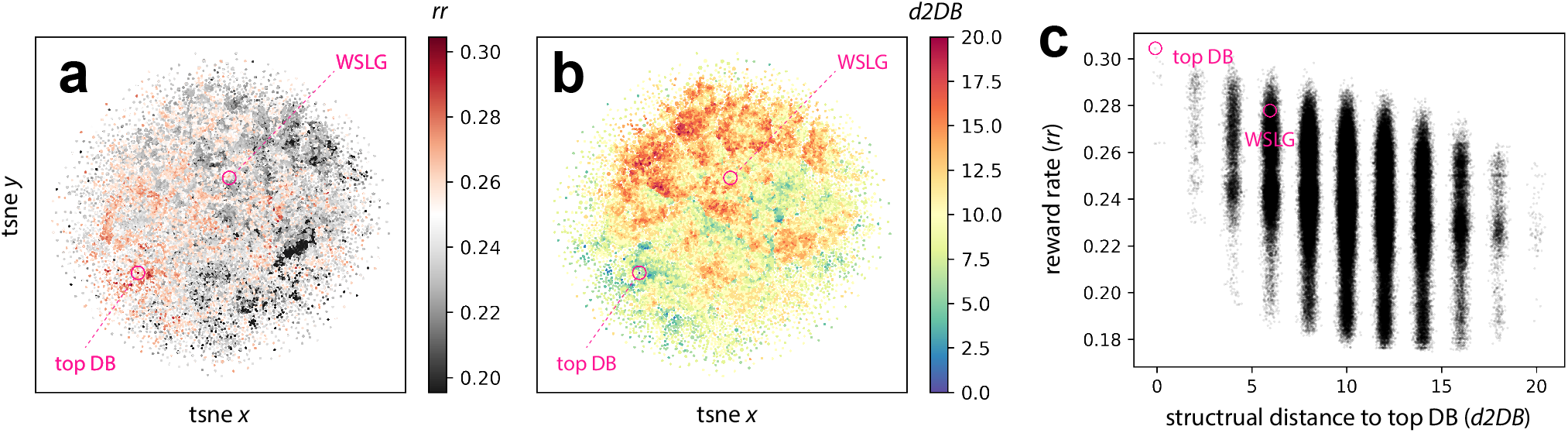
TSNE on the full program space. **(a)** The embedding with each program colored by its performance. **(b)** The embedding with each program colored by its structural distance from the top-performing discretized Bayesian (DB) program. **(c)** The scatter plot shows a rough trend between lower performance and larger distances from the top DB.

From the TSNE results shown in Figure S3, one can make a few immediate observations:

1. Without coloring the nodes, the embedding does not show a clear sign of clusters or hierarchical clusters.
2. Coloring each node with either its “reward rate” or “distance to the top performing DB” reveals that some intricate relationships between a program’s structure and performance, but the color separations are largely salt-and-pepper in nature.
3. Overall, if a program is closer to the top performing program (a discretized Bayesian), it is more likely to perform well (this conclusion can be reached from Figure S3c, even without doing TSNE).

From the above, one can see that TSNE serves as a useful first analysis step in order to obtain some intuition about a dataset. In our case, TSNE shows that there is not any obvious clustering structure that one can use to break down the program space into manageable chunks. This motivated us to adopt the tree embedding algorithm, rather than to use or customize an existing clustering algorithm.

Finally, in the table below, we summarize and compare some aspects of TSNE and TE applied to our problem:

**Table S1:**
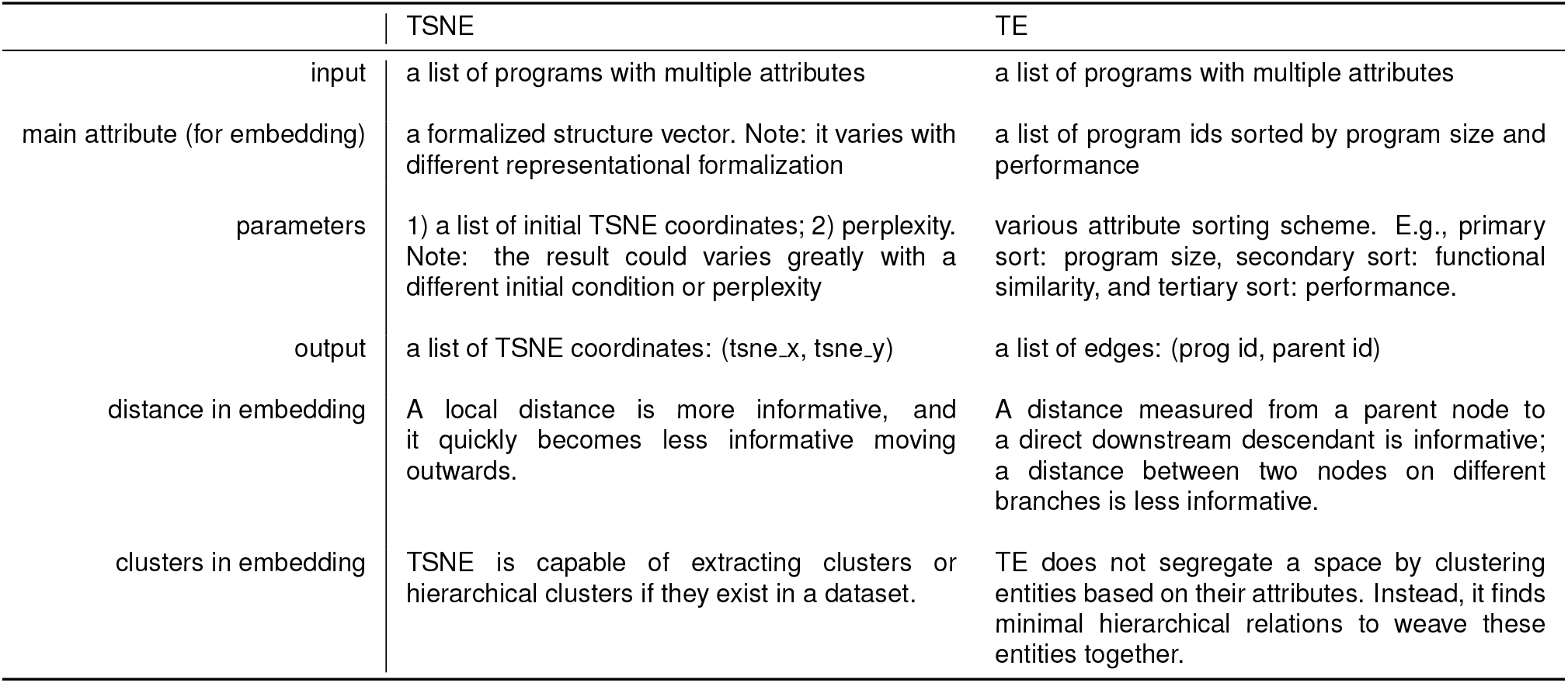
TSNE vs TE.

#### 2 Notions of task and function sloppiness, and how they could aid the study of generalizability

In the main text, we focused on a fixed task with changing functions along evolutionary lineages. To study generalizability, one would need a complementary view as well. Specifically, we ask how certain functions among good solutions persist under changes to a task. To further this discussion, we also introduce a two different notions of sloppiness in either a function space (function sloppiness) or a task space (task sloppiness), with the goal of describing generalizability in those terms. But first, we discuss how generalizability could be understood by studying relational structure in a solution space.

##### 2.1 The value of studying relationships among good solutions

In the main text, we discussed how generalizability is often studied by constructing and dissecting a single versatile solution that is capable of performing multiple tasks. While such an approach provides a realizable solution that can be immediately deployed to a practical problem, we argue that it offers limited insight into how generalizable behaviors work.

To see this, consider searching for a solution X that can solve both Task 1 or 2 under some resource constraint. It turns out that one cannot make any useful statements about generalizability without contrasting this one solution to something else, even though this solutions fits our intuition of what a generalizable solution is. To overcome this limitation, consider searching for two solutions Y and Z that can solve Task 1 and 2 respectively. Now if one contrasts solutions X and Y, or contrasts solutions X and Z, one can make statement such as:

> *“Nearly all behaviors commonly shared between X and Y do not exist in Z, and vice versa; this suggests a strong tradeoff between implementing behaviors for Task 1 and those for Task 2; the generalizability across these two tasks can then be quantified in terms of this tradeoff.”*

From even this simplest example, one can already see the importance of studying relationships among different solutions if one wants to study generalizability. Building upon this argument, we believe that capturing various relational structures in both task and solution spaces remains the key to understanding generalizability in various settings and problem domains. For this reason, we propose that an enumeration approach capable of discovering many good solutions, in combination with tools that can extract relational structures among those solutions, is best suited to study generalizable behavioral strategies.

##### 2.2 Task and function sloppiness are closely related

When thinking about a task and a solution as a point in space, it is rather difficult to intuit how changing one affects another. It is a very different matter when both the task and its solution are parameterized together (so that dialing this set of parameters changes both the task and solution); by varying these shared parameters, one effectively creates a task and a solution space. An example of such a case is an Bayesian agent optimized on a parameterized task; in this case, both the Bayesian observer and the task are parameterized identically (e.g., in terms of the hazard rate, reward contrast, and baseline reward rate in our two-armed bandit task). However, one would like to go beyond a Bayesian observer, given all of the reasons that we have highlighted in favor of studying relational structures of a whole solution space rather than dissecting the mechanisms within a single solution. For that, one needs to establish a connection between a task space and a solution space, sometimes without an explicit shared parameterization.

The tools and conceptual framework that we have been building aim to make such a connection. More specifically, we use an enumeration scheme to find the good solution space for a fixed task; we then perform Motif Decomposition (MD) on these solutions to create the corresponding *function space* (behavioral repertoire). Next (as discussed below), we can morph the task in a *task space*, and we can repeat the above two steps to observe how the function and task spaces change together.

Having conceptualized the connection between a task space and a function space, one can now see how sloppiness in the two different spaces might relate to one another.

1. By fixing a task, sloppiness in function space means that changing behaviors along a certain function axis does not compromise performance—we have observed this in the functional tree embedding, where one can traverse different lineages to reach a certain set of functions without compromising performance.
2. By fixing a set of functions, sloppiness in task space means that changing an aspect of the task along some parameterized axis does not compromise performance.

From the points above, it could even be more sensible to talk about sloppiness in a joint task-function space in which many tasks map to many functions and vice versa. In fact, this many-to-many mapping between tasks and functions resembles the mapping between perturbations and responses in robust biological systems [42]. Having said that, it remains to be seen as to whether this view of joint space is a useful conceptual framing.

**Figure S4:**
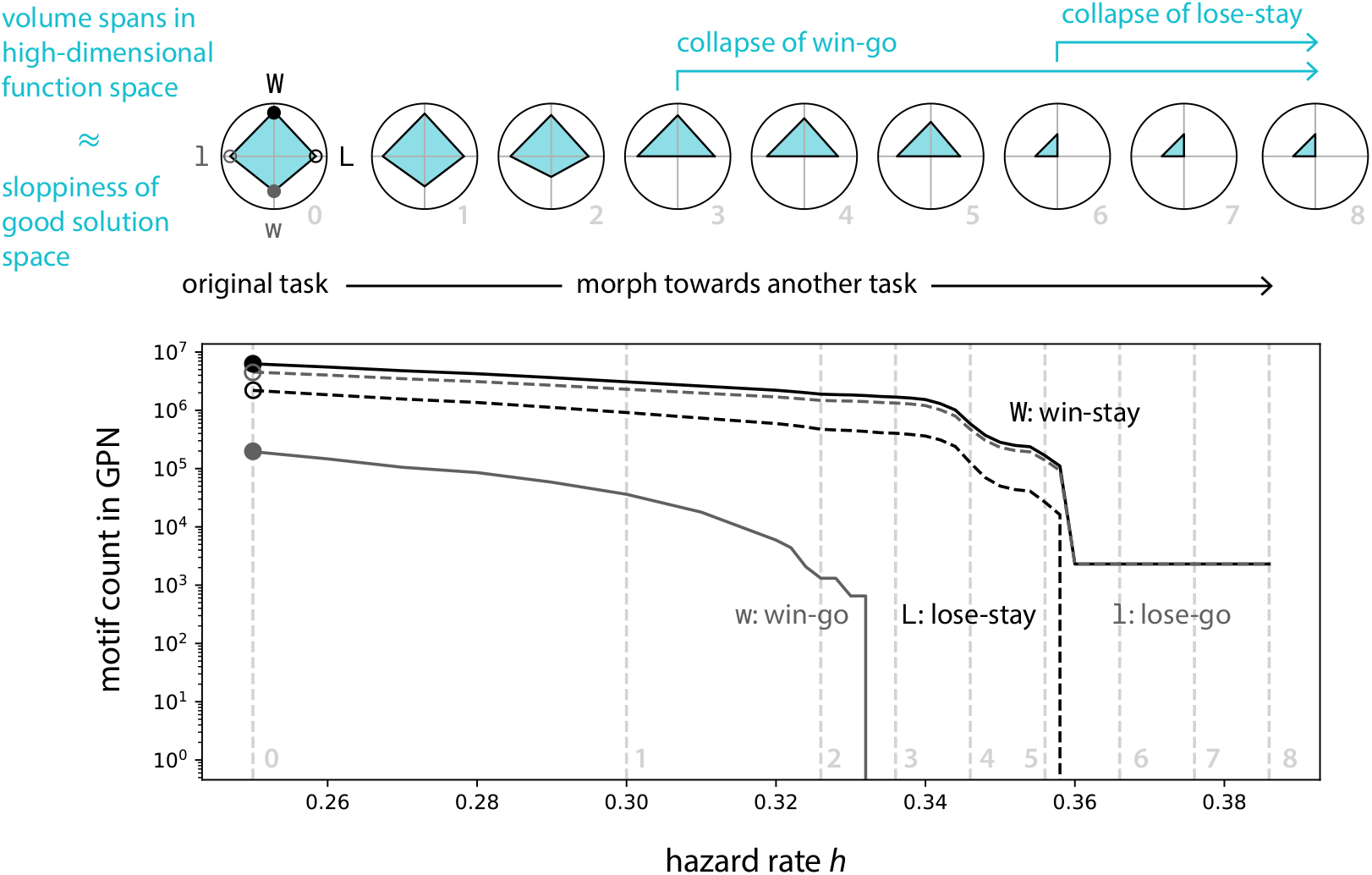
Generalizability could be understood in terms of how different functional axes vary with task structure. We use the lowest-order motif statistics to represent existing functions of the good solution space. When a task is morphed towards some unknown task along a certain axis (in this example, we use the hazard rate to illustrate this point), certain functions collapse more quickly than others. Those persisting functional axes therefore represent behaviors that can be generalized to another task that lies along the morphing direction.

##### 2.3 Persistent axes in high-dimensional function space hint at generalizable behaviors

To make these ideas more concrete, we use a minimal example to illustrate how the notion of sloppiness could connect to generalizability. In Figure S4, we plot the statistics of lowest-order motifs, and we observe how the “shape” of the function space (cyan polygon) changes with the task (an increasing hazard rate). Conceptually, morphing this one aspect of the task would point towards another qualitatively different task in a task space. A generalizable behavior then refers to a persistent set of functional axes in the function space. For example, “win-go” is a function that exists only transiently, and thus would not generalize well toward another task along the morphing axis, whereas “lose-stay” persists for much longer.

In ongoing efforts, we are trying to formalize this conceptual framework, and use it to better understand the origin of generalizability as a feature shared among many biological computations and behavioral strategies.

#### 3 When and why to use Motif Decomposition (MD)

Most generally, Motif Decomposition (MD) aims to bridge observed behaviors with the underlying rules that generate them. For example, in connectomics, the goal would be to link synaptic connectivity with neural activity. In the main text, we perform MD to bridge the structure a program (wiring) with its function (behavioral sequences). The details of the MD algorithm are discussed in Section 10.7. Here, we try to outline some insights that we obtained from performing MD on our problem.

##### 3.1 Motifs bridge structure and function

Provided that we define a motif to be an “action-outcome loop” in a long behavioral sequence, and provided that a loop is also directly depicted in the wiring of a program, it is straightforward to see how a change of wiring can cascade to make changes of motifs that eventually lead to changes of sequences. This enables us to understand how a single key mutation (a small change in structure) can modify behaviors to such a large extent.

It is import to point out that the above reasoning doesn’t justify the choice of using loops as motifs. As an alternative, one might use, say, all possible short sequences with a fixed length (i.e., n-grams, as discussed in Figure S5) for a decomposition:

**Table S2:**
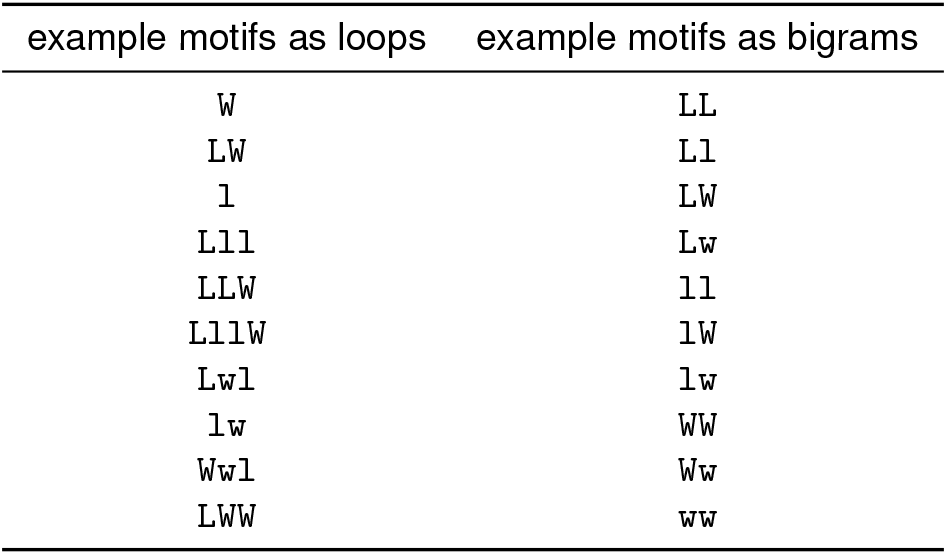
example motifs and bigrams. `W`: win-stay, `L`: lose-stay, `w`: win-go, and `l`: lose-go.

| example motifs as loops | example motifs as bigrams |
| --- | --- |
| W | LL |
| LW | Ll |
| l | LW |
| Lll | Lw |
| LLW | ll |
| LllW | lW |
| Lwl | lw |
| lw | WW |
| Wwl | Ww |
| LWW | ww |

However, this somewhat arbitrary choice of using bigrams does not explicitly take advantage of the structure of behavioral sequences. As a result, it is a highly redundant representation. In Section 3.3, we will quantify this redundancy when we discuss motif compositionality.

##### 3.2 The utility of Motif Decomposition

The reason that loops are such an effective representation for decomposing sequences has a lot to do with 1) the relatively small size (*M* ≤ 5) of all programs and 2) the relatively low hazard rate (*h* = 1/20) of the task. For these reasons, nearly all existing smaller loops in a program can be reached relatively quickly; a long enough sequence can then be decomposed into short motifs (a majority of which are less than or equal to length=5). As a result, a highly compressed motif representation can capture behaviors without much loss of information.

With the above reasoning, this specific motif decomposition that we adopt is not suitable for all problems. If the above two criteria are not met, motif decomposition of this particular sort is not useful. For example, a large dataset of short video clips might predominantly consist of a set of short open sequences rather than loops; in this case, it would take a very different decomposition algorithm to extract those open-ended sequence as motifs. Another example could be decomposing a set of static images into a set of feature vectors that can be linearly superimposed to reconstruct the original image. In such a case, a feedforward deep neural network serves the purpose well.

##### 3.3 Structured compositional motifs constrain task sloppiness

In the most general sense, a high degree of *compositionality* implies many different ways of constructing a *composite* from a set of *components*. In our case, figuring out how many ways a set of motifs can be combined to build a good program tells us how sloppy the solution space can be. In the discussion below, we compare Motif Decomposition (MD) with a commonly used n-gram analysis to show how MD generates motifs that efficiently capture the set of behaviors produced by the space of good programs. We then discuss how the compositional structure among all motifs constrains the sloppiness of the task—i.e., how many good solutions can exist.

**Figure S5:**
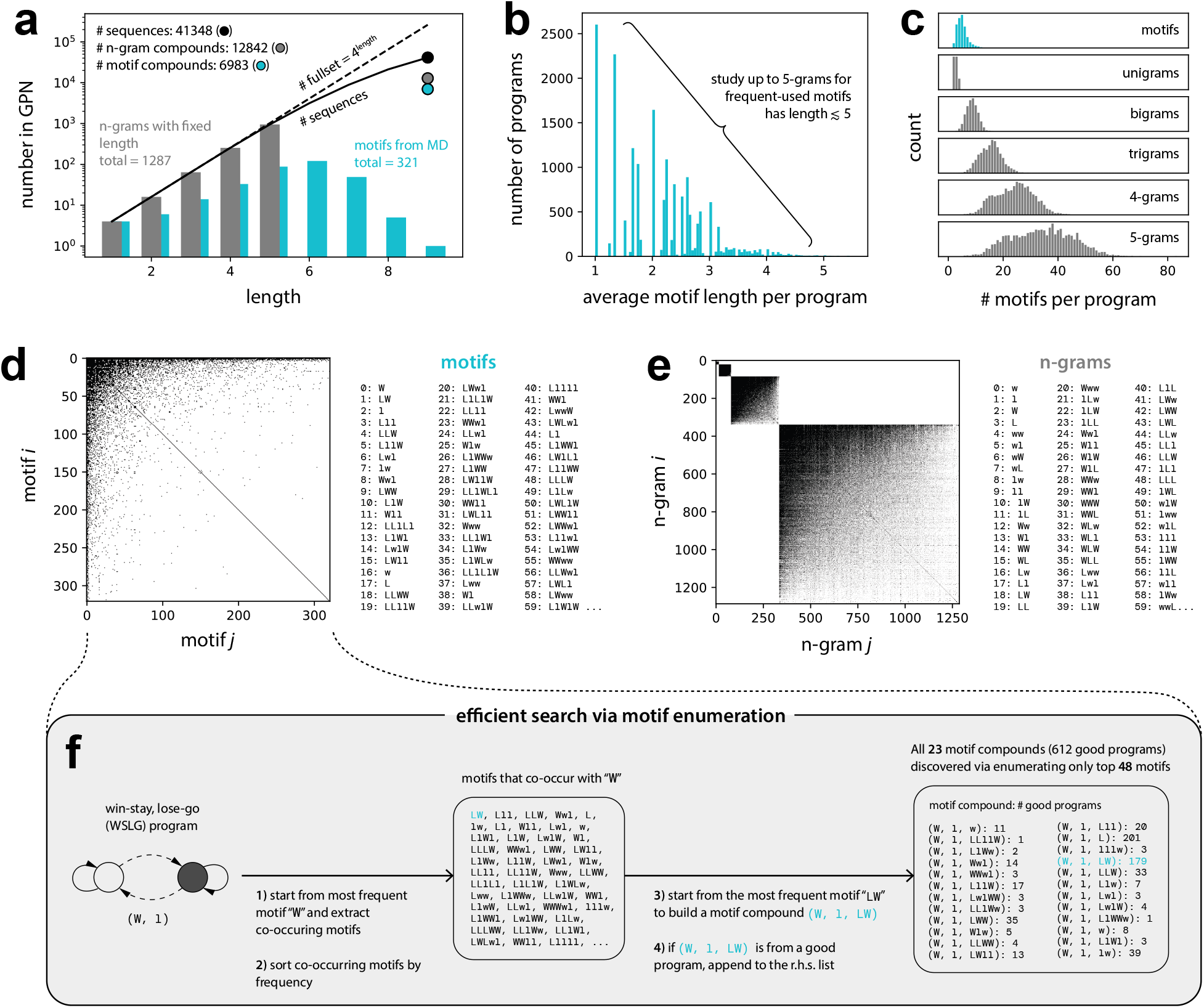
Motif Decomposition (MD) provides an efficient way to capture the behaviors of good programs and to explore a larger program space. **(a)** The set of motifs extracted from MD result in a more efficient representation of the good program space than do sets of n-grams. **(b)** We use the statistics of motif lengths to select the set of n-gram lengths. **(c)** A program can be represented with just a few motifs, in comparison to representing it with n-gram. **(d-e)** A sparse co-occurrence matrix for motifs shows that a compositional rule for motifs is highly structured. In contrast, such a rule for n-gram is much less constrained. **(f)** The structured compositional rules for motifs could enable an efficient search via motif enumeration.

###### Motif Decomposition (MD) efficiently captures the structure of the good program space

To get a sense of how task sloppiness might relate to motif compositionality, one needs to ensure that a motif compound indeed faithfully and efficiently represents a behavioral sequence. If a representation is highly redundant, it would give a false sense about how large a space can actually be. Here, we demonstrate this by contrasting the results from MD with those obtained from an n-gram analysis.

###### n-gram analysis

n-gram analysis, commonly used in language processing, is a generic approach to break down a long string of texts into multiple manageable statistical distributions. In our case, one may start from constructing 1st and 2nd-order unigram and bigram statistics:

**Table S3:** example of unigram and bigram statistics.

|  | unigram | bigram |
| --- | --- | --- |
| 1st-order stat | $p(\text{seq}) \forall \text{seq} \in \{L, W, l, w\}$ | $p(\text{seq}) \forall \text{seq} \in \{LL, Ll, LW, \dots, ww\}$ |
| 2nd-order stat | $p(\text{seq } i, \text{seq } j)$<br>$\forall (\text{seq } i, \text{seq } j) \in \{(L, L), (L, W), \dots\}$ | $p(\text{seq } i, \text{seq } j)$<br>$\forall (\text{seq } i, \text{seq } j) \in \{(LL, LL), (LL, Ll), \dots\}$ |

As one can see, unigrams and bigrams have limited expressive power in capturing the complexity of all program behaviors. To improve the granularity of these descriptions, one can consider higher-order n-grams. As shown in Figure S5a, we consider up to 5-grams (this is because nearly all of the frequently used motifs have lengths less than or equal to 5).

Note that for each program, there is a smaller set of all possible n-grams that could exist (with a nonzero probability); this set of n-grams can then be the lowest-order description for a program. The next higher-order description can be expressed as a correlation matrix for nonzero co-occurrences between two existing n-grams, as shown in Figure S5e. After sweeping from unigrams to 5-grams and from lowest-order to higher-order statistics, each program can then be collectively described by a set of n-gram statistics.

###### Motifs are highly efficient in capturing the behavior of small programs

Using the above approach, we can count how many unique n-grams are required to describe the entirety of the good program space, and we can compare that to the results obtained from MD. In Figure S5a, one can see that the total number of n-grams (1287) that one needs to capture the good program space is much larger than the total number of motifs (321), indicating that MD provides a more concise description of the space.

Next, we focus on the compositional properties of these n-grams by counting the number of different ways that n-grams can be combined to summarize a program. n-grams can be combined to form 12,842 possible compounds; motifs, on the other hand, form 6983 compounds (note that for motifs, we count how many compounds are used to describe the sequences produced by a program). Once again, this comparison shows that MD provides a more efficient description than do n-grams. Lastly, at individual program level, one can use far fewer motifs to describe any given program (Figure S5c).

It’s worth noting that MD provides a more efficient description of the good program space because it exploits the fact that all long sequences are built from a small number of relatively short action-outcome loops. MD uses this knowledge of the space to compress the set of program behaviors (summarized by a set of 41,348 sequences in total) down to a much more concise description (summarized by a set of 6983 compounds in total).

##### 3.4 Motif compositional rules are highly structured

From the results discussed above, one gets a sense about how structured the compositional rules can be for combining motifs or n-grams into permissible compounds. For example, there could in principle be up to 51,360 ways to put any two motifs together, and 5,461,280 for three motifs. Yet, the observed rules only allow 6983 motif compounds to describe the good program space. Here, we take a closer look at the properties of these compositional rules. In Figure S5d,e, one can see that the motif co-occurrence matrix is highly sparse and skewed (a few motifs co-occur with many, while most motifs co-occur with a few), whereas the n-gram co-occurrence matrix is much denser in comparison. This comparison implies that the compositional rules for building a motif compound are highly structured. It’s worth noting that top co-occurring motifs have mixed lengths ranging from 1 to 5. Such mixed-length compositional structures are completely missing in our n-gram analysis, and this could contribute to the inefficiency of an n-gram description.

##### 3.5 Multi-layered compositional structures constrain the good solution space

To conclude this section, we take a closer look at what structures constrain the size of good program space. A full analysis of motif statistics is beyond the scope of this paper, but we hope to provide the main intuition behind our observations. Below, we list some examples about the compositional properties of motifs and compounds. Note that here we use the word “compound” to broadly mean a collection of motifs rather than a narrower definition—representing a sequence—that we adopted in the main text.

###### From wiring to motif

- A set of motifs, i.e., a compound, is used as a functional fingerprint to capture the wiring of a program. Each compound consists just a few motifs for most programs (Figure S5c).
- An identical compound can represent many structurally different programs—e.g., 130 good programs can be described as (*W*, *l*), which is an identical description of the 2-state WSLG program.

###### From motif and compound

- Not just motifs, but many compounds are themselves compositional too—e.g., a compound (*W, l*) while representing a program already can be further combined with 23 other motifs to describe another good program (see Figure S5f motif enumeration for examples)

###### From compound to sequence

- An identical sequence can be described by different compounds—e.g., a sequence *“LlllWlLll”* (which exists in 140 good programs) can be described in 12 different ways:

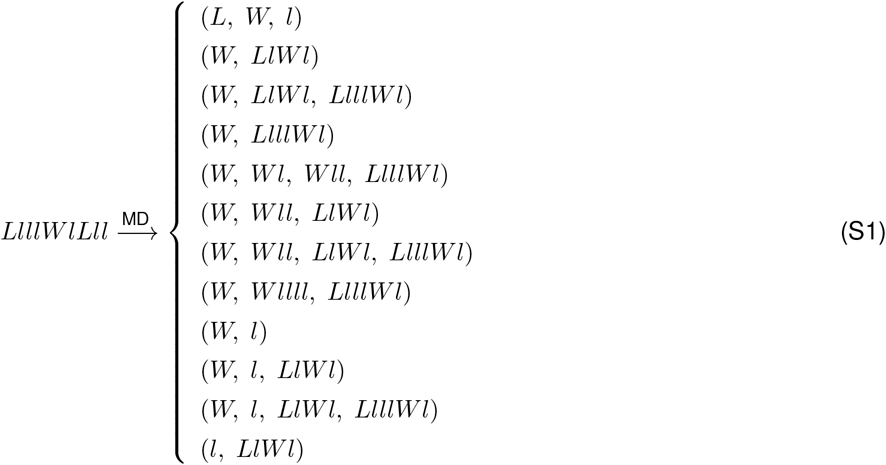

Taken together, one can see how these multi-layered compositional structures constrain the space of good solutions. In the next section, we provide a potentially power approach for exploring a much larger solution space using a constrained enumeration that leverages this compositional structure.

#### 4 Efficient search via Motif Enumeration

In the section above, we illustrated how to use motif decomposition as a compact description of program behaviors, and we showed how it helps us understand the multi-layered nature of task sloppiness. In this final section, we sketch out how motifs could be used to construct another class of efficient search algorithms, beyond the evolutionary algorithm that we demonstrated in the main text (see Section 9.4 for details).

##### 4.1 A search that enumerates behaviors instead of structures

It’s worth pointing out the difference between motif enumeration and our original full program enumeration:

1. In program enumeration, one does not add any additional constraints (beyond a resource constraint that limits number of program states) that could bias the search, whereas in motif enumeration, one takes advantage of the structures of good programs to search more efficiently.
2. In program enumeration, one enumerates the wiring directly, whereas in motif enumeration, one enumerates at a level that is closer to behavior.

The latter enumeration thus requires a mapping from behavioral motifs back to the wiring of a program. In the discussion below, we assume that such a mapping exists and can be performed efficiently, and we focus instead on the diversity that emerges at the level of motifs.

##### 4.2 Leveraging known functions to constrain an enumeration could enable exploration in a much larger solution space

In the previous section, we showed that the compositional structures of motifs can be complex (e.g., there exist additional higher-order co-occurrence matrices, beyond the pairwise one that we show in Figure S5d), and thus translating those structures into an efficient search algorithm is in itself an interesting and challenging problem. Below, we use a minimal example (Figure S5f: efficient search via motif enumeration) to demonstrate how such a search could be performed in principle:

1. One could start from the WSLG program, with its motif compound (*W, l*).
2. For an individual motif, one could then list and sort the corresponding co-occurring motifs based on their frequency—e.g., *“LW*,” “*Lll*,” and *“LLW*” are the top three motifs that co-occur with “*W*.”
3. Next, one could concatenate these motifs to form new compounds—e.g., (*W, l*) + *LW* → (*W, l, LW*), etc.
4. Each new compound now represents a set of new programs (assuming the mapping from a compound to a program can be performed efficiently) that can be readily evaluated.

As shown in Figure S5f, after going through only 48 of the top co-occurring motifs, this algorithm finds all 23 existing length-3 compounds that cover 612 good programs (i.e., that cover 14% of good program space in a single step).

Note that in this example, we only construct a relatively small compound, which is likely to represent a relatively small program. One can imagine using a larger compound as a starting point, or using a more elaborate enumeration rule to output a larger compound. This approach then presents an intriguing opportunity to enumerate towards a much larger solution space (with programs that have many more states, beyond *M* = 5). The potential advantages of doing motif enumeration can be summarized as follows:

1. Since both motifs and their compositional rules are derived directly from all known good programs, motif enumeration leverages the known structures of good program space to constrain the search in an unknown space.
2. By separating the search into two steps—first, enumerating and evaluating behavioral motifs, and second, finding realizable programs for each motif compound—the search at each step could yield much smaller enumeration. This translates into a smaller computational cost or memory usage on a computer.

Taken together, motif decomposition and motif enumeration point towards a much larger and exciting playground within which a space of good solutions can be broken down, analyzed, and understood. Moreover, the novel insights gained from this exploration can then be leveraged and translated into a generative search algorithm to discover new and previously unreachable solutions, which can in turn be used to gain a deeper understanding of the space of good solutions. It remains to be seen how such a bootstrapping should best be constructed, and what level of understanding can be reached about a problem within various domains of biology, neuroscience, cognitive science, or AI research.

### Part II MATHS & CODES

#### 5 Bayesian formalism

In this section, we discuss all approaches related to the Bayesian formalism, including how one can leverage the concise description of a Bayesian agent to efficiently find a small set of good resource-limited programs, called discretized Bayesians (DBs). At the end of this section, we discuss how it is possible, from the Bayesian perspective, to have a complex nonlinear scaling between the size of the good program space and a task parameter, even when the optimal Bayesian performance only scales smoothly.

##### 5.1 Bayesian inference

###### Task structure

A behavioral task can be described by two parameterized conditional probability distributions: 1) the world dynamics, and 2) the reward delivery. We refer to these two probability distributions as the “task structure”.

**Table S4:**
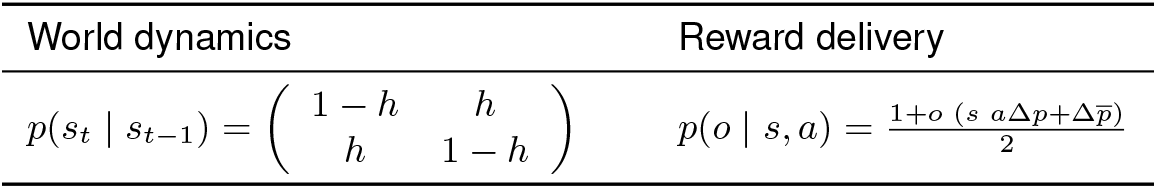
Task structures.

where all variables and parameters are listed below:

**Table S5:**
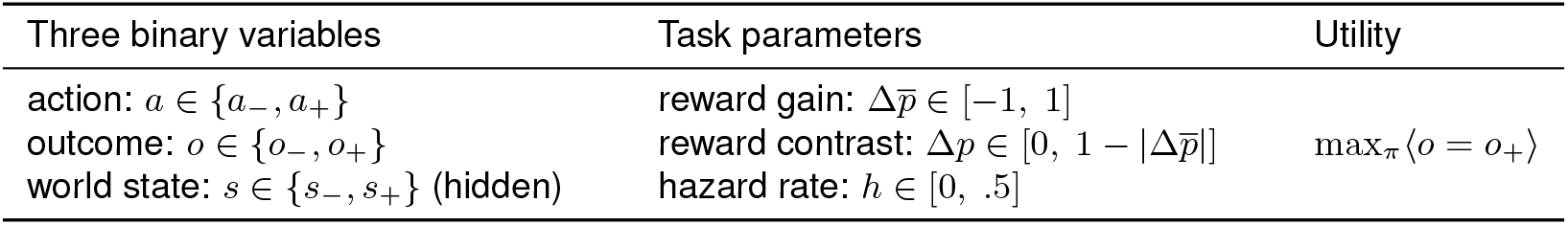
Task variables, parameters, and utility functions.

Note that from the table above, the reward gain 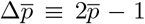 is defined as a centered and normalized baseline reward rate 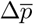

###### Using Bayesian inference to map from task structure to an iterative belief update

In a nutshell, a single step in Bayesian inference updates one’s belief about the world by incorporating a new piece of evidence. This statement can be described as follows:

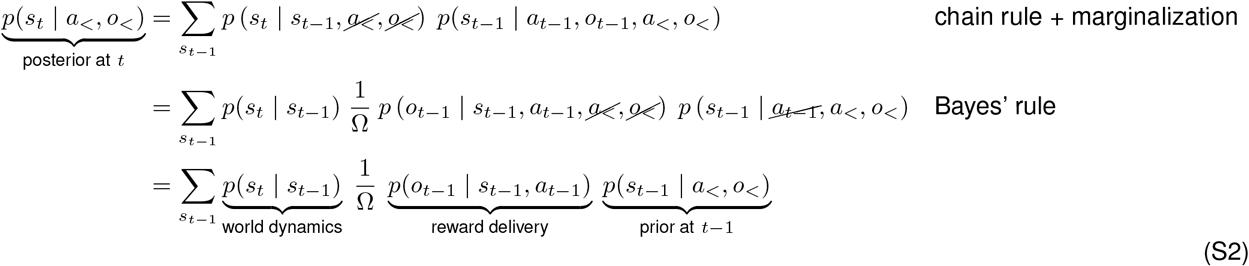

Since 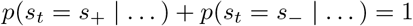, one only needs to keep track of a one-dimensional *belief value*:

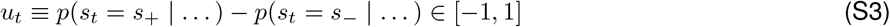

Substituting the task structure into Equation (S2) yields the following iterative belief update:

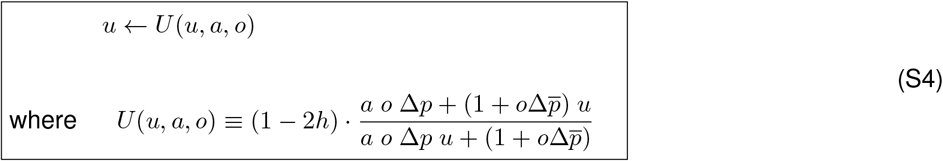

Note that in the above expressions, we use the following simplified notation for all binary variables:

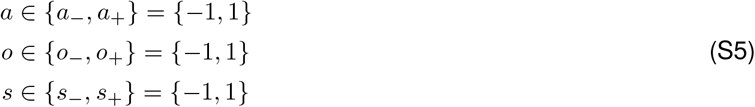

##### 5.2 Bayesian RL problem

A Bayesian RL problem is a factorization of a behavioral task into the two separate problems of 1) deriving an optimal inference and 2) finding an optimal policy. The former can be done using Bayesian formalism, as discussed above. The latter requires optimization. Fortunately, this problem is equivalent to a standard RL problem with a fully observable MDP over (infinite) belief states (these belief states can be thought of as a fine-grained discretization of the belief value in Equation (S3)). With this equivalence, a standard RL algorithm such as value iteration can be used to efficiently find the optimal policy.

###### Mapping the task to an RL problem

Given knowledge of the task, a Bayesian agent uses Bayesian inference, via Equation (S4), to recursively update its belief:

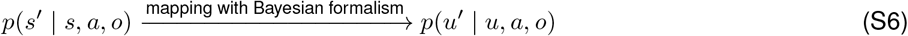

The remaining problem is to find a policy that optimizes cumulative reward:

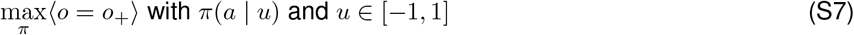

Equation (S6) and Equation (S7) can be then mapped to a standard RL problem with a fully-observable state space *u*:

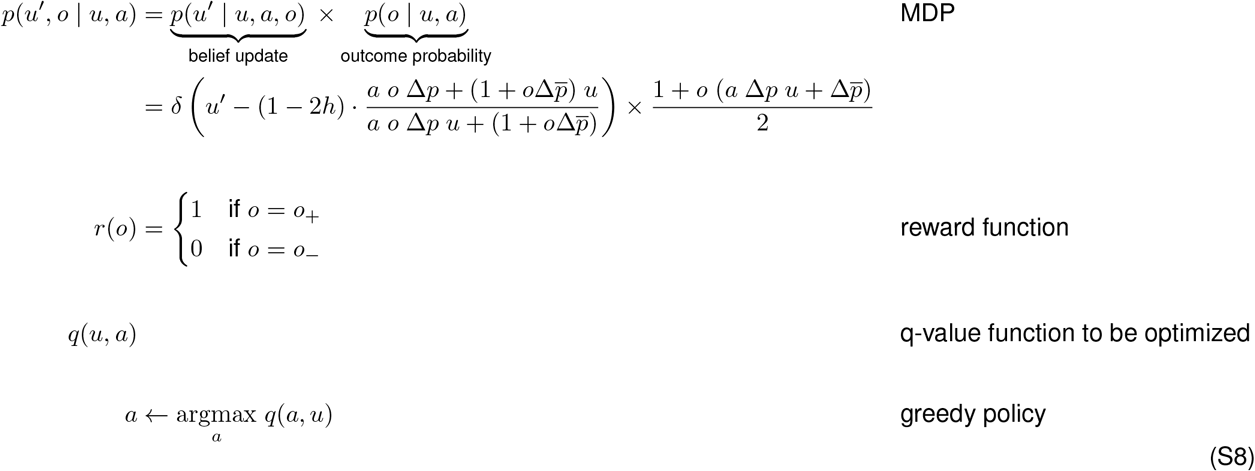

where the outcome probability can be derived from the original *reward delivery* in the task structure:

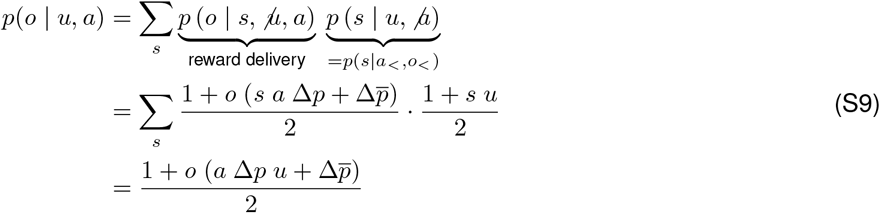

###### Value Iteration

Practically speaking, in order to use RL to derive an optimal policy, one needs to discretize a continuous belief value into finite number of belief states. Here, we use 200 non-overlapping, equally-sized bins to tile the belief space: *u* ∈ [−1, 1]. Since our problem is relatively small, it is efficient to use value iteration to optimize the policy:

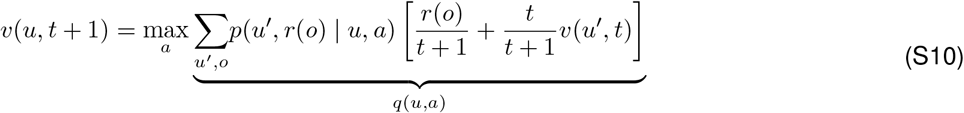

Note that here, we modify standard value iteration [43] with a built-in running average of reward over an infinite horizon. The optimal policy is then found as follows:

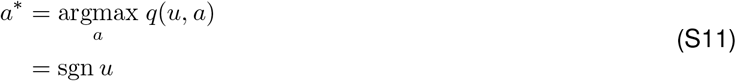

This optimal policy corresponds to a naive reward-seeking policy:

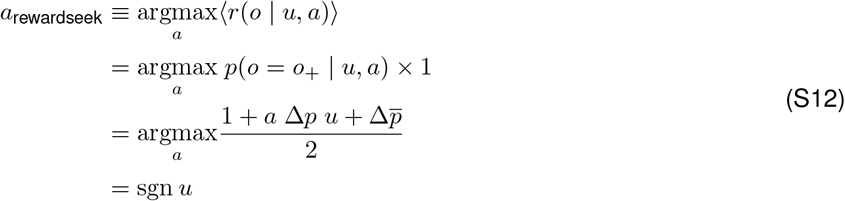

Given that the optimal policy does not change with task parameters, the particular two-armed bandit task that we study here is relatively simple from the Bayesian perspective. It is for this reason that we were surprised to find that modifying the Bayesian inference machinery can induce such complexity in the observed solution space.

##### 5.3 Discretizating the optimal Bayesian agent (DB)

The point of discretizing a Bayesian agent is to 1) qualitatively capture Bayesian behaviors with finite resources, and 2) derive an efficient search algorithm. With these goals in mind, one should avoid including excessive details in their algorithm that might 1) dilute the essence of the Bayesian computation, and 2) cover the full program space extensively (opposite of being efficient).

To preserve the essence of the Bayesian computation, we directly discretized the one-dimensional belief value into *M* ∈ {2, 3, 4, 5} equal bins, and we derived the finite-state transition matrix by finding the discrete state *m* that is closest to a transitioned belief value:

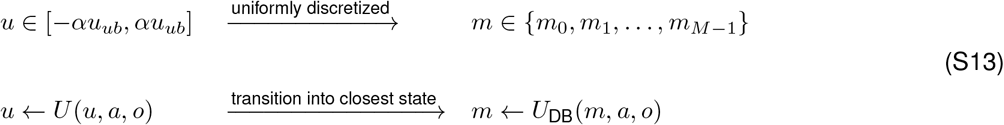

where *α* ∈ (0, 1] is a *sweeping parameter* to control the full range of discretization, and

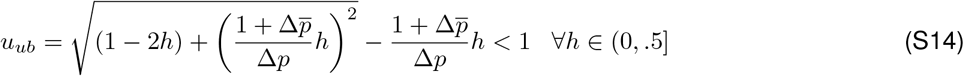

is the belief fixed-point upon continual winning (see Section 5.5 for derivation).

The last step in constructing DBs is to attach a deterministic action to each state *m* (note that the optimal policy found via value iteration is deterministic). Here, we simply use the optimal (reward-seeking) policy:

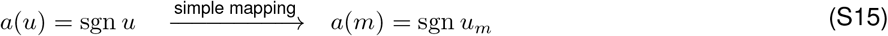

where *u_m_* is the corresponding belief value of the discrete state *m*. This discretization process generates a handful of DBs that are then filtered by a set of “rule-out rules” that eliminate invalid DBs (see Section 7.2 for details). Before evaluating these valid DBs, we decided to include a set of neighboring DBs that result from ambiguous transitions (explained in more detail below).

##### 5.4 Constrained enumeration for discretized Bayesians (DBs)

From the above uniform discretization, one note that the transitioned state is indecisive if the transitioned belief value happens to fall in the middle of two discrete states. Simply picking the nearest state may result some unwanted bias in the resultant DBs. To counter that, we simply consider all combination of nearest and next nearest states pairs. For example, consider enumerating two pairs in a three-state DB:

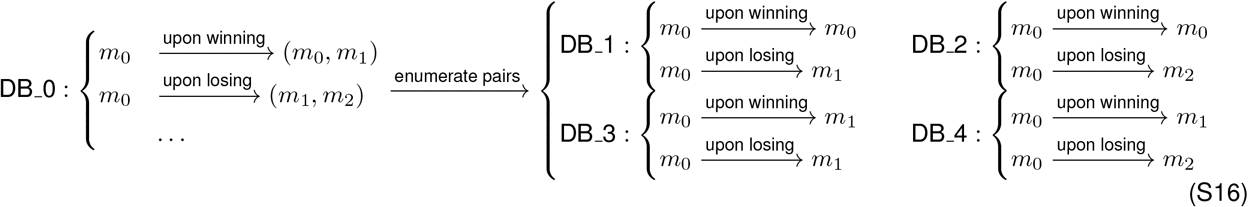

From the above enumeration, more DBs are found (after passing rule-out rules). And note that DB enumeration is a very constrained one for all the variants derived from the original discretization still keep the bi-directional integration feature (the inherent node ordering for a monotonic belief update upon consecutive winning or losing) of the original Bayesian inference.

##### 5.5 Fixed point analysis of the optimal Bayesian agent

The aim of this fixed point analysis is to categorize the behaviors of an optimal Bayesian agent in different regions of the task space. Here, we sweep hazard rate while fixing reward gain and reward contrast:

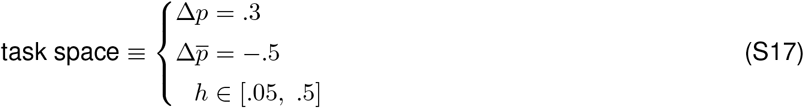

###### Five special belief values *u*

There are five special belief values that will help us to categorize Bayesian behaviors:

**Table S6:** Definitions of five special belief values.

| definition | solution | name |
| --- | --- | --- |
| $U(u_{ub}, a_+, o_+) = u_{ub}$ | $u_{ub} = \sqrt{(1-2h) + \left(\frac{1+\Delta\bar{p}}{\Delta p}h\right)^2} - \frac{1+\Delta\bar{p}}{\Delta p}h$ | belief upper bound |
| $U(u_{cr}, a_+, o_-) = 0$ | $u_{cr} = \frac{\Delta p}{1-\Delta\bar{p}}$ | critical belief |
| $u_{ll} \equiv U(u_{ub}, a_+, o_-)$ | $u_{ll} = (1-2h) \cdot \frac{-\Delta p + (1-\Delta\bar{p}) u_{ub}}{-\Delta p u_{ub} + (1-\Delta\bar{p})}$ | belief loss upon losing |
| $u_{gw} \equiv U(0, a_+, o_+)$ | $u_{gw} = (1-2h) \cdot \frac{\Delta p}{1+\Delta\bar{p}}$ | belief gain upon winning |
| $u_{gl} \equiv U(0, a_-, o_-)$ | $u_{gl} = (1-2h) \cdot \frac{\Delta p}{1-\Delta\bar{p}}$ | belief gain upon losing |

where *U* is the belief update function Equation (S4):

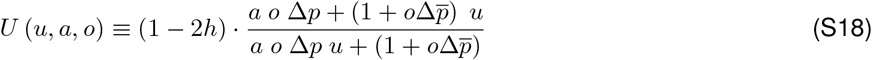

The belief upper bound *u_ub_* is defined as a fixed point upon winning beyond which no belief value is allowed at infinite horizon, *t* → ∞. The critical belief *u_cr_* got its name because the action taken by opt-B switches from lose-stay to lose-go upon losing:

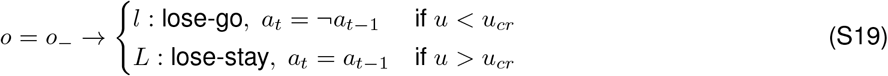

The condition for a high-likelihood of lose-stay can be translated as 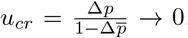. In other words, it is when a losing outcome provides little information about the current world state, i.e., 1) when reward gain is negative 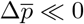 so that losing is expected; or 2) when reward contrast is low Δ*p* ≈ 0 so that neither losing nor winning can be very informative. As for the roles of the rest of the three belief values will become clear in the discussion below.

###### Four critical hazard rates

*h* The four critical hazard rates are defined as the cross over between any of the two special belief values defined above:

**Table S7:** Definitions of four critical hazard rates.

| definition | solution | value |
| --- | --- | --- |
| $u_{ll}(h) = u_{gw}(h)$ | $h_{c0} = \frac{1}{2} + \frac{1-\Delta\bar{p}}{1+\Delta p^2-\Delta\bar{p}^2} - \frac{4}{\Delta p^2+(3-\Delta\bar{p})(1+\Delta\bar{p})}$ | .112 |
| $u_{ll}(h) = u_{cr}(h)$ | $h_{c1} = \dots$ | .203<br>(analytical expression in Box S1 below) |
| $u_{gw}(h) = u_{cr}(h)$ | $h_{c2} = \frac{-\Delta\bar{p}}{1-\Delta\bar{p}}$ | 1/3 |
| $u_{ub}(h) = u_{cr}(h)$ | $h_{c3} = \frac{1-\Delta\bar{p}}{4} - \frac{\Delta p^2}{4(1-\Delta\bar{p})}$ | .36 |

**Figure S6:**
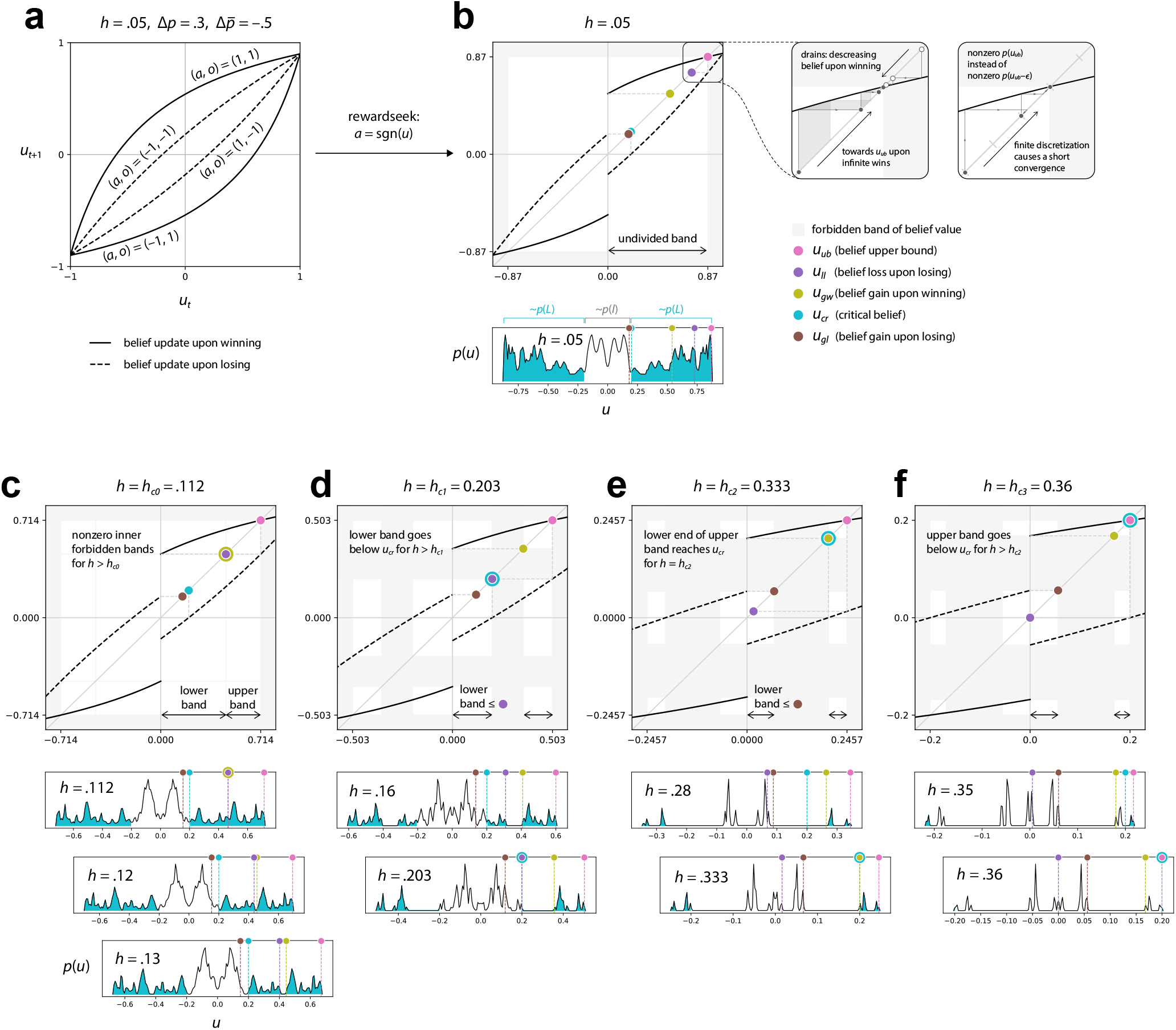
Fixed point analysis on optimal Bayesian. **(a)** Belief update derived from Bayesian formalism. The four monotonic curves specify four different action-outcome pair respectively. **(b)** Belief update under the reward-seeking (optimal) policy. The shaded region marks forbidden belief values. Five colored dots shows the five special belief values at a given hazard rate. How these belief values change positions relative to each other gives rise to a rich Bayesian behaviors undergoing a change in hazard rate. The right inserts explain how the probability of belief upper bound *u_ub_* can be overestimated. **(c-f)** The evolution of belief bands and belief distribution w.r.t changing relative positions of the five special belief values under a increasing hazard rate.

###### Box S1 Analytical expression of *h*_*c*1_

h_c1 = (1/(12*(dpm - 1)**3))*(2*(dpm - 1)*(dp**2 + 3*(dpm - 1)**2) + 2**(1/3)*(dp**6*(9*dpm - 5)*(dpm - 1)**3 - 18*dp**4*(dpm - 1)**6 + 9*dp**2*(dpm - 5)*(dpm - 1)**7 + 3*3**.5*((dpm - 1)**8*(-(dp**2 - (dpm - 1)**2)**2)*(dp**8*(2*dpm - 1) - 2*dp**6*(dpm - 1)**2*(4*dpm - 3) + dp**4*(dpm - 1)**2*(12*dpm**3 - 23*dpm**2 + 50*dpm - 23) - 8*dp**2*(dpm - 2)*(dpm - 1)**4*(dpm + 1)**2 + 2*(dpm - 1)**6*(dpm + 1)**3))**.5)**(1/3) + (2**(2/3)*(dp**4*(3*dpm - 1)*(dpm - 1)**2 - 6*dp**2*(dpm - 1)**5 + 3*(dpm + 1)*(dpm - 1)**6))/(dp**6*(9*dpm - 5)*(dpm - 1)**3 - 18*dp**4*(dpm - 1)**6 + 9*dp**2*(dpm - 5)*(dpm - 1)**7 + 3*3**.5*((dpm - 1)**8*(-(dp**2 - (dpm - 1)**2)**2)*(dp**8*(2*dpm - 1) - 2*dp**6*(dpm - 1)**2*(4*dpm - 3) + dp**4*(dpm - 1)**2*(12*dpm**3 - 23*dpm**2 + 50*dpm - 23) - 8*dp**2*(dpm - 2)*(dpm - 1)**4*(dpm + 1)**2 + 2*(dpm - 1)**6*(dpm + 1)**3))**.5)**(1/3)),

where 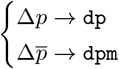

This expression is acquired by using `Solve` and `Simplify` functions in Mathematica [44]

#### 6 Sloppiness and the collapse of the good program space

##### 6.1 Sloppiness as a property of task and solution spaces

The notion of task and function sloppiness—positively associated with the the number of good solutions—is discussed briefly in Section 2. In this section, we take a closer look at how the this number scales with 1) different task structures and 2) different constraints on solution space:

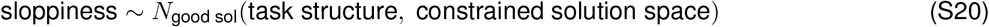

There are many different ways to formalize these two contributors. To formalize the task structure, we compute the behavioral difference between an optimal Bayesian (opt-B) agent and a win-stay-lose-go (WSLG) program. For Bayesian formalism essentially map the task structure as a whole into an iterative belief update. As to constrain the solution space, we first use discretized Bayesians (DBs) to study how many resource are required to approach opt-B performance. In so doing, we gain some insights on what fraction of the full solution space could turn out to be good—i.e., the size of good program space. And then we evaluate all programs on all tasks to see how well the good program space as a whole captures opt-B behaviors and how much freedom (sloppiness) these good programs are allowed to deviate from them. Formally, we try to show:

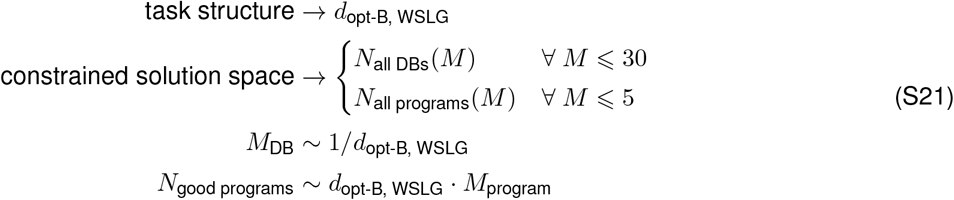

where *d* denotes behavioral difference and *M* denotes number of states in a DB or program. Note that the above formalization is only meant to describe an approximate association instead of a precise linear or inverse-linear relationship. Here we would like to capture that 1) many states *M*_DB_ are needed for a DB to catch up opt-B performance when *d*_opt-B,WSLG_ is small, and 2) the size of good program space grows with either an increasing *d*_opt-B, WSLG_ or *M*_program_.

##### 6.2 The many stages of behavioral convergence

With all special belief values and critical hazard rate defined, one can take a closer look on their consequences regarding the behavioral convergence between optimal Bayesian (opt-B) and win-stay-lose-go (WSLG) program when gradually increasing hazard rate from *h* = 0 to .5:

1. starting from a small hazard rate, all five belief values are well spread out, and the allowable belief values form a single continuous band;
2. above *h*_*c*0_, a forbidden band of belief value emerge because the only those belief value with *u* > *u_ub_* (not allowable) can transition into this band gap upon losing;
3. while the band gap grows, a growing portion of the lower band goes below critical belief *u_cr_*, and it causes a gradual decrease of lose-stay probability *p*(*L*);
4. above *h*_*c*1_, the lower band is fully submerged under *u_cr_*;
5. while *u_cr_* falls between lower and upper band, the ratio *p*(*L*)/*p*(*l*) remain approximately constant;
6. as the upper band reaches *u_cr_* at *h*_*c*2_ and starts to go below *u_cr_*, lose-stay probability undergoes another drop;
7. above *h*_*c*3_, lose-stay probability drops to zero, and opt-B converges to WSLG in all aspects.

**Figure S7:**
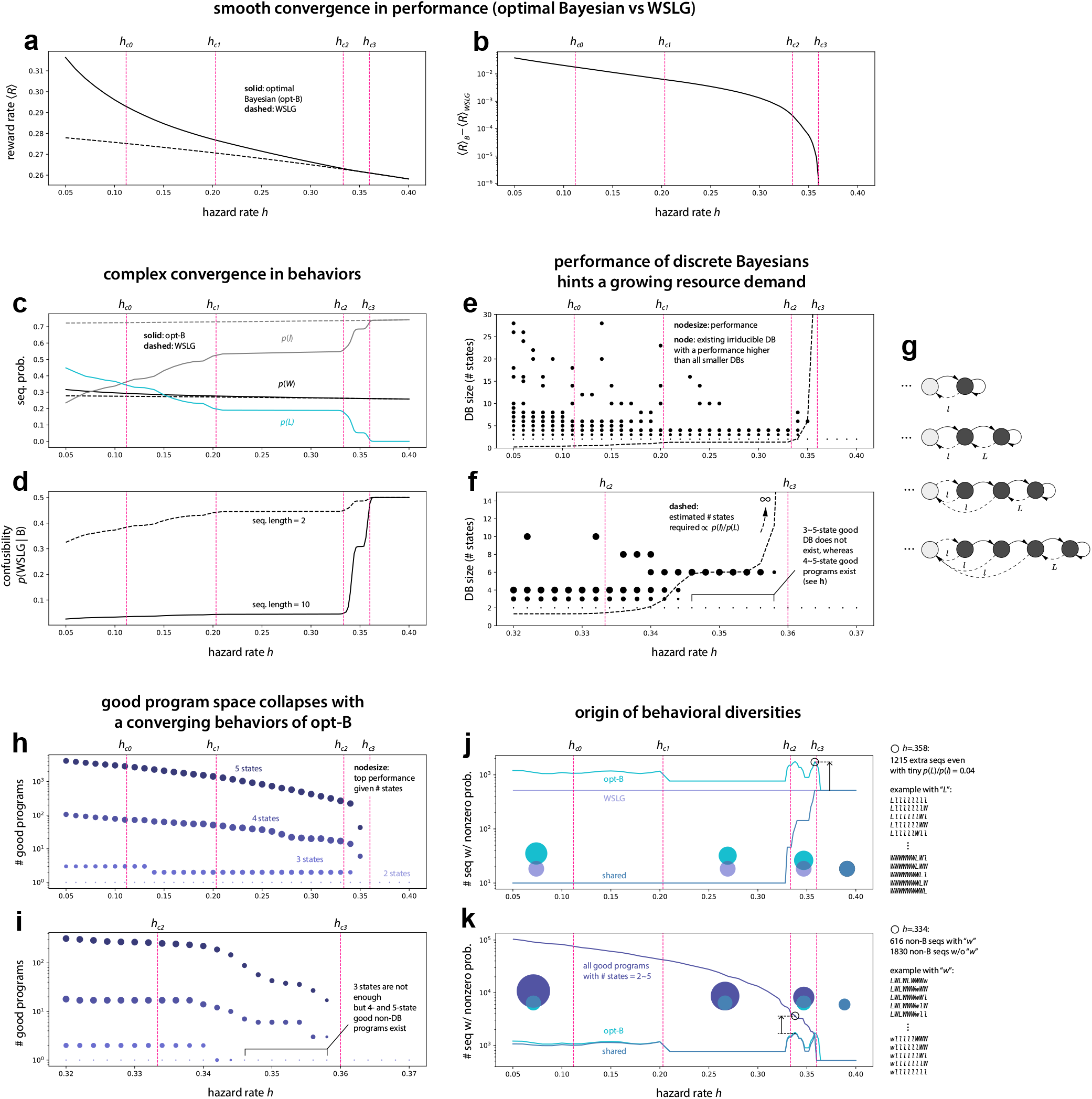
A task-related sloppiness determined by structures of the task and constraints on solution space. **(a)** Converging performance between optimal Bayesian (opt-B) agent and WSLG program under an increasing hazard rate. **(b)** Performance difference rapid drop to zero as *h* → *h*_*c*3_. Note that this curve is smooth without showing any sign of a complex Behavioral changes in opt-B under a changing hazard rate. **(c)** First-order sequence statistics as a function of hazard rate. Note that even with this lowest-order behavior, opt-B shows complex changes with an increasing hazard rate. Specifically the ratio *p*(*L*)/*p*(*l*) changes in a somehow unpredictable manner. **(d)** The corresponding confusion probability computed with short and long sequences. For long behavioral sequences, opt-B behavior clearly undergoes a phase transition near *h*_*c*3_ and *h*_*c*4_. **(e-f)** At each hazard rate, a DB that can make an incremental improvement from the closest smaller DB is marked. As hazard rate increases, it becomes harder and harder to find the next DB that can incrementally improve towards opt-B with merely an addition of a few resources (program states). The dashed line is a simple fit indicating the growing demands of resources is proportional to the delicate ratio *p*(*l*)/*p*(*L*) a DB has to achieve. **(g)** A larger DB has a better chance in achieving a skewed ratio of *p*(*L*)/*p*(*l*). **(h-i)** A collapse of the good program space as a function of hazard rate and program size. The size of the good program space—hence sloppiness—follows closely with a changing behavior of opt-B. **(j)** An explosion in behavioral deviation between opt-B and WSLG program can be understood as how number of permissible sequences grows combinatorially large even with a small increase in *p*(*L*)/*p*(*l*). **(k)** An explosion in behavioral diversity between opt-B and the whole good program space can be understood similarly as how a small deviation from opt-B in lower-order statistics can lead to combinatorially many permissible sequences.

**Table S8:** Evolution of belief bands upon increasing hazard rate.

| task range | belief bands | band evolution |
| --- | --- | --- |
| $h \in [0, h_{c0})$ | $u \in [-u_{ub}, u_{ub}]$ | singular continuous band |
| $h \in [h_{c0}, h_{c1})$ | $u \in [-u_{ub}, u_{ub}]$ | onset of forbidden band at $h_{c0}$<br>→ growing band gap with increasing $h$ |
| $h \in [h_{c1}, h_{c2})$ | $u \in [-u_{ub}, -u_{ll}], [-u_{ll}, u_{ll}], [u_{ll}, u_{ub}]$ if $u_{ll} > u_{gl}$<br>$u \in [-u_{ub}, -u_{gl}], [-u_{gl}, u_{gl}], [u_{gl}, u_{ub}]$ if $u_{ll} < u_{gl}$ | lower band submerged under $u_{cr}$ at $h_{c1}$<br>→ $u_{cr}$ falls within band gap |
| $h \in [h_{c2}, h_{c3})$ | $u \in [-u_{ub}, -u_{gl}], [-u_{gl}, u_{gl}], [u_{gl}, u_{ub}]$ | upper band reaches $u_{cr}$ at $h_{c2}$<br>→ part of upper band goes below $u_{cr}$ |
| $h \in [h_{c3}, .5]$ | $u \in [-u_{ub}, -u_{gl}], [-u_{gl}, u_{gl}], [u_{gl}, u_{ub}]$ | upper band submerged under $u_{cr}$ at $h_{c3}$ |

It is worth to point out that opt-B, with its reward-seeking policy, never switch action upon winning, i.e., *p*(*w*) = 0 where *w* denotes “win-go” action. A useful coarse-grained description for opt-B behavior can therefore be a vector of first-order sequence distribution:

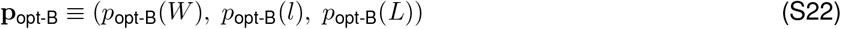

Similarly, WSLG and thus the difference between opt-B and WSLG can be defined as

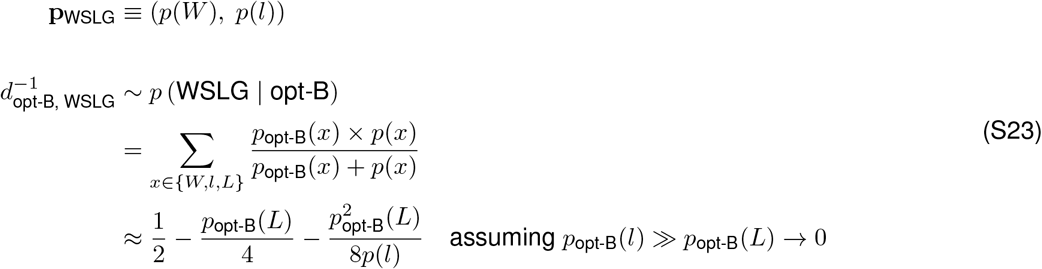

Note that we choose to use the confusion probability (confusion matrix is discussed in Section 10.1) to capture the behavioral difference:

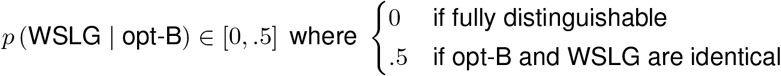

In Figure S7d, we plot *p* (WSLG | opt-B) as a function of hazard rate. One can see how opt-B converges to WSLG as *h* → *h*_*c*3_ = .36. Crucially, before the convergence, the distinction between opt-B and WSLG becomes sharp if one use longer sequences (length=10) to characterize them. This fast increase of behavioral difference with increasing sequence length is the main cause of high degree of sloppiness—for many other programs that generate various sets of sequences lying between WSLG and opt-B are allowed to exist with good performance. This point is discussed below in more details.

###### Box S2 Minor numerical error as a result of a finite discretization

In Figure S7b zoom-in insert, we illustrate how finite discretization can affect the convergence of belief distribution near *u_ub_*. For such a convergence could happen much faster with coarser bins which results an exaggerated probability at *u_ub_* in stead of peaking at *u*_peak|inf. discr_ = *u_ub_* – *ϵ* with vanishing probability at *u_ub_*: *p*(*u_ub_*) = 0.

##### 6.3 Increasingly more resources are required to marginally improve performance at high hazard rates

The stiffness of the solution space near the last critical hazard rate, *h* ≈ *h*_*c*3_ = .36, can be understood by estimating how much extra resource required to further improve from WSLG. To do so, we evaluate all 528 discretized Bayesians (DBs) with their size ranging from *M* = 2 to 30 states. These DBs are found using two different discretization processes (see Box S3) across various hazard rate *h* ∈ [.05, .4] within which the smallest DB is simply WSLG. In Figure S7(e,f), one can see that a 3-state DB can further improve upon WSLG for a lower hazard rate *h* < *h*_*c*2_ = .33. This condition starts to shift when *h* ≈ *h*_*c*2_ where the improvement at 3-state level decreases moving towards higher hazard rate. And for *h* ⩾ *h*_*c*3_, having 3, 4, or even 5 states fails to make any improvement.

At first glance, it seems counterintuitive that making a smaller improvement (at higher hazard rate) requires more resources than a larger improvement (at lower hazard rate). To see why, one can focus on the first order behavioral difference between opt-B and WSLG, i.e., *p_B_*(*L*). For example, at *h* = .1, the ratio between lose-stay and lose-go is *p_B_*(*L*)/*p_B_*(*l*) = 1.1 for which one can sketch a DB like Figure S7g that has a similar amount of lose-stay and lose-go edges and expect a large improvement from WSLG given that such a DB emulating opt-B behaviors much better. In contrast, for *h* = .35, one has *p_B_* (*L*)/*p_B_*(*l*) = .08 which is a rather small adjustment from WSLG. To achieve such a skewed ratio, one needs to add more states in a DB in order to dilute the proportion of lose-go edges to just the right amout for a descent improvement (if not degrading the performance by overshooting). In Figure S7f dashed line, we see that a simple estimate on how much states required for any improvement from WSLG roughly captures the results from our numerical experiment:

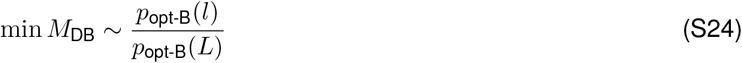

In our numerical experiment, no good DBs with more than 8 states (and less than 30 states) are found for *h* ⩾ *h*_*c*2_ = .33.

It is curious to point out that the good program space as a whole can capture nearly all opt-B behaviors as illustrated in Figure S7k (see curves opt-B vs shared); however, a significant deviation happens when the hazard rate is either very low or near the phase transition: *h* ≈ 3.5. In the case of low hazard rate, a larger DB is required to approach opt-B performance for a long integration is needed. A program space with *M* ⩽ 5 therefore does not have quite enough resources to capture all necessary Bayesian behaviors. At the other extreme with large hazard rate, a large amount of resources is again needed to generate a tiny percentage of lose-stay L behaviors, and thus the program space we have cannot mimic such a delicate opt-B behaviors for the same reason.

###### Box S3 Two discretization processes to generate a library of DBs

In Figure S6c, we note that a band gap of forbidden belief values opens around *h* = *h*_*c*0_. This band gap ends up lasting all the way to the last critical hazard rate *h*_*c*3_ with an ever increasing width. It’s therefore beneficial to consider a discretization without putting any precious resources in those belief values. To do so, we remove those band gap, concatenate the remaining belief values, and discretize them uniformly. Together with earlier version of discretization. We evaluate a library of both DBs in Figure S7(e,f).

##### 6.4 A sudden collapse of the good program space

In this section, we focus on a narrow range between two critical hazard rate, *h* ∈ [*h*_*c*2_,*h*_*c*3_], which clearly display a phase transition in opt-B behaviors. In Figure S7d, we have seen a sharp increase in confusion probability *p*(WSLG | opt-B) once *h* passes *h*_*c*2_. Correlated to this observation is a sudden collapse of the good program space—i.e., WSLG program is the only good solution left:

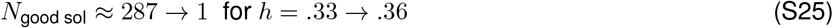

It is curious to point out that there is a very narrow plateau around *h* ≈ .35 in both *p*(WSLG | opt-B) and *N*_good sol_ (see Figure S6g for a zoom-in plot). And during that short window of increasing hazard rate, the opt-B behaviors do not change much, neither do the good program space. The cause of this transient stillness can be understood by carefully examining how the belief distribution evolves from *h* = .33 → .35 → .36 in Figure S6(e,f)).

From these results of the fixed point analysis on opt-B and how it correlates to the size of good program space, we have demonstrated that

1. a simple analytical belief update rule from Bayesian inference does not translate to a simple Bayesian behavior (see complexity of belief distribution in Figure S6(b-f)))
2. a smoothly changing task structure (e.g., hazard rate) and Bayesian performance, does not necessarily translate to a smoothly changing Bayesian behavior or a smoothly changing good program space;

###### Box S4 Enumerating only 6961 (out of 268,536) programs on all hazard rates

It turned out that, for generating Figure S7(h,i)), one does not need to evaluate the full program space again. Because the good program space at higher hazard rate is contained by the good program space at lower hazard rate (up to some numerical errors due to finite discretization of belief values). This observation is done by full enumeration at three hazard rate: *h* ∈ {.15, .25, .35}. With this observation, in principle, one can just run the 4230 good programs at *h*_min_ = .05 for all higher hazard rate. But to be on the safe side, we relax the criterion and also include programs that have performance 99 to 100% WSLG. This would expand the space from 4230 to 6961. After evaluating these 6961 programs at different hazard rate, there are only up to 3 programs that ever fall out of the original 4230 good programs.

This inclusiveness (or unchanged nature) of the good program space implies that the task is qualitatively the same even with different hazard rate. It is therefore important for us to move away from a simple parameterized task if we ever wish to study generalizability in a task space where the spaces of good solutions for these tasks overlap in some complex way.

##### 6.5 The number of permissible behaviors explodes with slight deviations from the optimal Bayesian agent

To close this section, we take a look of the origin of this sudden collapse of the good program space. More intuitively, one can view this collapse as an explosion in reverse. In Figure S7(h,i)), one can see that the sequences an opt-B can produce completely overlap with WSLG if *h* ⩾ .36 + *ϵ* (here *ϵ* ≠ 0 because of discretization errors discussed in Box S2). As one decreases the hazard rate *h* ≲ .36, the number of sequences the opt-B possess increases whereas the number of sequences shared with WSLG drops rapidly. The right text panel in Figure S7j illustrates how this can happen even with a tiny percentage change in its lowest order statistics, i.e.,

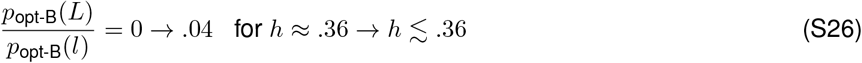

Although the amount of lose-stay *L* is small, it suddenly generates vast amount of new combinations with the original two letters: *W, l*. This kind of sloppiness created by the combinatorial freedom also exist at the level of non-Bayesian behaviors. In Figure S7k, one can see that

1. the good program space as a whole possesses nearly all the opt-B sequences, and
2. there are far more sequences beyond opt-B behaviors.

The text panel on the right shows how a small amount of win-go *w* (something an opt-B never does) behavior creates a large number of novel sequences.

#### 7 Program enumeration

The challenge about enumeration is to identify relevant constraints one can put into the algorithm such that it can be done in a manageable way. E.g., being able to fit into available memories on a computation cluster, being fast enough to finish in hours instead of days, etc. These constraints broadly can be categorized into two groups:

1. *parametric constraints*, e.g., *resource constraint*, or number of program states in our case;
2. *non-parametric constraints*, e.g., *rule-out rules* that eliminate undesirable solutions. In our case, a large program that has identical behaviors as one of the smaller one is undesirable.

In this study, we choose an upper bound of 5 program states for the resources constraint. And we use a complete set of rule-out rules that ensures all valid programs are unique. In the discussion below, we show detailed steps for how such a full program enumeration can be done.

##### 7.1 Multiple tables represent a single program

In this study, a behavioral program is a deterministic Markov chain, which is a graph without any inherent notion of node ordering. However, to enumerate a graph, one has to embed it with some choice of node labeling. This is partly why the full program enumeration requires rule-out rules to eliminate repeated programs, despite the fact that they might have different representation. Below is an example that highlights two different representations of the same program:

**Table S9:** Two distinct representation of the same underlying program.

| | $m$ | $m' \mid \text{losing}$ | $m' \mid \text{winning}$ | $m$ | $m' \mid \text{losing}$ | $m' \mid \text{winning}$ |
| --- | --- | --- | --- | --- | --- | --- |
| tabular representation | $a_- \leftarrow 0$ | 1 | 1 | $a_- \leftarrow 1$ | 0 | 0 |
| | $a_+ \leftarrow 1$ | 0 | 1 | $a_+ \leftarrow 0$ | 1 | 0 |
| | | $\Downarrow$ | | | $\Downarrow$ | |
| program tuple: (outmap, inmap) | $\left( (a_-, a_+), \begin{pmatrix} 1 & 1 \\ 0 & 1 \end{pmatrix} \right)$ | | | $\left( (a_-, a_+), \begin{pmatrix} 0 & 0 \\ 1 & 0 \end{pmatrix} \right)$ | | |

Note that each *M*-state program can be represented as a tuple with an *M*-dimensional binary `outmap` vector and a *M* × 2 `inmap` array with state label *m* ∈ [0, *M* – 1]. Below we list two types of rule out rules that operates at the level of this program tuple and remove invalid ones. What remain in the list is then a complete set of unique programs—i.e., no two graphs are the same.

##### 7.2 Task-independent rule-out rules

The first set of three rule-out rules are based on some generic features of Markov chain itself which are independent of our task choice. These three rules attempt to filter out:

1. ill-behave Markov chains, and
2. repeated programs through a node permutation.

**Table S10:** Task-independent rule-out rules.

| rule | note |
| --- | --- |
| Rule 1) Remove identical graphs under node permutation | The most obvious rule-out rule recognizes that the underlying graph structure of a program is unchanged regardless how one index the nodes to construct the corresponding tabular representation. |
| Rule 2) Remove reducible Markov chains | Rule 2 & 3 keeps only well-behaved Markov chains. A reducible Markov chains contains sinks—a subset of nodes that absorbs all occupancy during random walk, and/or drains—the subset of remaining nodes. |
| Rule 3) Remove periodic Markov chains | A periodic Markov chain causes a non-converging distribution of node occupancy. Rule 2 & 3 can be checked using an open python library: QuantEcon [45] |

##### 7.3 Task-dependent rule-out rules

While the above three rules generally eliminate all repeated or ill-behaved Markov chains, there are still programs turned out to be equivalent because of some symmetries or structures present in the task. In other words, even though each program might have a distinct Markov chain, two programs are considered equivalent if they generate identical action-outcome sequence in the same task.

**Table S11:** Task-dependent rule-out rules.

| rule | note |
| --- | --- |
| Rule 4) Remove identical graphs under node inversion | Since the task is symmetrical in two actions, flipping the policy: $a_{\pm} \rightarrow a_{\mp}$ would neither change the outcome trajectory nor performance. |
| Rule 5) Remove programs that contain mergers | Mergers are nodes that can be combined without changing a program's behavior. In the section below, we show how this rule can detect all types of mergers. |

##### 7.4 Algorithmic steps to rule out a merger program

Below we focus on the detailed steps in the merger rule-out rule, i.e., Rule 5 for 1) it is hard to intuit and 2) it enables a way to measure structural distance between two programs with different sizes. The logic behind checking if a program contains any mergers is to try every possible way to merge multiple groups of nodes, and see if under a particular merging, the resultant program produces that same behavior. Knowing this, the first step is to find all unique node partitions.

###### Step 1) partition *M* program states (nodes) into all possible groups

For a given number of states, say *M* = 5, one can create either 1, 2, 3, or 4 groups with their sizes:

| # groups | possible group sizes |
| --- | --- |
| 1 | (5,) |
| 2 | (4,1), (3,2) |
| 3 | (3,1,1), (2,2,1) |
| 4 | (2,1,1,1) |

For a particular group size tuple, one can generate all possible grouping for node ids, and then remove all repeated grouping. For example, the following two partitions are identical:

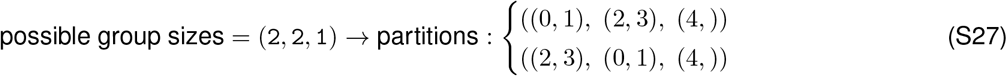

###### Step 2) remove partitions that contain different actions in any groups

If a group within a partition contains multiple nodes that do not label the same action, these node cannot be merged. For example, given an `outmap=(-,+,+,+,+)`, a `partition=((0, 1), (2, 3), (4,))` can be relabeled as `((-,+), (+,+), (+,))` and since the first group contains different actions, this partition is no merger.

###### Step 3) relabel node ids with group ids, reduce inmap accordingly

In this step, we create a mapping from the original node ids, say `m=(1, 2, 3, 4, 5)` with a smaller set of group ids, say `g=(A,B,C)`. For example,

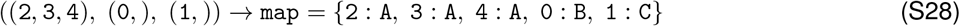

Then we replace the node ids in transition matrix—i.e., `inmap` accordingly:

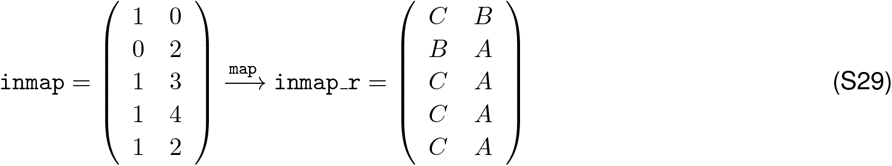

###### Check if each group in inmap has consistent winning & losing transitions

From the above, one notes that for group `(2, 3, 4)`, the first (losing) column contains only a single transition to C; similarly, for the second (winning) column, it contains only a transition to `A`. This implies that this program is indeed a merger program, and can be reduced to a smaller program under this particular partition.

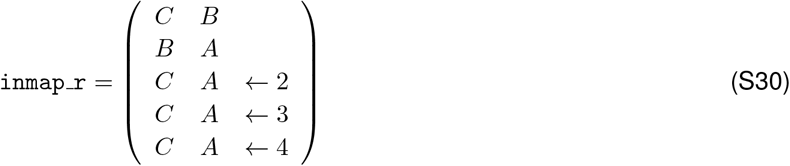

##### 7.5 Optimizing algorithmic steps for a full enumeration

With all rule-out rules formalized. One can combine them in a most efficient order for optimizing enumeration speed, memory cost, etc. on cluster computers. Below, we show a particular way to achieve such:

###### Step 1) enumerate outmap with standardized ordering

If two tabular representation can ever coincide after node permutation, they would have the same `outmap`. Given this, one can fixed `outmap` to be any of the following vectors throughout enumeration and only vary `inmap`.

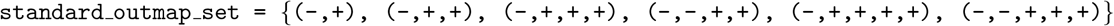

This set always have more or equal number of *a*_+_ than *a*_−_, and all *a*_−_ gather to the left side.

###### Step 2) enumerate inmap

For a given program size, say *M* = 4, there could be *m* ∈ [0, 1, 2, 3] for each entry, and thus *M*^2*M*^ different `inmaps` for a fixed *M*. Each `inmap` later will pair with all compatible `outmap` to generate a program tuple: `program=(outmap, inmap)`. The total number of program tuples scales with *M* as follow:

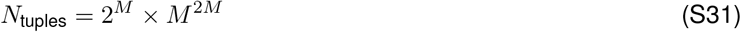

###### Step 3) remove inmaps with sinks and drains

This step is the fastest and does not involve `outmap`, so we use it as the first filter to eliminate majority of invalid programs.

###### Step 4) remove mergers

For each program tuple `program`, we use Rule 5 discussed above to eliminate all merger programs.

###### Step 5) remove repeated programs

This step is generally much slower given that it require cross checking between a given program and all programs in the list of being valid. To do so, we loop over all generate all existing program tuples `(outmap, inmap)`. For a given tuple, we first generate all permuted version that does not change the `outmap` (i.e., node permutation that would output the same or mirror version of the original `outmap`). Then we check if any permuted tuple already exist in the list of valid program. If not, we append this tuple to the list.

###### Step 6) remove reducible and periodic inmap

The reason we put this rule at the bottom because checking these feature in Markov chain with QuantEcon is slow compared to the above. And it removes much fewer programs as well.

With the full enumeration algorithm in place, we can enumerate up to *M* = 5 with ease (a bit of a stretch for *M* = 6, and not possible for *M* = 7). Below, we show some statistics to summarize the result:

**Table S12:** Number of program tuples enumerated vs number of unique program.

| $M$ | # unique tuples | # unique programs |
| --- | --- | --- |
| 2 | 64 | 5 |
| 3 | 5832 | 124 |
| 4 | 1,048,576 | 4979 |
| 5 | 312,500,000 | 263,428 |

From this result, one gets a sense about how full enumeration blows up as *M* grows. It is therefore important to constrain an enumeration either through an evolutionary algorithm (discussed in Section 9.4 below) or through leveraging some structures that could be beneficial to task (see discussion in Section 4.2).

#### 8 Program evaluation

##### 8.1 Belief Distribution Propagation (BDP)

The most common way to evaluate a Markov model is to use Monte Carlo simulation, in which one needs a long trajectories for the statistic (e.g., state occupancy) to converge. In our case, we can bypass this issue by propagating an entire belief distribution over time. The convergence is order-of-magnitude faster at the cost of needing to store the full distribution in memory rather than a single belief value over the course of iteration.

###### BDP for the optimal Bayesian agent

To evaluate the optimal Bayesian agent, we start from an initial uniform distribution across belief values *u* ∈ [−1, 1] discretized into 200 bins. In the first iteration *t* = 0, we propagate the probability of each belief value based on all four action-outcome pairs; this then generates four new belief values at *t* = 1 with corresponding probabilities:

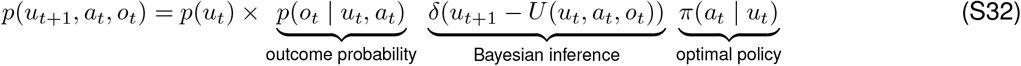

where *U*(*u, a, o*) is the belief update we derived in Equation (S4), *p*(*o* | *u*, *a*) is the outcome probability derived in Equation (S9), and *π* (*a* | *u*) is the optimal policy derived in Equation (S11). The updated belief distribution *p*(*u*_*t*+1_) is then simply the above summing over all action-outcome pairs and integrating over previous belief distribution *p*(*u_t_*):

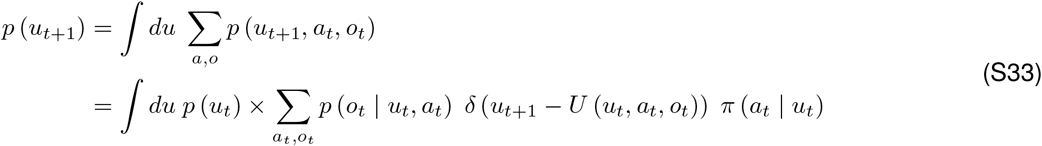

###### BDP for a small program

BDP for small program follows the same logic as the above. The only extension is to have a joint distribution of both program states and belief states. The update rule is therefore:

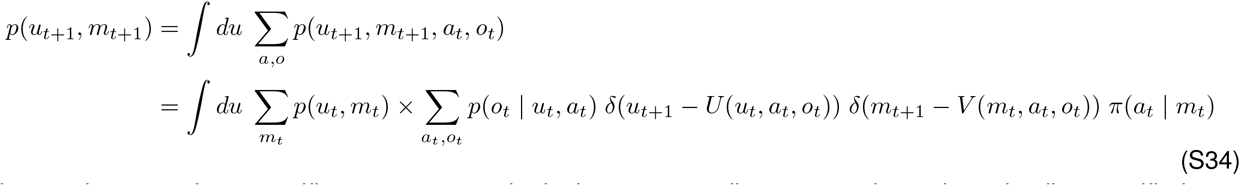

where *m* is a state for a specific program, *V* and *π* is the corresponding state update rule and policy specified as a deterministic Markov chain (see Section 7 for details). Note that what BDP does is equivalent to find the eigenvector (belief distribution) of the top eigenvalue in a joint transition matrix of a given Markov chain. The eigenvector approach in fact run faster than our BDP, but it is harder to scale up to a larger joint distribution like the one we used in SDP discussed below.

##### 8.2 From belief distribution to performance

Having found the converged belief distribution *p*(*u*) or *p*(*u, m*) for an agent, it is straightforward to compute the corresponding reward rate:

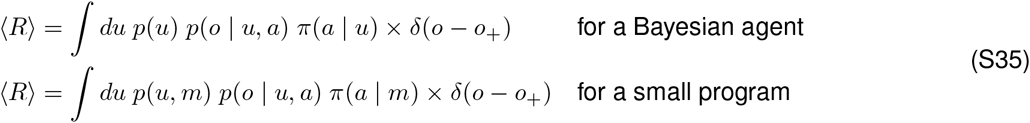

##### 8.3 Sequence Distribution Propagation (SDP)

The goal of SDP is to iteratively build a sequence distribution by updating belief distribution while increasing depth. Unlike BDP, which only keeps a distribution of at the last time step, SDP keeps upto *l*-steps to correctly generate a sequence distribution with length *l*.

###### Two auxiliary transition matrices

To simplify the analytical expression for the update rules, we defined two auxiliary transition matrices *A* and *B*:

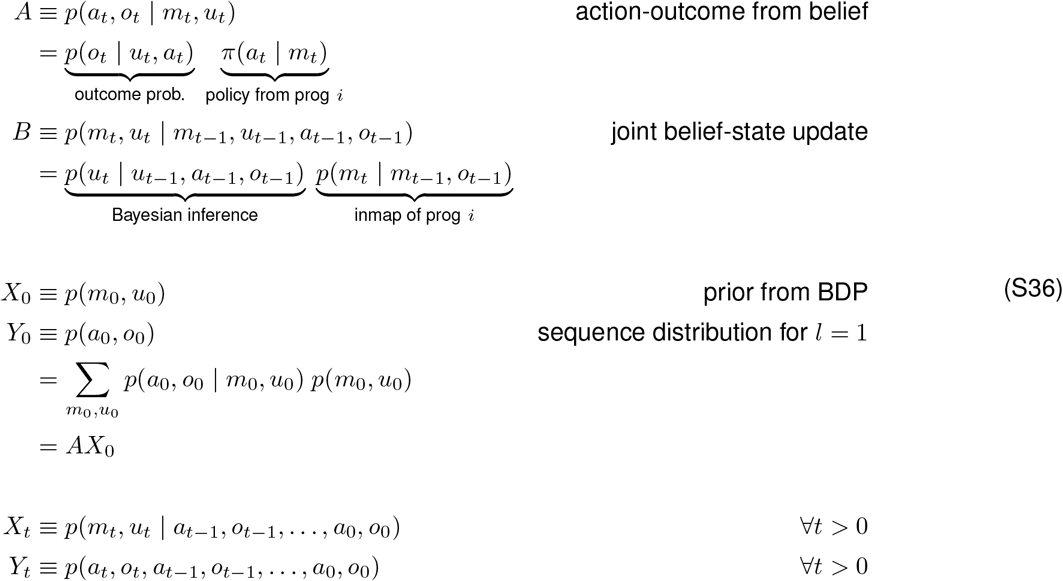

###### Iterative update rules

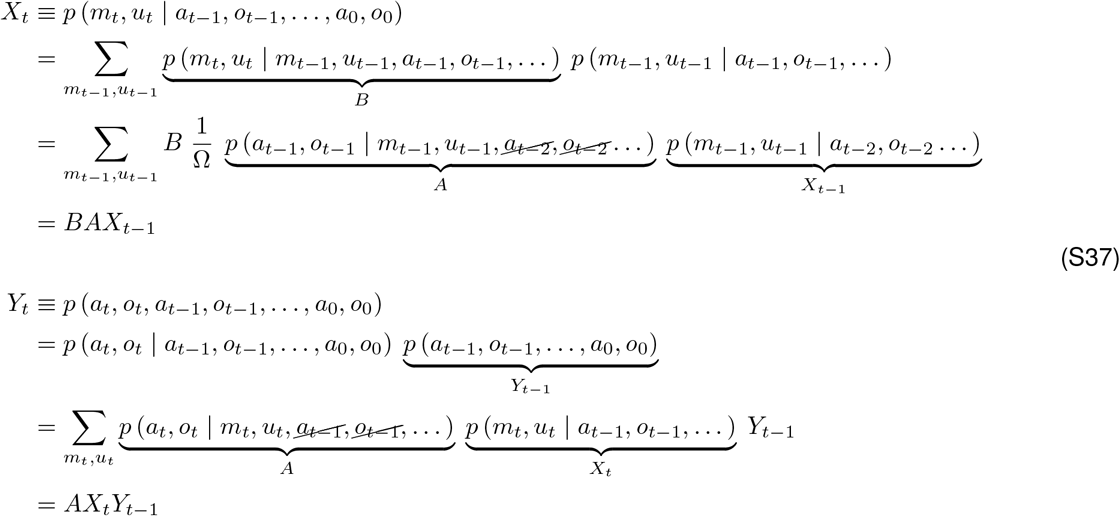

###### SDP for Bayesian agent

It’s straightforward to simply the above update rule to apply to Bayesian agent:

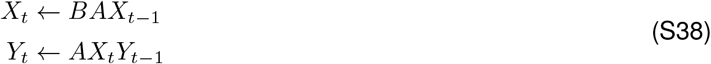

where

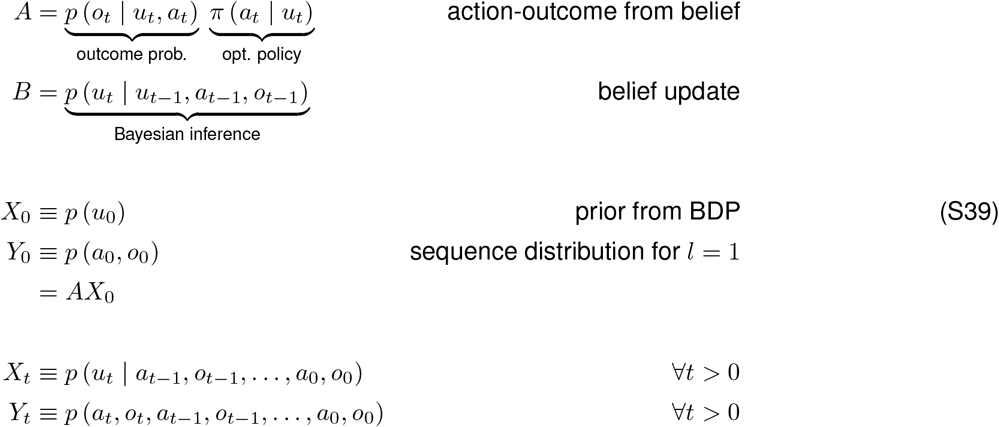

In this study, we propagate a sequence distribution to *l*_max_ = 10, which is a distribution of 4^10^ = 1,048,576 distinct sequences.

#### 9 Extracting relational structures from a program space

##### 9.1 Structural distance

###### When two programs have the same size

It is relatively easy to define structural-distance for two same-size programs. One simply chooses one of the two to have a fixed tabular representation (a fixed node ordering), while permuting the nodes of the other program until the difference between two tabular representations is minimized. Here we use binary difference—i.e.,

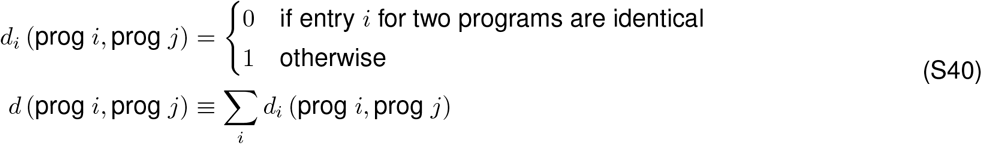

Note that this permutation includes flipping the policy (*a*_±_ → *a*_∓_ doesn’t change the performance due to the symmetry of the task) —i.e., a program with flipped policy is considered to be the same program.

###### When two programs have different sizes

With merger programs at hand (see the merger rule-out rule in Section 7.4 for details), it is now easy to define structural-distance for two different-size programs as well. Consider two programs with size *M* and *M* + Δ*M*, one can simply

1. list all the valid merger programs equivalent to *M* with size *M* + Δ*M*, and
2. find the one merger program that would give the minimal structural-distance from another program with size *M* + Δ*M*

###### Verifying the structural distance between two programs

In Figure S9, we show how program 24 and program 7655 are separated by only a single mutation, despite the fact that they appear to be described by very different graph representations. This example highlights the set of the operational steps that we used in computing structural distance: 1) finding a merger program, 2) permuting nodes, 3) flipping all actions, and 4) mutating.

##### 9.2 Tree Embedding (TE)

The main goal of TE is not to capture all relational structures from a program space, but to capture the minimal relations that could help us to make sense of the structures hidden in such a space. Having this in mind, it make sense to use tree for it contains minimal number of edges (number of edges equals number of nodes) among all classes of graphs. In this study, we also keep smaller programs closer to the root. The reason of doing so, because the more states a program has the more functionally diverse it could be. Doing TE this way therefore potentially captures such a growing functional diversity as one traversing outwards from the root.

**Figure S8:**
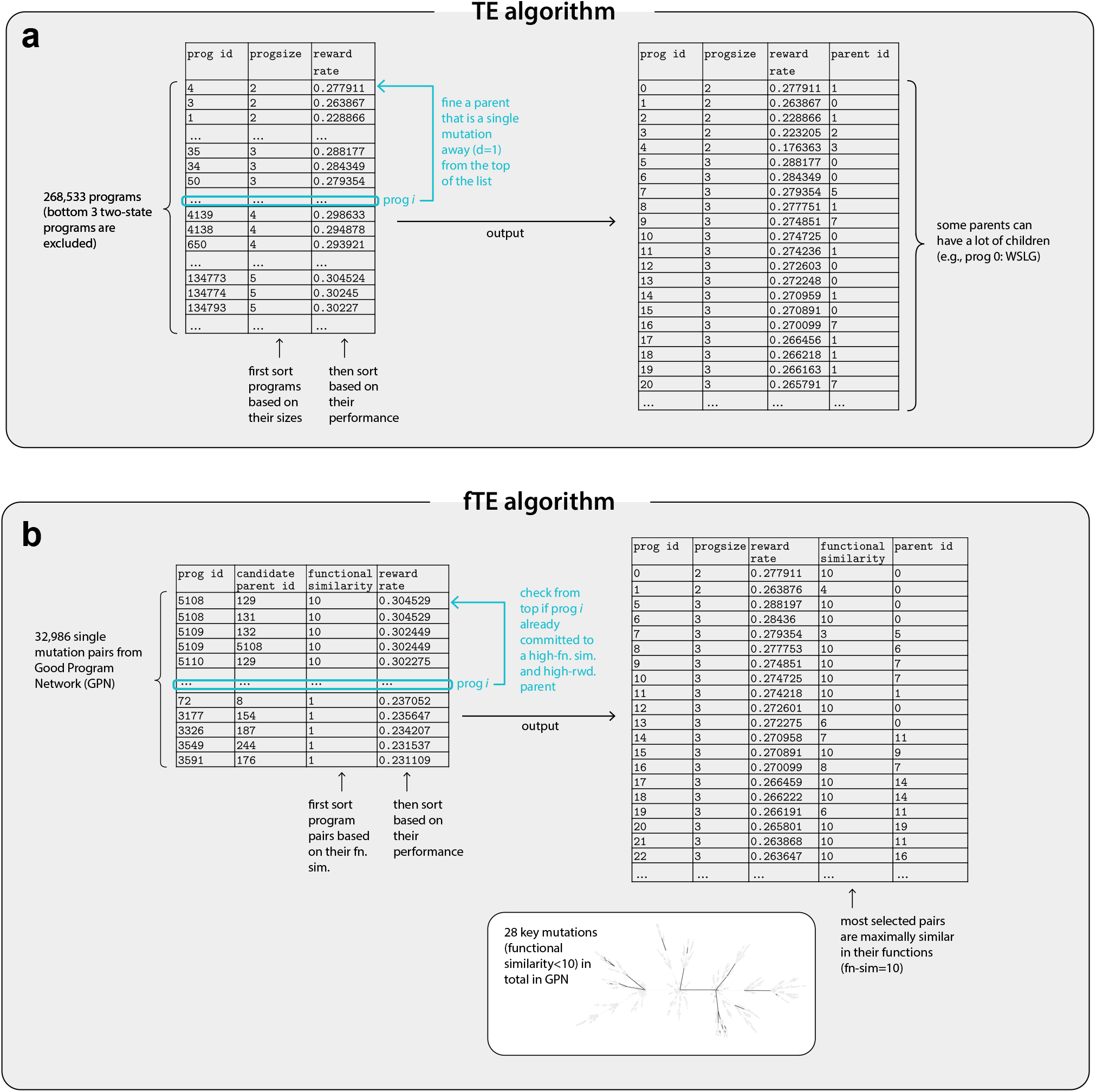
Tree Embedding (TE) & functional Tree Embedding (fTE) algorithm. **(a)** TE consists of two major algorithmic steps: 1) sort a full list of program according to their individual attributes, and 2) find a single mutation parent from the top of the list. **(b)** In contrast, fTE starts from sorting a list of single mutation pairs with parent size no bigger than the children. The sort prioritizes functional similarity before performance. Insert: the resultant tree shows that all key mutations amount to a tiny fraction (28 out of 4492) of Good Program Network, indicating fTE successfully found a smooth embedding in program functions.

###### Two algorithmic steps in TE

In Figure S8a, we illustrate the two steps for TE algorithm:

*Step I: Sort programs* according to 1) *program size*, small to large and 2) *performance*, high to low
*Step II: Find one parent with single mutation* for a program *i* from the top of the list.

**Figure S9:**
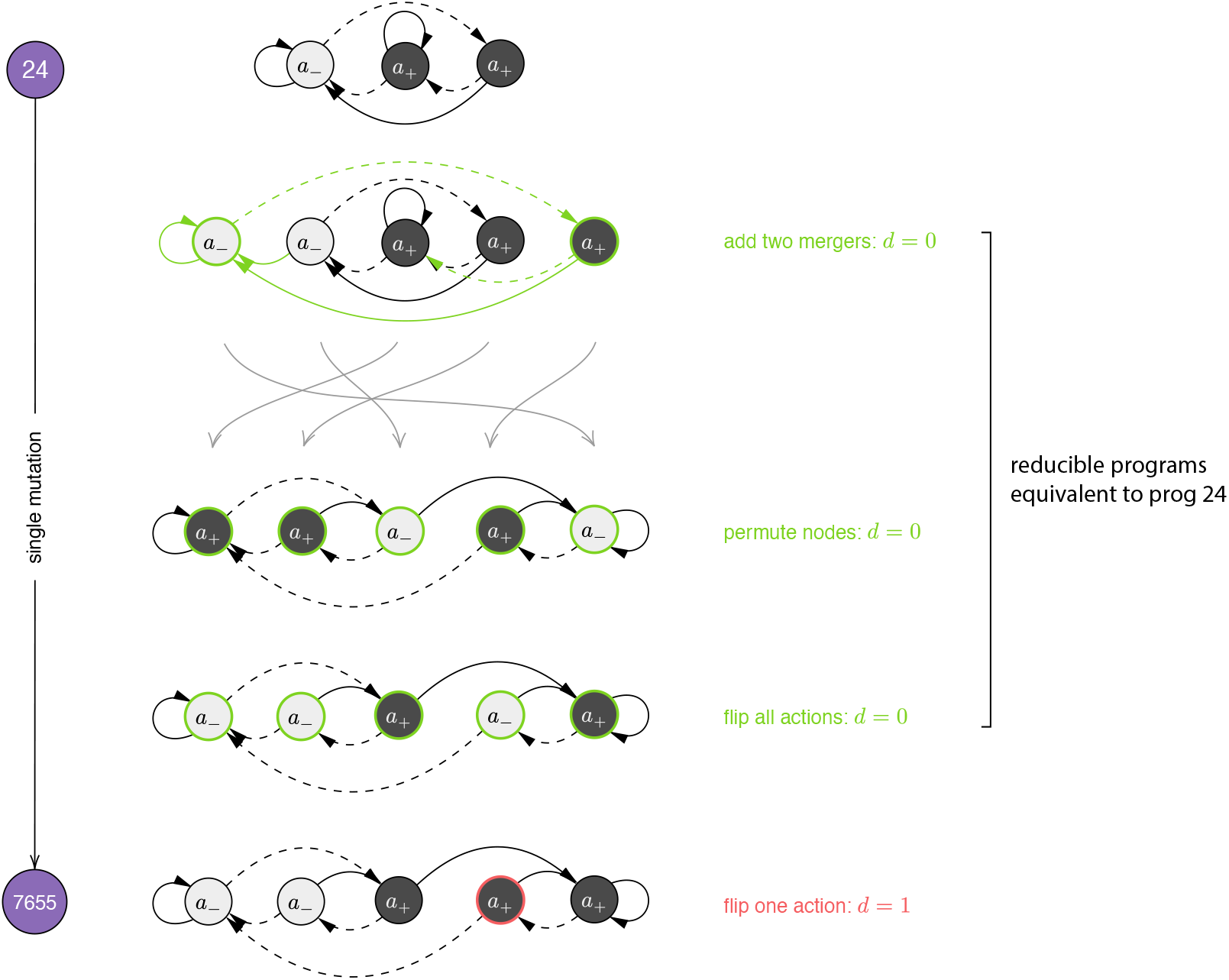
Algorithmic steps in verifying structural distance. In this example, prog 7655 is only a single mutation away from prog 24, despite the apparent difference in their wiring. Note that either 1) growing a program by adding mergers, 2) permuting nodes, or 3) flipping all actions creates an equivalent program with identical behaviors.

The first step of TE is simply to sort the programs according to their size and performance. This way, one makes sure that all five unique two-state programs are root nodes of the tree and most of the larger programs are closer to the leaf nodes. In the second step, one loop through the list of all programs (program *i* in Figure S8a) and search for the first program *j* on the same list that is only a single mutation away. Since this program list is sorted from small to large for program size and from high to low performance, it preferentially connect a program *j* to a program relatively small in size and high in performance.

##### 9.3 Randomized Tree Embedding (TE) predicts robustness of evolutionary algorithms

In the standard TE above, we discover that nearly all good programs are closely connected through single mutations (see Figure 3 in the main text and Figure S12), and vice versa. One might questions whether this observation is simply an artifact of the secondary sorting based on their performance. To address this issue, we randomize the performance in the list and carry out the embedding. The result is shown in Figure S10. As one can see that the conclusion is largely unchanged, and the program space is indeed highly structured in the relations of their performance. This robustness test with randomized TE thus predicts the success of evolutionary algorithm for one can start from a descent program and easily finds many more good programs that are just one mutation away. The result from the evolutionary algorithm is discussed next.

**Figure S10:**
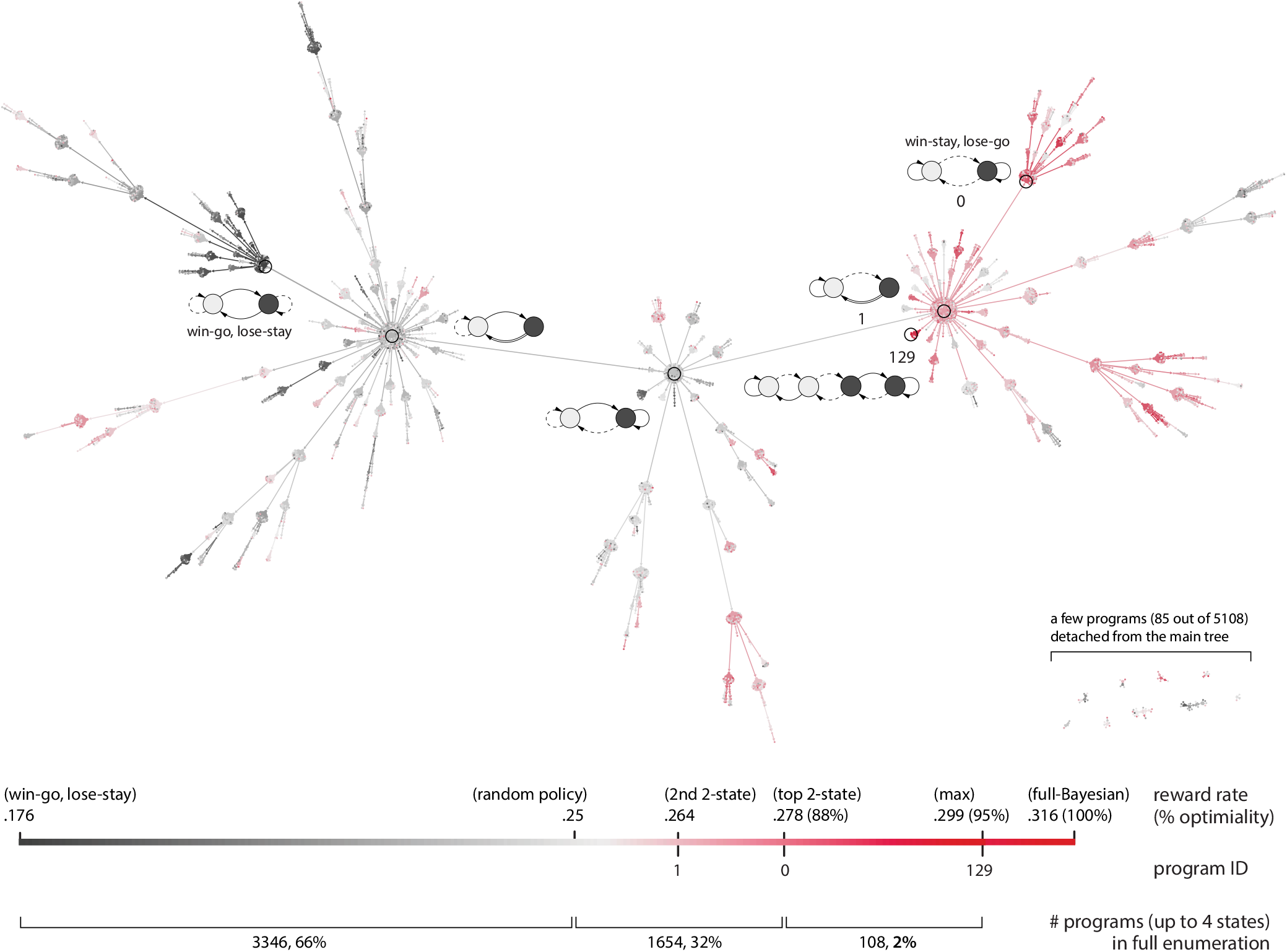
Reward-randomized Tree Embedding. In this particular tree embedding, we no longer sort program according to their performance; that is, a program no longer biased toward selecting a high-performance parent. The fact that nearly all good programs are still closely connected to form a single sub-tree implies that the full program space is indeed highly structured (rather than an artifact of our embedding algorithm).

##### 9.4 Efficient search via an evolutionary algorithm

The main features of our version of evolutionary algorithm are that:

1. we consider all possible mutations at each generation rather than doing random mutations;
2. we use a selection criterion that is not a fixed performance threshold;
3. we hibernate unselected mutants with a chance to be mutated later.

It seems that implementing the above features requires specifying many algorithmic rules. Yet surprisingly, this can be done with a relatively simple scheme with four distinct operations as illustrated in Figure S11a and steps below:

###### program_reservoir

The local enumeration operates on a single data frame called program reservoir (PR). PR is a spread sheet with each row stores all relevant attributes for a unique program. Below is an example:

To grow this list, one may start from a small set of valid programs, and use the following four operations in sequence at each epoch. Each operation takes in individual programs on the current list and update program reservoir according to some specific rules. Although these rules can be arbitrarily complex based on all the program attributes, we aim for finding the most simple and yet effective rules.

| prog id | parent prog id | prog size | reward rate | t_born | partition_born | t_morph | partition_morph | status |
| --- | --- | --- | --- | --- | --- | --- | --- | --- |
| 3 | None | 2 | .263867 | 0 | None | [1] | [(1,1), (2,1)] | morph |
| 4 | None | 2 | .277911 | 0 | None | [ ] | [ ] | idle |
| ... | ... | ... | ... | ... | ... | ... | ... | ... |

###### morph_op

morph_op closely resembles the random mutation step in an evolutionary algorithm. The only difference is that in our case, we enumerate every possible mutation without any randomness or sub-sampling involves. Below is the pseudocode:

1. Grab all programs with status labeled “`morph`.”
2. Assign “`t_morph`” as current epoch to the selected programs.
3. Assign two simplest partitions, i.e., `partition_morph=[(1, 1, …, 1), (2, 1, …, 1)]`, to individual programs for program size *M*. (note that first partition is for generating mutants without changing program size, while the second one finds mutants with size *M* + 1).
4. Based on `partition_morph`, generate mutants with appropriate merger programs (these mutants are all the possible 1-mutations).
5. Filter invalid mutants with the standard rule-out rules (see Table S10 and Table S11 for details).
6. Output a list of mutants each with attributes: “`t_born`” and “`partition_born`”
7. Change all the status of “`morph`” to “`done`.”

###### append_op

The function of append op is only to check whether a new mutant is not in program reservoir yet. Here one simply need to reuse the following two rule-out rules designed for the full enumeration:

1. Remove identical graphs under node inversion.
2. Remove identical graphs under node permutation.
3. This last step is just to assign status to newly appended programs “`idle`.“

###### eval_op

In the evaluation operation, we simply find the corresponding reward rate for an un-evaluated program from the database generated from the full enumeration. But for *M* ≥ 6 where such a database doesn’t exist, one can run BDP (see Section 8.1) to acquire reward rate.

###### select_op

The selection operation is a crucial step in our local enumeration. Depending on the number and reward rates of the selected programs, the algorithm will explore different part of the program space with different rates. But the bottom line for an effective selection rule is to find maximal fraction of good programs with minimal exploration in program space. Below is a simplest selection rule to achieve both objectives at the same time.

1. Grab all programs with status “`idle`.”
2. Compute `num_morph = log(num_idle) * fac_morph`
3. Assign new status from “`idle`” to “`morph`” for `num_morph` top performing programs, and leave the rest “`idle`.”

This specific selection rule has a few advantages as mentioned above:

1. It is equivalent to flexibly adjusting the reward threshold for selecting good programs. There could be a particular epoch where new mutants doesn’t perform well. And since the rule demands to still select `~log(num_idle)` programs, such a local barrier wouldn’t hinder the local enumeration.
2. By using selecting `~log(num_idle)` many programs, the rule bends an exponentially growing enumeration to become a polynomially growing one. This allows one to let program evolves for a long time (e.g., a hundred epochs)
3. Such a selection rule never discard a program in program reservoir. Rather, it hibernates them as “idle” programs. And when a certain epoch doesn’t generate much good programs, some of the top hibernating programs can be woken up and ready to be mutated.

**Figure S11:**
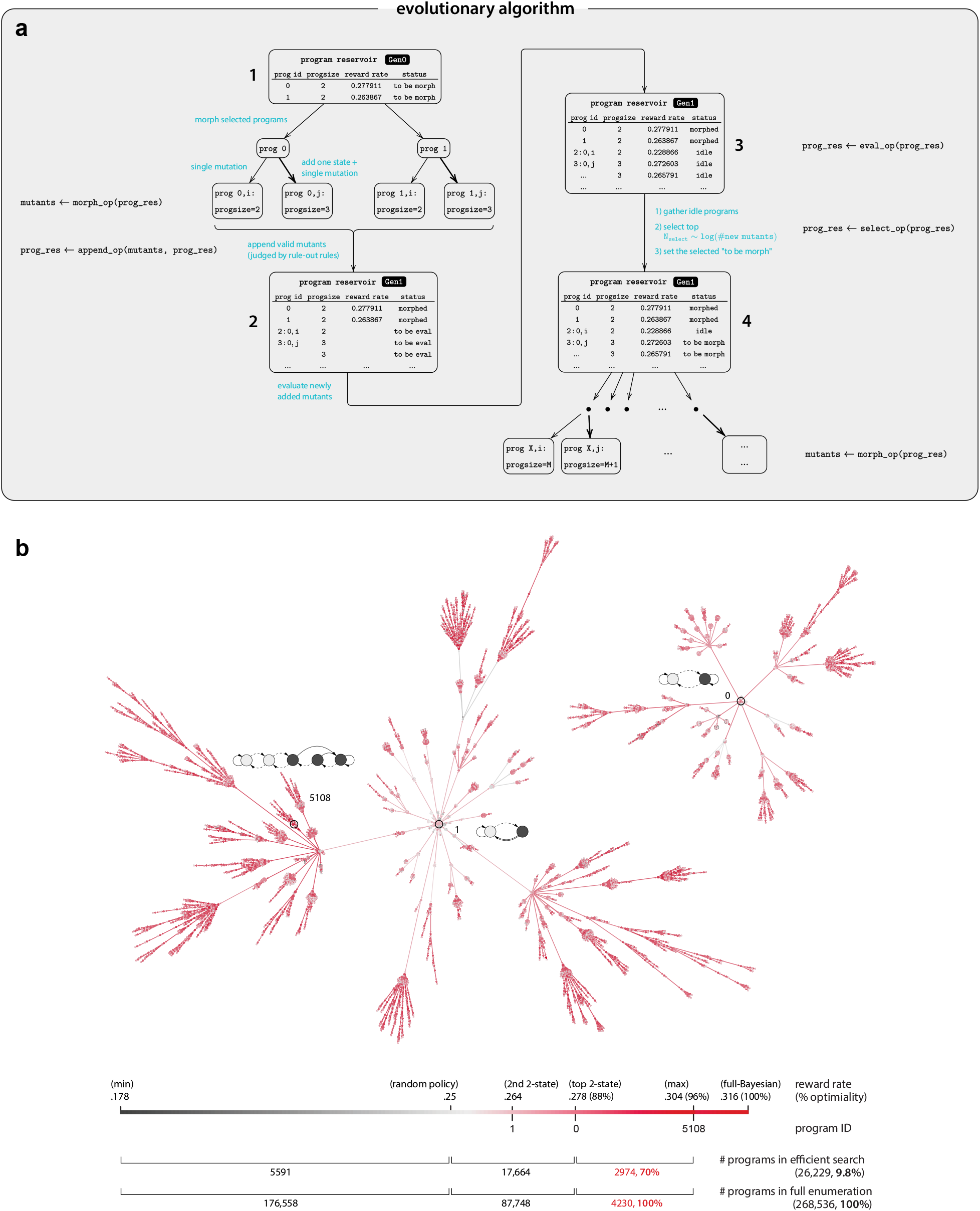
Evolutionary algorithm for an efficient search. **(a)** Algorithmic steps for the evolutionary algorithm. For each generation, a set of new mutants are validated, evaluated, and selected for the next generation. The algorithm adopts a fixable selection criterion such that not only the very top performing program generates off-springs. Note that we evaluate all possible mutants without any sub-sampling. **(b)** Visualization of how the full program space is explored after a search of 16 generations. The algorithm explore only 10% of the full space while acquiring 70% of all good programs. Note that the larger tree corresponds to a root of a lower performance program, indicating that it is beneficial to keep slightly lower performance ones during the search.

###### Results

From the table in Figure 3f in the main text and Figure S11b, one can see that the evolutionary algorithm efficiently find majority of good programs with a search that is only a fraction of the full program space. Interestingly in Figure S11b, one can see that the larger tree is actually rooted in the second highest performing two-state program instead of WSLG program. This observation also highlights that there is a reason to keep relatively lower performing programs in the process of evolution as well.

##### 9.5 Visualizing the full program space (*M* ≤ 5) with a reduced Tree Embedding (rTE)

In Figure 3 in the main text, we visualize a smaller program space (*M* ⩽ 4, 5108 programs) directly from TE with Gephi[23] effortlessly. However, visualizing the full space with *M* ⩾ 5 with much more than 10,000 nodes (268,536 nodes to be exact) is not doable in Gephi. Additional ingenuity is therefore needed for reducing the full size tree. One way to do this is to group neighboring programs based on their local and global relational structures.

###### Local relational structure

As discussed in Section 9.1, two nearby programs with the same or different size can be categorize by a partition of their corresponding merger program; e.g.,

| size change upon mutation | all possible partition type |
| --- | --- |
| $M = 3 \rightarrow 3$ | (1, 1, 1) |
| $M = 3 \rightarrow 4$ | (1, 1, 2) |
| $M = 2 \rightarrow 4$ | (1, 3), (2, 2) |

The partition type is therefore a way to measure whether two descendants are structurally similar—i.e., two descendant programs are similar if they share the same merger program, or the same partition after mutated from their parent.

###### Global relational structures

We use two global relational structures for rTE. The first one is called “*branch*” in which we trace backwards from an individual program to the root. The second one is called “*eccentricity*” in which we trace forwards from an individual program to the farthest leaf node.

###### branch

As discussed above, every program in TE has a certain partition except the root. The first step in rTE is to find the sequence of partitions, or the branch, that goes from the root to a target program. To do so, one simply follow the immediate parent of the target program, and find the parent of the parent, etc., until reaching the root. Next, one can simply replace this chain of program ids with their corresponding partitions:

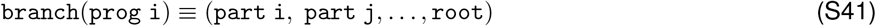

###### eccentricity

Computing eccentricity (i.e., traversing length from the current node to the farthest reachable node) for a program is straightforward, one just need to take previous branches (but with program ids instead of partitions) and search the longest segment. For example, for program 42

1. Grab all branches that contain `42`: `[11, 22, 33, 42, 56, 78]` `[11, 22, 33, 42, 77, 99, 111]` `[11, 22, 33, 42, 77, 88]`
2. Get the segments towards the leaf node: `[56, 78]` `[77, 99, 111]` `[77, 88]`
3. Find the maximal length: `eccentricity(42) = 3`

**Figure S12:**
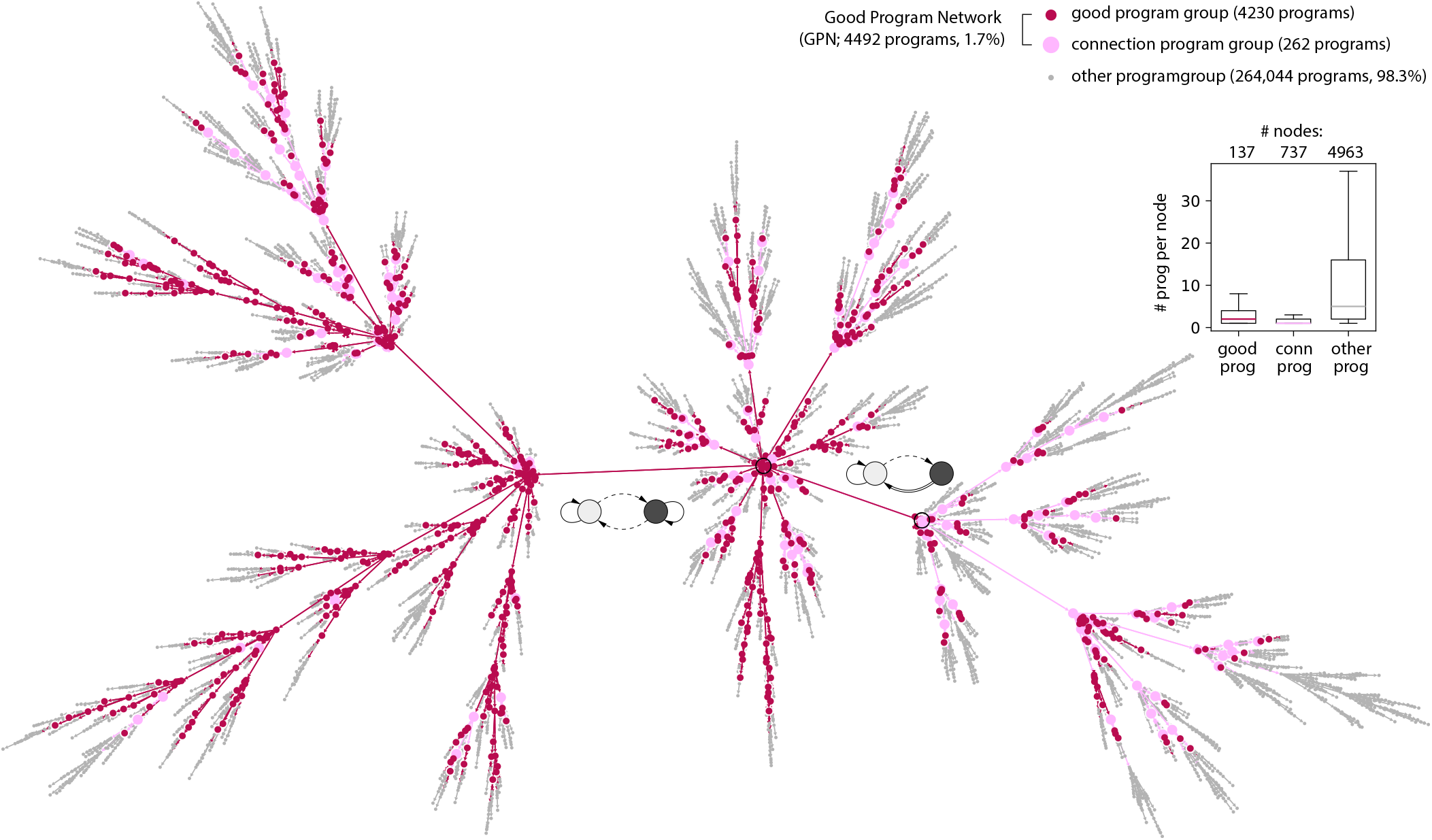
Visualizing the full program space via reduced Tree Embedding (rTE). Visualizing TE with 268,533 nodes (programs) in Gephi is challenging. We therefore use rTE to group nodes that are similar in their relational structure on the tree. Specifically, a group of programs are merged if both 1) their position relative to the root node (`branch`) are identical and 2) their position relative to the leaf node (`eccentricity`) are identical. Note that we keep the connection programs (with performance lower than WSLG program) which glue multiple separate lineages together into a single component Good Program Network (GPN)

###### Grouping programs with identical relational tuples

`(branch, eccentricity)` Now that each program has a new label: `(branch, eccentricity)`, one can group those programs with the identical tuple together as a compound node. The parent of such a compound node would be another compound node without the leaf partition and with a strictly larger eccentricity. Finally, with all compound nodes and their parents assigned, we have a reduced tree ready to be visualized. The result is shown in Figure S12 with the reduced number of nodes: 5837 from the original 268,533 nodes.

##### 9.6 Extracting the Good Program Network (GPN) from a Tree Embedding (TE)

From the tree embedding (TE) discussed earlier, one notes that nearly all good programs are closely connected to form a single component sub-tree. One can easily see this is indeed the case in Figure S12 in which the full program space is visualized in rTE discussed above. The extracted Good Program Network (GPN) consists of 4492 programs within which all programs except the 262 connection programs have the performance higher or equal to WSLG program. Note that in this visualization, GPN seems to take over most of the program space, but this is a false perception due to this particular way of visualization given that only 1.7% of the full space is GPN.

It’s worth noting that we decided to keep those 262 (6% of GPN) connection programs because we would like to study the relational structure among all 4230 good programs. And having these connection programs holding every detached branches together allows one to study how these programs are connected though some evolutionary lineages to the full extent.

#### 10 Extracting functional structures from the Good Program Network (GPN)

In this section, we discuss in detail how functional Tree Embedding (fTE) is performed, and all the necessary mathematics used in the embedding. Moreover, we derive the notion of key and sloppiness mutations—that are used to understand how GPN programs diversify—from these mathematics. Lastly, we show detailed algorithmic steps for Motif Decomposition (MD) and discuss how it can be used to understand whether a group of program can be locally or globally identified.

##### 10.1 *confusion matrix, confusability*, and *identifiability*

Both confusability and identifiability measures how a program can be identified among a group of other programs based on its behavioral sequences. Formally it can be defined as

**Table S13:**
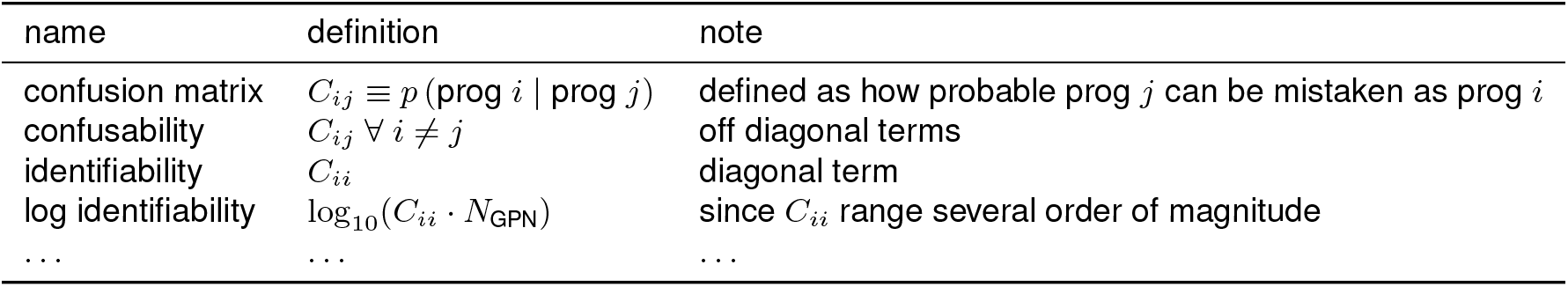
definition of confusion matrix and identifiability.

| name | definition | note |
| --- | --- | --- |
| confusion matrix | $C_{ij} \equiv p(\text{prog } i \mid \text{prog } j)$ | defined as how probable prog $j$ can be mistaken as prog $i$ |
| confusability | $C_{ij} \forall i \neq j$ | off diagonal terms |
| identifiability | $C_{ii}$ | diagonal term |
| log identifiability | $\log_{10}(C_{ii} \cdot N_{\text{GPN}})$ | since $C_{ii}$ range several order of magnitude |
| ... | ... | ... |

With the above conceptualized, the rest is to define the confusion matrix:

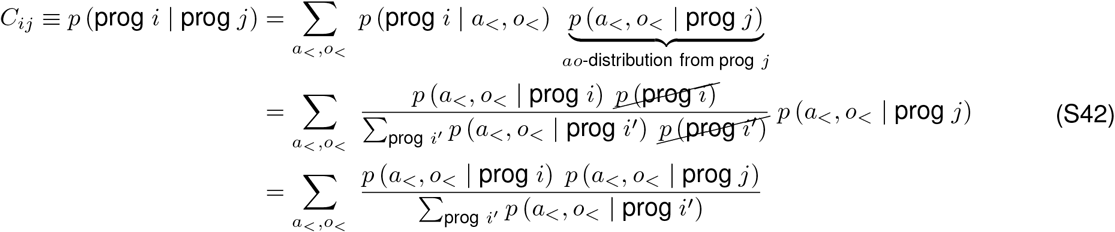

In the above derivation, we assume that all programs within the group have the same probability whereas all programs outside the group have zero probability—i.e.,

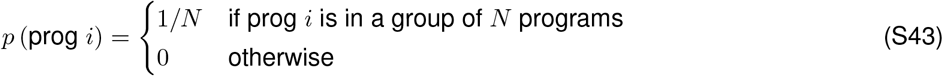

##### 10.2 structural distance matrix

Previously in TE, we only computed structural distance when checking if a candidate parent has distance *d* = 1 (see definition of distance in Section 9.1). Here we will do the same computation but for all pair of programs within GPN: In fTE, one needs search a parent among all single-mutation programs, computing this pairwise distance matrix is therefore necessary.

**Table S14:** definition of structural distance matrix.

| name | definition | note |
| --- | --- | --- |
| *structural distance matrix* | $d_{ij} \equiv d(\text{prog } i, \text{prog } j)$ | |
| ... | ... | ... |

##### 10.3 functional similarity matrix

**Table S15:** definition of functional similarity matrix.

| name | definition | note |
| --- | --- | --- |
| *functional similarity matrix* | $F_{ij} \equiv \sum_l F_{ij}^l$ | $F_{ij}^l$ is defined below as with a fixed sequence length $l$ |

The fixed length functional similarity matrix can be defined as follow:

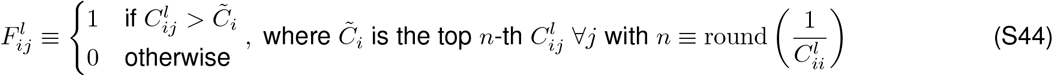

Intuitively, given a program *i*, the inverse identifiability tells us how many program *j* would be confused with it; i.e., 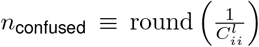. These confused programs then takes a binary value 1: 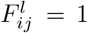. Some programs are functionally close such that they cannot be differentiated even with long sequences, whereas some programs are functionally distinct that can be easily distinguished even with short sequences. In our case, that translate into a range from 1 to 10 in total functional similarity:

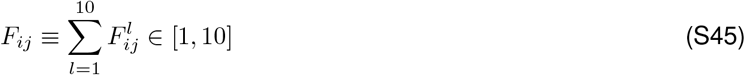

##### 10.4 A functional Tree Embedding (fTE) of the Good Program Network (GPN)

One major goal of fTE is to ensure that the embedding space is as functionally smooth as possible (more discussion in the main text); that is, there should be as many neighbors (*i, j*) with their functional similarity *F_ij_* = 10 as possible. With all necessary mathematics, we are now ready to formalize fTE algorithm as illustrated in Figure S8b:

1. From structural distance matrix *d_ij_*, append a list of program pair (prog *i*, prog *j*) with *d_ij_* = 1 and *M_i_* ⩽ *M_j_* (the size of parent program *j* is not larger).
2. Append the corresponding functional similarity from *F_ij_* for each selected program pair.
3. Append the corresponding reward rate < *R* > for each program *i* in selected pairs.
4. Firstly, sort this program pair list based on *F_ij_*, and secondly sort it based on < *R* >.
5. Loop from the top of the list, check if prog *i* is already in the final list; if not, append the row into the final list.

As one can see in Figure S8b with first 20 programs, fTE successfully find a functionally smooth embedding in which most program pairs are functionally similar *F_ij_* = 10 with only 28 (out of 4492) pairs has *F_ij_* < 10.

##### 10.5 Key and sloppy mutations

From fTE, we see two emerging notions: key and sloppy mutations that can now simply defined as

It’s important to stress that the notion of key and sloppy mutations should not be taken as a binary classification but, more appropriately, a continuum. In this study, we only use this binary naming to provide a more intuitive discussion.

**Table S16:** definition of key and sloppy mutations.

| name | definition |
| --- | --- |
| *key mutation* | $\text{prog } j \rightarrow \text{prog } i \text{ with } F_{ij} < l_{\max}$ |
| *sloppy mutation* | $\text{prog } j \rightarrow \text{prog } i \text{ with } F_{ij} = l_{\max}$ |

##### 10.6 Functional labels of programs in the Good Program Network (GPN)

fTE provides an insightful relational structure, a skeleton, to hold all good programs together through some branching evolutionary lineages. In the main text (Figure 4), we categorize these programs into 4 distinct classes based on their function and wiring.

###### Structurally Bayesians programs (DBs)

Structural Bayesian is defined simply as discretized Bayesians (DBs) in which their wiring display a clear bi-directional belief integration (see Section 5.3 for details in constructing and enumerating DBs)

###### Functionally Bayesians programs (fDBs)

As the name suggests, a functional Bayesian (fDB) are functionally indistinguishable from structural Bayesians (DBs). To define fDB, we first compute how much an arbitrary program *i* in GPN confuses with all 65 existing DBs, or how much program *i* can be identified among them:

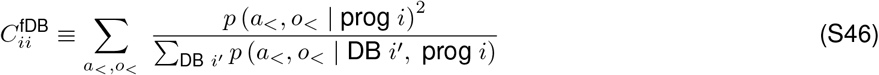

The full derivation of the above is shown in Equation (S42). Next we need to pick a threshold 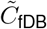 in which

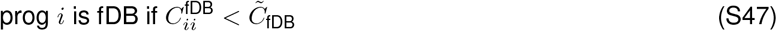

It turns out that this threshold can be defined unambiguously based on the scatter plot in Figure 4b in which there are only two outlier DBs if we use 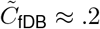.

###### Functionally identifiable programs

From Equation (S42), we know that program *i* has its identifiability among all GPN programs defined as

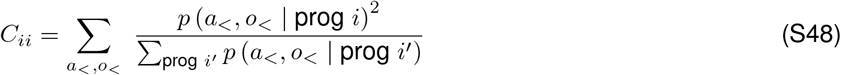

To define an identifiable programs in a parameter-free fashion, we use “halving principle of classification” in which top *n* identifiable programs add up half of the global identifiability:

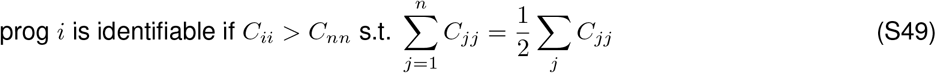

Note that in the above index *j* is sorted so that *C*_11_ > *C*_22_ > … > *C*_jj_ > … > *C_nn_*.

###### Other unidentifiable programs

Unidentifiable program is defined simply as the opposite from the above:

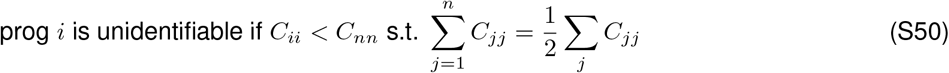

###### Grouping functionally identifiable programs

In Figure 4, we further cluster all functionally identifiable programs using `community_louvain`. In their corresponding confusion matrix, we can see 9 nearly functionally orthogonal program groups. Though this clustering result gives one a sense of functional diversity among good programs, it is important to stress (as discussed in Section 1.6) that these programs within a group should not be seen as homogeneous, and those individuality could matter in some other context.

##### 10.7 Motif Decomposition (MD)

In Section 3, we discussed in depth why MD is crucial in understanding the functional diversity in Good Program Network (GPN), and how it could provide a powerful way to explore a larger program space that a more complex task may demand. Below, we show the algorithmic steps for MD in detail.

###### Step 1) extract identifiable sequences for a given program

The reason that we do MD with only identifiable sequences because these sequences represent the identity of this particular program among all other programs in GPN. The extraction of identifiable sequences in a program *i* is similar to extracting identifiable program which we discussed in Equation (S49):

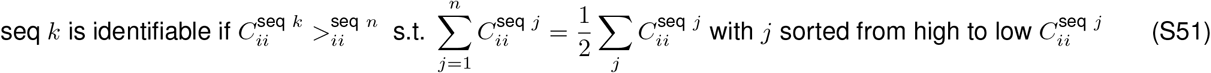

where

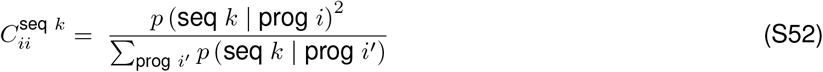

###### Step 2) use identifiable sequences to discover motifs as action-outcome loops with length l_motif_ ⩽ *l*_max_ = 10 from identifiable sequences

Given a set of identifiable sequences, one can convert them into “win-lose-and-stay-go” sequences. For example:

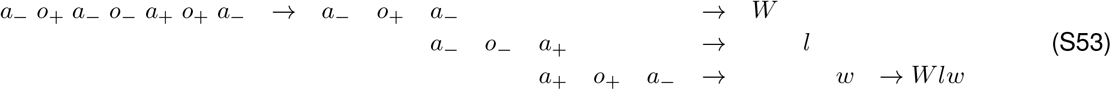

where

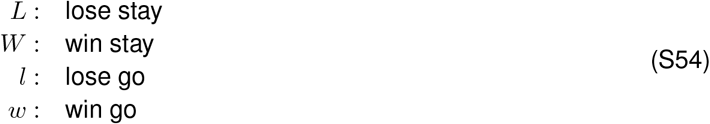

Next, we enumerate all possible candidate loops, and check if all possible long sequences a given candidate loop can generate exist among all nonzero sequences. For example:

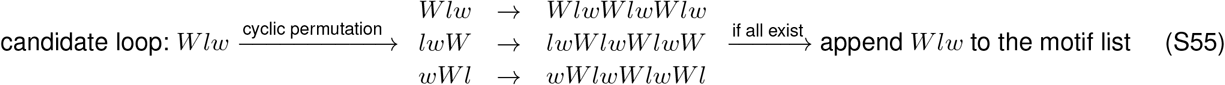

###### Step 3) decompose all nonzero sequences in terms of extracted motifs

In this final step, we use a soft decomposition approach aiming to use a set of motifs (or motif compound) to explain a given sequence with the following features:

1. prioritizing using shorter motifs
2. without double counting the contribution of each motifs
3. avoiding algorithm with hyperparameters (no hyperparameter means no need to fine tune to get a “better” result)

Surprisingly, there is a relatively simple way to implement the above. As an example, consider decomposing the following sequence of prog 24 in GPN:

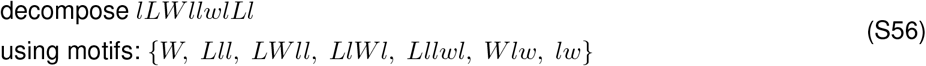

We first start from sorting these motifs based on length

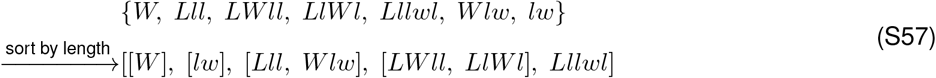

Then for a given length, check how much each motif explain the remaining unexplained part of the sequence the most:

**Table S17:**
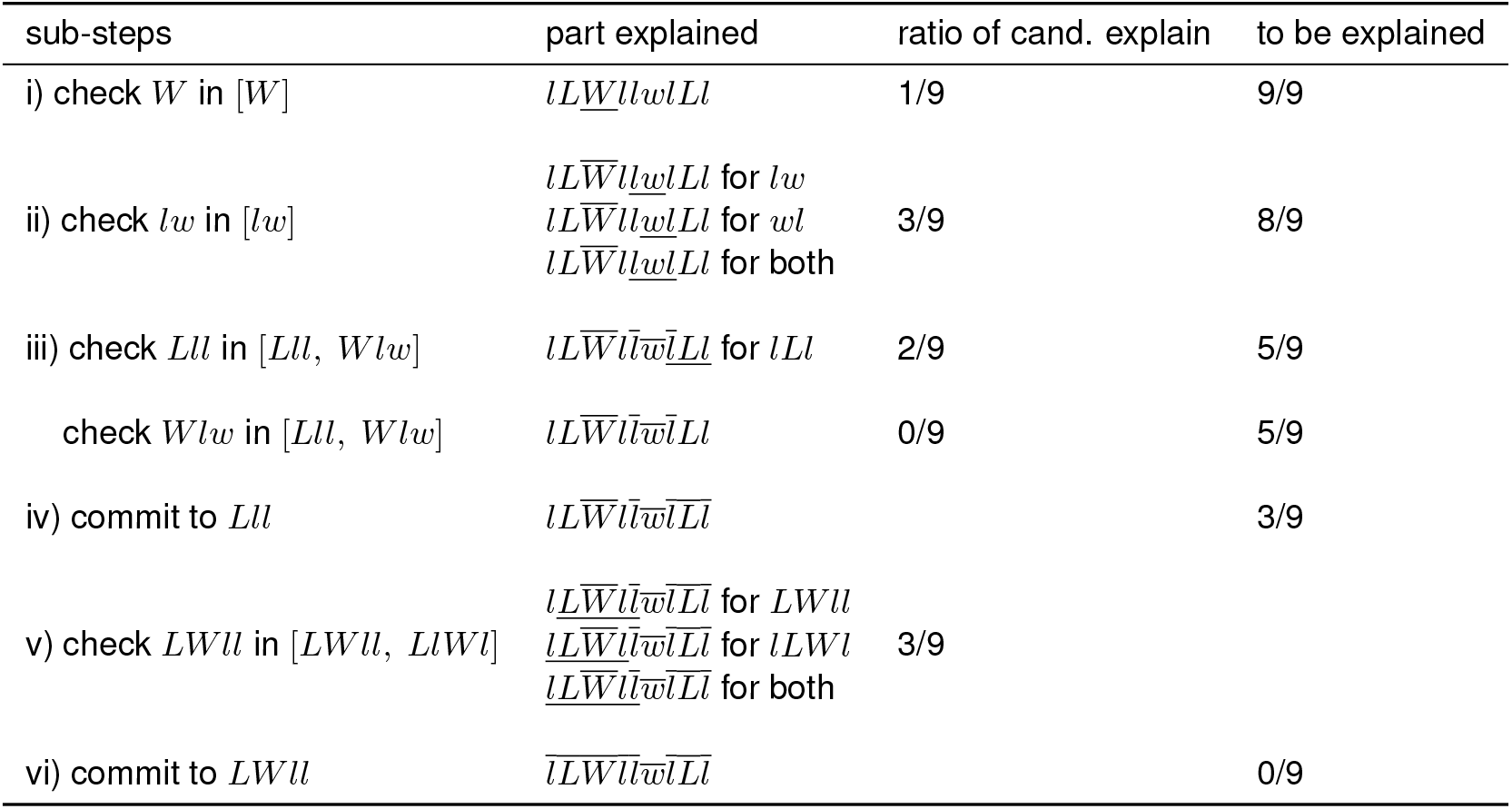
Algorithmic steps of Motif Decomposition (MD).

where the underline indicates the candidate explanation, whereas the overline indicates the finalized explanation. The last step where *LWll* is committed without checking *LlWl* is because all the remaining parts are explained, so it is not possible for *LlWl* to top it. The final result of this particular example is therefore:

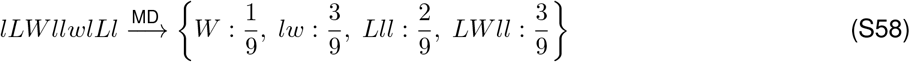

##### 10.8 Locally identifiable programs can be globally unidentifiable

In Figure 5 in the main text, we focus on a local clusters of descendant of program 9 and 24. In doing so, we gain insights regarding the notion of key and sloppy mutations. There, we defined key and sloppy mutation globally Table S16. However, one can easily think of a situation where having a local notion of key and sloppy mutations could be useful. For example, after only a few steps in an evolutionary algorithm, not much of the good program space is explored, and thus global information about distant part of GPN is not available.

In Figure S13a, we show that many descendants of program 9 surpass the criterion to be functionally distinct locally, and thus being classified as undergoing a key mutation. Within those, three program stand out as the most identifiable ones: program 24, 6592, and 8261. Interestingly, only program 24 remains to be identifiable globally, whereas the other two programs becomes part of a sea of functional Bayesians. In Figure S13e, we show two other functional Bayesians: program 6118 and 5982 that generates overlapping behavioral sequences.

**Figure S13:**
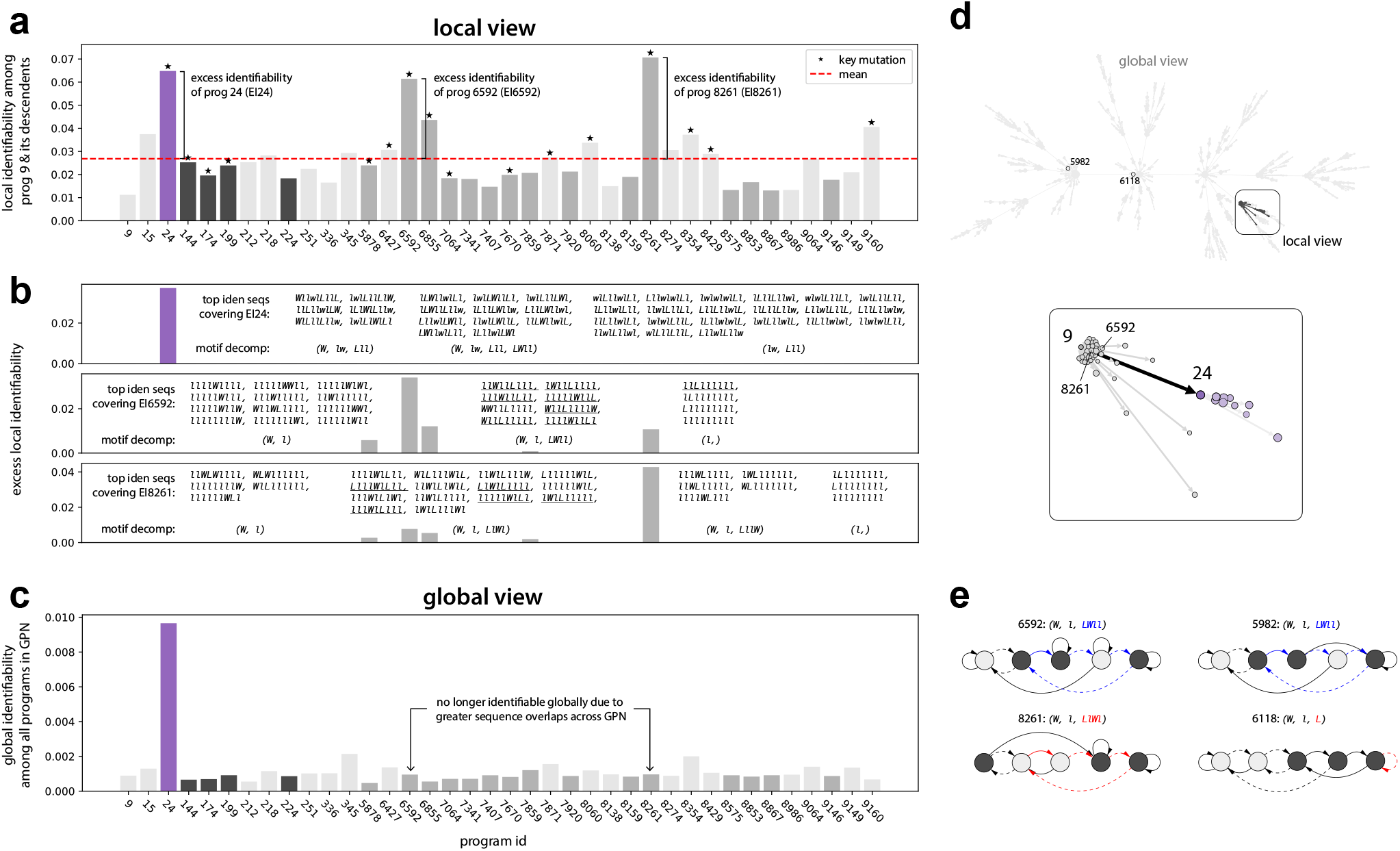
Local vs global notations of key mutation and sloppy mutation. **(a)** Many locally identifiabiliable programs emerge from local key mutations. Among those, three programs (prog 24, 6592, and 8261) stand out as the most functionally unique. **(b)** These three programs are most identifiable for they generate many unique sequences that other programs do not share. To confirm this, we re-plot (a) but only keeping those sequences that most contribute the identifiability of the corresponding programs; e.g., for prog 24 (purple), those sequences are listed in the insert. The fact that all other programs lost their identifiability shows these behavioral sequences can only be generated by prog 24. **(c)** only prog 24 remains to be globally identifiable, while prog 6592 and 8261 become a part of the functional-Bayesian group. **(d)** Local clusters of the descendants of prog 9 is highlighted in Good Program Network (GPN) embedded with fTE. **(e)** Two distant programs: prog 5982 and 6118 are used as examples to illustrate that many programs in GPN have similar behaviors. Specifically, the sequences underlies in middle panel of (b) are shared between prog 6592 and 5982; similarly those in the bottom panel in (b) are shared between prog 8261 and 6118. Note that these shared sequences can be realized by the same or different combination of motifs (discussed in the main text).

#### 11 ProgEnum (all codes)

In this section, we provide basic instructions for generating all of the results in this paper. We encourage readers to modify our code to fit their own needs, for example to develop novel enumeration schemes, evolutionary algorithms, or more versatile tree embeddings. We look forward to hearing from you about how these ideas might be useful in different problem domains.

For more detailed information and documentation, please visit our github repository https://github.com/HermundstadLab/ProgEnum

##### 11.1 Begin to explore

As a starting point, we provide three main datasets that were generated from sequentially running the main codes. Install the lastest version of pandas, and use `example_code_to_begin.py` to load the following dataframes:

**Table S18:**
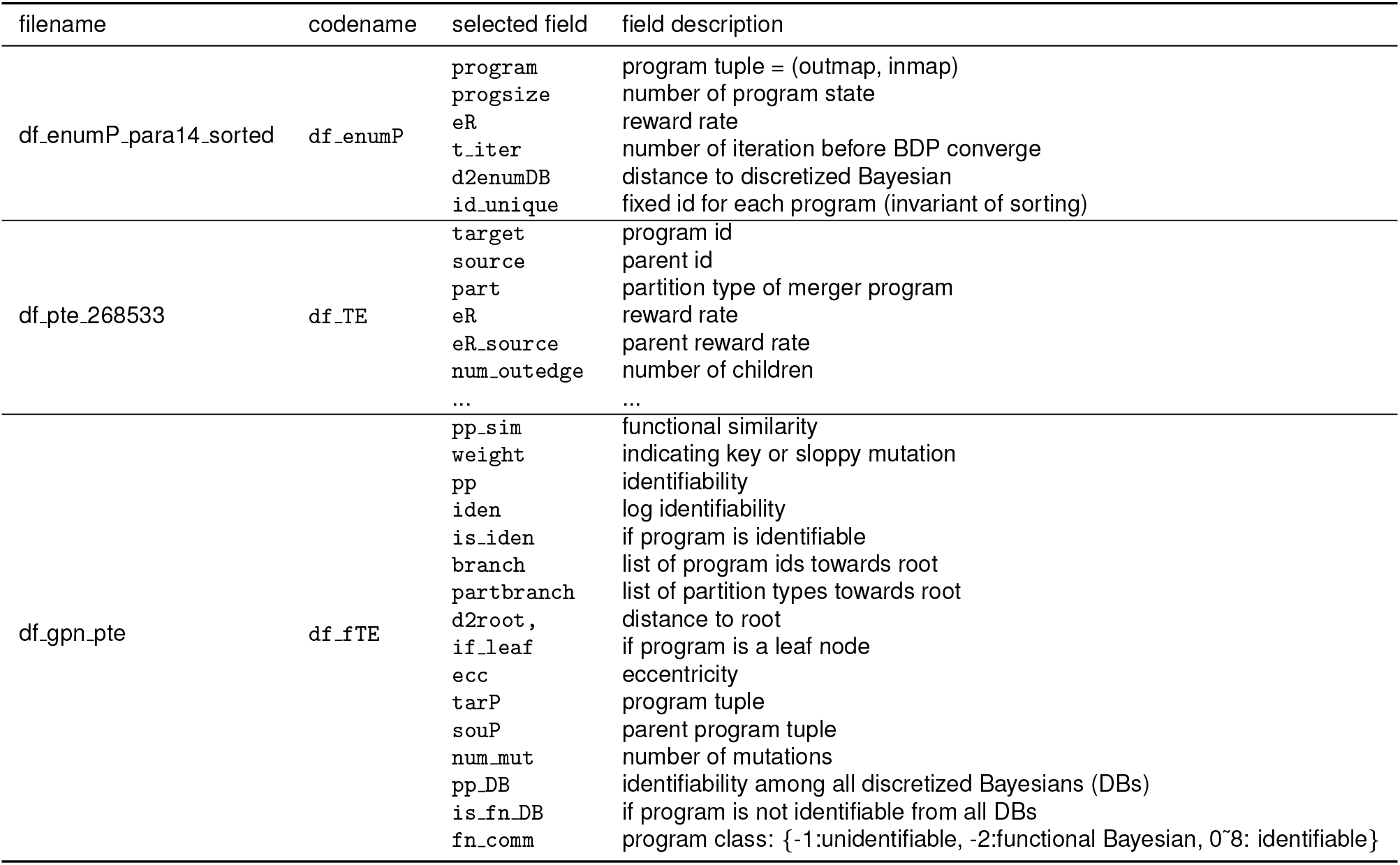
Selected datasets. Run `example_code_to_begin.py` to start exploring.

##### 11.2 Full usage

Below we provide a minimal instruction for running our code (visit our github repository for full documentation):

1. Run all `main_.py` sequentially to generate all necessary datasets.
2. Run `fig_.py` or `print_.py` separately to create plots or display data.

A minimal description for running each code is listed in the tables below:

**Table S19:** Main codes. These codes have to be run sequentially.

| filename | algorithm | note | resource | run time |
| --- | --- | --- | --- | --- |
| main_0_df_task.py |  | generate df_task | laptop |  |
| main_1a_df_task_evalB.py | BDP_B | append reward rates of opt-B to df_task | laptop |  |
| main_1b_df_task_genDB.py |  | append list of DBs to df_task | laptop | 11m |
| main_1c_df_task_evalDB.py | BDP | append reward rates of DBs to df_task | cluster, 96 cpus | 25m |
| main_2a_df_enumDB.py | DB, enumDB | generate df_enumDB_para14 | laptop | 12m |
| main_2b_df_enumDB_eval.py | BDP | append reward rates of enumDBs to df_enumDB_para14 | cluster, 96 cpus | 13m |
| main_3a_df_enumP.py | rule-out rules | generate df_enumP | laptop | 43m |
| main_3b_df_enumP_eval.py | BDP | append reward rates of small programs to df_enumP_para14 | cluster, 96 cpus | 1.8h |
| main_3c_df_enumP_d2enumDB.py | structural distance | append minimal distance to DBs to df_enumP_para14 | laptop | 24m |
| main_3d_df_enumP_sorted_for_pte.py | TE | generate df_enumP_para14_sorted, df_enumP_para14_rand for TE and reward-randomized TE. | laptop |  |
| main_3e_df_enumP_tsne.py | TSNE | generate df_enumP_para14_tsne | cluster, 96 cpus | 2h |
| main_4a_pte.py | TE | generate df_pte_268533 for gephi visualization | cluster, 96 cpus | 5h |
| main_4b_rpte.py | rTE | generate df_rpte_268533 | laptop | 50m |
| main_4c_rpte2gpn.py | GPN | generate df_gpn | laptop |  |
| main_5_lenumP.py | evolutionary algorithm | generate df_lenumP5_toplog2 | laptop | 15m |
| main_6a_gpn_sdp.py | SDP | generate df_gpn_Y_lists | laptop | 30m |
| main_6b_gpn_pprog.py | confusion matrix | generate df_gpn_pprog_arr | cluster, 96 cpus × 45 jobs | 10m |
| main_6c_gpn_simprog.py | functional similarity | generate df_gpn_simprog_arr | laptop |  |
| main_6d_gpn_dprog.py | distance matrix | generate df_gpn_dprog_arr | cluster, 96 cpus × 45 jobs | 5m |
| main_6e_gpn_pte.py | TE | generate df_gpn_pte | laptop |  |
| main_6f_gpn_fnDB.py | functional Bayesian | append identifiability among DBs to df_gpn_pte | laptop |  |
| main_6g_gpn_fnComm.py | functional labels | append functional labels to df_gpn_pte | laptop | 5m |
| main_6h_df_gpn_local_keymut.py | local key mutation | generate df_gpn_local_keymut | laptop |  |
| main_6i_gpn_seq2motif.py | MD | generate df_gpn_seq_motif_0, df_gpn_seq_motif_1, and df_gpn_seq_motif_2 | cluster, 96 cpus | 30m |
| main_7a_df_enumTaskB.py | enumerate hazard rate | generate df_enumTaskB_special_h | laptop | 5m |
| main_7b_df_enumTaskDB.py | enumerate hazard rate; two types of DB | generate df_enumTaskDB_data | cluster, 96 cpus | 3h |
| main_7c_df_enumTaskP.py | enumerate hazard rate; BDP on extended GPN | generate df_enumTaskP_data | cluster, 96 cpus × 68 jobs | 5m |

**Table S20:** Codes for figures and displaying data. These codes can be run independently.

| filename | figures, tables, boxes |
| --- | --- |
| fig_bar_keymut & iden_local vs global.py | Figure S13 |
| fig_bar_local keymut.py | Figure S13 |
| fig_bar_mut leading to fn sim vs dissim.py | Figure 4 |
| fig_bar_novel motifs inherited.py | Figure 5 |
| fig_fineDB.py | Figure 2 |
| fig_imshow_fnComm.py | Figure 4 |
| fig_scatter_fnDB.py | Figure 4 |
| figS_boxplot_rTE.py | Figure S12 |
| figS_imshow_why TE.py | Figure S1 |
| figS_plot_why motif.py | Figure S5 |
| figS_scatter_enumTaskB_DB.py | Figure S7 |
| figS_scatter_enumTaskP_highDimSloppiness.py | Figure S4 |
| figS_scatter_enumTaskP_seqVenn.py | Figure S7 |
| figS_scatter_enumTaskP.py | Figure S7 |
| figS_scatter_tsne.py | Figure S3 |
| figS_table_lenumP.py | Figure 3 |
| print_critical h.py | Table S7 |
| print_descendants of prog 9 and 24.py | Figure 5, Figure S13 |
| print_extended good program space for enumTaskP.py | Box S4 |

**Table S21:** Source codes.

| filename | note |
| --- | --- |
| core_bdp.py | BDP for evaluating small program |
| core_bdpB.py | BDP for evaluating optimal Bayesian |
| core_enum.py | enumerating program |
| core_enumDB.py | discretizing and enumerating Bayesian |
| core_pte.py | all functions required for TE |
| core_sdp.py | SDP for small program |
| core_sdpB.py | SDP for optimal Bayesian |

## References

[1] P. R. Montague, P. Dayan, C. Person, and T. J. Sejnowski, “Bee foraging in uncertain environments using predictive hebbian learning,” Nature, vol. 377, no. 6551, pp. 725–728, 1995.

[2] M. Vergassola, E. Villermaux, and B. I. Shraiman, “‘infotaxis’ as a strategy for searching without gradients,” Nature, vol. 445, no. 7126, pp. 406–409, 2007.

[3] S. D. Boie, E. G. Connor, M. McHugh, K. I. Nagel, G. B. Ermentrout, J. P. Crimaldi, and J. D. Victor, “Information-theoretic analysis of realistic odor plumes: What cues are useful for determining location?,” PLoS computational biology, vol. 14, no. 7, p. e1006275, 2018.

[4] E. Fujioka, I. Aihara, M. Sumiya, K. Aihara, and S. Hiryu, “Echolocating bats use future-target information for optimal foraging,” Proceedings of the National Academy of Sciences, vol. 113, no. 17, pp. 4848–4852, 2016.

[5] S. B. M. Yoo, J. C. Tu, S. T. Piantadosi, and B. Y. Hayden, “The neural basis of predictive pursuit,” Nature neuroscience, vol. 23, no. 2, pp. 252–259, 2020.

[6] M. M. Botvinick, “Hierarchical models of behavior and prefrontal function,” Trends in cognitive sciences, vol. 12, no. 5, pp. 201–208, 2008.

[7] P. Shamash, S. F. Olesen, P. Iordanidou, D. Campagner, N. Banerjee, and T. Branco, “Mice learn multi-step routes by memorizing subgoal locations,” Nature Neuroscience, vol. 24, no. 9, pp. 1270–1279, 2021.

[8] A. Loisy and C. Eloy, “Searching for a source without gradients: how good is infotaxis and how to beat it,” Proceedings of the Royal Society A, vol. 478, no. 2262, p. 20220118, 2022.

[9] J.-J. O. de Xivry, S. Coppe, G. Blohm, and P. Lefevre, “Kalman filtering naturally accounts for visually guided and predictive smooth pursuit dynamics,” Journal of neuroscience, vol. 33, no. 44, pp. 17301–17313, 2013.

[10] A. Solway, C. Diuk, N. Córdova, D. Yee, A. G. Barto, Y. Niv, and M. M. Botvinick, “Optimal behavioral hierarchy,” PLoS computational biology, vol. 10, no. 8, p. e1003779, 2014.

[11] J. O’Doherty, M. L. Kringelbach, E. T. Rolls, J. Hornak, and C. Andrews, “Abstract reward and punishment representations in the human orbitofrontal cortex,” Nature neuroscience, vol. 4, no. 1, pp. 95–102, 2001.

[12] L. P. Sugrue, G. S. Corrado, and W. T. Newsome, “Matching behavior and the representation of value in the parietal cortex,” science, vol. 304, no. 5678, pp. 1782–1787, 2004.

[13] M. P. Karlsson, D. G. Tervo, and A. Y. Karpova, “Network resets in medial prefrontal cortex mark the onset of behavioral uncertainty,” Science, vol. 338, no. 6103, pp. 135–139, 2012.

[14] B. A. Bari, C. D. Grossman, E. E. Lubin, A. E. Rajagopalan, J. I. Cressy, and J. Y. Cohen, “Stable representations of decision variables for flexible behavior,” Neuron, vol. 103, no. 5, pp. 922–933.e7, 2019.

[15] A. Rajagopalan, R. Darshan, J. E. Fitzgerald, and G. C. Turner, “Expectation-based learning rules underlie dynamic foraging in drosophila,” bioRxiv, 2022.

[16] C. C. Beron, S. Q. Neufeld, S. W. Linderman, and B. L. Sabatini, “Mice exhibit stochastic and efficient action switching during probabilistic decision making,” Proceedings of the National Academy of Sciences, vol. 119, no. 15, p. e2113961119, 2022.

[17] R. Bellman, “A problem in the sequential design of experiments,” Sankhyā: The Indian Journal of Statistics (1933-1960), vol. 16, no. 3/4, pp. 221–229, 1956.

[18] V. D. Costa, V. L. Tran, J. Turchi, and B. B. Averbeck, “Reversal learning and dopamine: a bayesian perspective,” Journal of Neuroscience, vol. 35, no. 6, pp. 2407–2416, 2015.

[19] P. Vertechi, E. Lottem, D. Sarra, B. Godinho, I. Treves, T. Quendera, M. N. Oude Lohuis, and Z. F. Mainen, “Inference-based decisions in a hidden state foraging task: Differential contributions of prefrontal cortical areas,” Neuron, vol. 106, no. 1, pp. 166–176.e6, 2020.

[20] Y. Liu, Y. Xin, and N.-l. Xu, “A cortical circuit mechanism for structural knowledge-based flexible sensorimotor decision-making,” Neuron, vol. 109, no. 12, pp. 2009–2024, 2021.

[21] J. H. Sul, H. Kim, N. Huh, D. Lee, and M. W. Jung, “Distinct roles of rodent orbitofrontal and medial prefrontal cortex in decision making,” Neuron, vol. 66, no. 3, pp. 449–460, 2010.

[22] M. L. Tsetlin et al., Automaton theory and modeling of biological systems, vol. 102. Academic Press New York, 1973.

[23] M. Bastian, S. Heymann, and M. Jacomy, “Gephi: an open source software for exploring and manipulating networks,” in Third international AAAI conference on weblogs and social media, 2009.

[24] Y. Hu, “Efficient, high-quality force-directed graph drawing,” Mathematica journal, vol. 10, no. 1, pp. 37–71, 2005.

[25] R. L. Rivest and R. E. Schapire, “Diversity-based inference of finite automata,” Journal of the ACM (JACM), vol. 41, no. 3, pp. 555–589, 1994.

[26] T. Aynaud, “python-louvain x.y: Louvain algorithm for community detection.” https://github.com/taynaud/python-louvain, 2020.

[27] E. C. Tolman, “Cognitive maps in rats and men.,” Psychological review, vol. 55, no. 4, p. 189, 1948.

[28] T. E. Behrens, T. H. Muller, J. C. Whittington, S. Mark, A. B. Baram, K. L. Stachenfeld, and Z. Kurth-Nelson, “What is a cognitive map? organizing knowledge for flexible behavior,” Neuron, vol. 100, no. 2, pp. 490–509, 2018.

[29] W. F. Młynarski and A. M. Hermundstad, “Adaptive coding for dynamic sensory inference,” Elife, vol. 7, p. e32055, 2018.

[30] M. Zacksenhouse, R. Bogacz, and P. Holmes, “Robust versus optimal strategies for two-alternative forced choice tasks,” Journal of mathematical psychology, vol. 54, no. 2, pp. 230–246, 2010.

[31] F. Lieder and T. L. Griffiths, “Resource-rational analysis: Understanding human cognition as the optimal use of limited computational resources,” Behavioral and brain sciences, vol. 43, 2020.

[32] J. Pearl, Heuristics: intelligent search strategies for computer problem solving. Addison-Wesley Longman Publishing Co., Inc., 1984.

[33] G. Gigerenzer and W. Gaissmaier, “Heuristic decision making,” Annual review of psychology, vol. 62, no. 1, pp. 451–482, 2011.

[34] G. Gigerenzer and R. Selten, Bounded rationality: The adaptive toolbox. MIT press, 2002.

[35] K. J. Miller, M. M. Botvinick, and C. D. Brody, “From predictive models to cognitive models: Separable behavioral processes underlying reward learning in the rat,” bioRxiv, p. 461129, 2021.

[36] P. Krueger, F. Callaway, S. Gul, T. Griffiths, and F. Lieder, “Discovering rational heuristics for risky choice,” PsyArXiv Preprints, Jan. 2022.

[37] W. Młynarski, M. Hledík, T. R. Sokolowski, and G. Tkačik, “Statistical analysis and optimality of neural systems,” Neuron, vol. 109, no. 7, pp. 1227–1241, 2021.

[38] K. S. Brown and J. P. Sethna, “Statistical mechanical approaches to models with many poorly known parameters,” Physical review E, vol. 68, no. 2, p. 021904, 2003.

[39] A. A. Prinz, D. Bucher, and E. Marder, “Similar network activity from disparate circuit parameters,” Nature neuroscience, vol. 7, no. 12, pp. 1345–1352, 2004.

[40] B. C. Daniels, Y.-J. Chen, J. P. Sethna, R. N. Gutenkunst, and C. R. Myers, “Sloppiness, robustness, and evolvability in systems biology,” Current opinion in biotechnology, vol. 19, no. 4, pp. 389–395, 2008.

[41] G. R. Yang, M. R. Joglekar, H. F. Song, W. T. Newsome, and X.-J. Wang, “Task representations in neural networks trained to perform many cognitive tasks,” Nature neuroscience, vol. 22, no. 2, pp. 297–306, 2019.

[42] J. M. Whitacre, “Biological robustness: paradigms, mechanisms, and systems principles,” Frontiers in genetics, vol. 3, p. 67, 2012.

[43] R. S. Sutton and A. G. Barto, Reinforcement learning: An introduction. MIT press, 2018.

[44] I. Wolfram Research, “Mathematica, version 12.3.” https://www.wolfram.com/mathematica, 2022.

[45] QuantEcon, “Quantecon: A high performance open source python code library for economics.” https://github.com/QuantEcon/QuantEcon.py, 2021.

